# MAP4K inhibition by the CNS penetrant inhibitor famlasertib restrains medulloblastoma dissemination without developmental toxicity

**DOI:** 10.64898/2026.05.11.724200

**Authors:** Marc Thomas Schönholzer, Meng-Syuan Lin, Shen Yan, Veronica Akle, Amin Allalou, Bernard Ciraulo, Dejana Versamento, Levi Luca Kopp, Dina Hochuli, Tanja Bleiker, Annik Welti, Stephan C.F. Neuhauss, Martin Baumgartner

## Abstract

Cerebellum tissue invasion and dissemination are major drivers of recurrence and metastatic spread in medulloblastoma (MB), yet no invasion-inhibitory therapy is currently available. The serine/threonine kinase MAP4K4 is highly expressed in MB and promotes invasive behavior downstream of growth factor signaling, while its physiological postnatal downregulation suggests that disrupted developmental control may contribute to tumor pathogenesis. Here, we investigate pharmacological targeting of MAP4K4 using famlasertib, a CNS-penetrant, neuroprotective MAP4K inhibitor, as a strategy to suppress invasion in MB.

Using 3D invasion assays and quantitative live-cell microscopy, we demonstrate that famlasertib markedly reduces invasive behavior and single-cell motility of MB cells. Organotypic cerebellum tissue and orthotopically implanted zebrafish larval models further confirm anti-invasive efficacy without detectable developmental toxicity at effective concentrations. Phospho-proteomics and functional analyses revealed that MAP4K inhibition alters kinase signaling linked to cytoskeletal remodeling, leading to suppressed F-actin dynamics, increased cell clustering, and enhanced cortical accumulation of tight junction protein 1 (TJP1).

Collectively, our findings confirm MAP4Ks as therapeutically targetable regulators of MB invasion and establish famlasertib as a candidate migrastatic agent that restrains tumor cell dissemination while preserving developmental integrity.

## Introduction

Brain and nervous system tumors rank as the second most prevalent type of cancer affecting children globally, with medulloblastoma (MB) as the most frequent malignant tumor within this category ^1^. MB has been molecularly classified into four subgroups (Wingless (WNT), Sonic Hedgehog (SHH), Group 3, and Group 4, encompassing a total of 12 subtypes ^2,3^. Current treatment approaches consist of surgical resection, cytotoxic chemotherapy, and radiotherapy ^4^, yet long-lasting side effects linked to tumor growth and treatment are common ^5–7^, highlighting the need of non-toxic therapies to increase survivors’ quality of life.

The dysregulation of the molecular processes that control neural development is associated with tumorigenesis in MB ^8–10^, and the hijacking of neurodevelopmental epigenetic programs promotes metastatic dissemination ^11^. Several receptor tyrosine kinases (RTK) including PDGFR and ERBB2, their downstream effectors of the MAPK pathway as well as PI3K-AKT signaling components were implicated as motility drivers in various MB *in vitro* and *in vivo* models (reviewed in ^12^). A pro-migratory mediator downstream of RTK activation is the Ser/Thr kinase mitogen-activated kinase kinase kinase kinase 4 (MAP4K4) ^13^, which is an actin cytoskeleton regulatory kinase upregulated in MB ^14^ that promotes invasiveness via PKCθ activation and VASP S157 phosphorylation ^15^. Although migration-promoting factors in MB are known ^16^, effective migrastatic therapies ^17^ and evidence of their clinical benefit in MB are still lacking. Cytoskeleton-targeting drugs show potential but are limited by nonspecific toxicity, highlighting how toxicity towards normal cells remains a major challenge for cytoskeleton-targeting anti-cancer drugs ^18^. Poor CNS tolerance also obstructed further development of the MAP4K4 inhibitor **C29** ^19^ for CNS malignancies and prompted the subsequent development of GNE489 to restrict blood-brain barrier (BBB) penetration ^20^. More recently, a screen for agents that protect neurons from undergoing ER-stress-induced apoptosis in the context of neurodegeneration identified the MAP4K inhibitor prostetin12k ^21^. Prostetin12k has the international nonproprietary name famlasertib, and a phase I clinical trial of famlasertib (prosetin; NCT05279755) is currently underway to evaluate its safety and pharmacokinetics in healthy volunteers and in patients with amyotrophic lateral sclerosis (ALS). Notably, famlasertib crosses the blood–brain barrier and does not exhibit overt toxicity in mice ^21^.

The potent MAP4K inhibitory activity of famlasertib, together with its reported ability to cross the blood– brain barrier and its neuroprotective properties, prompted us to investigate whether it can suppress invasion-promoting mechanisms in MB. Using microscopy-based 3D invasion and 2D migration assays, we quantitatively assessed the effects of famlasertib and characterized compound-induced phenotypes associated with reduced cell invasion and motility. Organotypic tissue and zebrafish larval and models were employed to evaluate toxicity and to test anti-invasive activity in physiologically relevant contexts. Phosphoproteomic profiling uncovered molecular targets of famlasertib in tumor cells and identified the F-actin regulator paxillin and the cell–cell adhesion protein TJP1 as novel candidate effectors of MAP4K signaling.

## Material and Methods

### Cell culture and cell lines

DAOY (male, 4Y), D283 (male, 6Y, ATCC, USA), and HEK-293T cells were cultured in IMEM or DMEM (Sigma), respectively. UW228 (female, 9Y), ONS-76 (female, 2Y), HD-MB03 (male, 3Y), and D425 (male, 6Y) cells were kindly provided by John Silber (USA), Michael Taylor (Canada), Till Milde (Germany), and Henry Friedman (UK), and cultured in DMEM or RPMI-1640 (Sigma). WT and TJP1 KO MDCK-II cells were provided by Alf Honigmann (Germany), and cultured in MEM. MB cell line identities were confirmed by SNP typing (Multiplexion GmbH, Germany). Media were supplemented with 10% FBS, 1% Penicillin-Streptomycin (ThermoFisher Scientific), and 1% GlutaMAX (Gibco), and cells were maintained at 37 °C with 5% CO₂. UW228 LA-EGFP and ONS-76 LA-EGFP lines were generated by lentiviral transduction with pLenti-LA-EGFP. Mycoplasma contamination was excluded using LookOut® PCR (#MP0035, Sigma Aldrich) and MycoAlert (Lonza, Visp, Switzerland).

### Spheroid invasion assay (SIA)

SIA was performed and quantified according to ^22^. Briefly, 2500 cells/100 µl were seeded in cell-repellent 96-well plates (Corning) and incubated 24 h to form spheroids. On day 2, 70 µl medium was replaced with 70 µl of 2.7% collagen I (CellSystems). Hydrogels were overlaid with 100 µl serum-free medium containing 2× growth factors/inhibitors. Cells invaded the matrix for 24 h, were stained with Hoechst 33342 (1:2000), and imaged at 5× using an Operetta system (Revvity, Inc., Waltham, MA, USA). Invasion distance from spheroid borders was quantified with Harmony 4.9 software.

### Single cell motility analysis (nuclear tracking)

Live-cell imaging of nuclear fluorescence was performed on a Nikon Ti2 inverted microscope using 10×/0.3 NA or 20×/0.75 NA objectives with GFP and Cy3 filters. Cells were imaged for 12 h with frames every 3–6 min; excitation was applied only during acquisition, with no detectable bleaching. Before tracking, the movies were downsized from 2424x2424 pixels by a factor of 3 or 4, to 808x808 or 606x606 pixels, respectively, using local pixel average mean. Contrast was enhanced by limiting and redistributing the dynamic range to 8-bit. The global background intensity was estimated as the 50th percentile of the intensity distribution across the whole-time sequence. Then the intensity values were coerced to 1 to 3-times the background value, normalized to 0–255 range and converted to 8-bit. Nuclei tracking was performed by scripting the Fiji plugin TrackMate ^23^. Nuclei were detected via Laplacian of Gaussian (radius 25-40 µm, threshold 0.5–1.5), and trajectories linked with SparseLAPTracker (LINKING_MAX_DISTANCE 20–50 µm; GAP_CLOSING, SPLITTING, MERGING disabled). Mother and daughter cells were tracked separately. Only traces ≥60 min with non-zero displacement and mean speed were used for analysis and plotting.

### Organotypic cerebellum slice culture and TJP1 quantification in the tissue context

OCSC cultures were authorized by the Cantonal Veterinary Service of Zürich (ZH079/23) and performed as described in ^24^. Briefly, cerebellar slices were prepared from P9-P11 wild-type C57BL/6JRj mice, and cultured on Millipore insert (PICM 03050, Merck) for 12-14 days with brain slice medium^25^ before co-culture with tumor spheroid. A maximum of three slices were placed per insert, and media were changed daily for two weeks. Tumor spheroids were generated by seeding ONS-76 LA-EGFP or D425 cells in low-attachment, round-bottom 96-well plates (4520, Corning, USA) at a density of 5000 cells per well and culture for 48 hours. One spheroid was then placed per slice, and the tumor spheroid-slice co-cultures were treated for five days with famlasertib or tilfrinib or an equivalent volume of solvent (DMSO).

After permeabilizing and blocking, anti-ZO-1 (D6L1E) antibody (1:200, 13663, Cell Signaling), Anti-calbindin antibody [EP3478] (1:1000, ab108404, Abcam), Anti-Glial Fibrillary Acidic Protein (GFAP) antibody (1:200, ab53554, Abcam), Anti-Nuclei Antibody [clone 3E1.3] (1:200, MAB4383, Merck) were used for immunofluorescence analysis. The inserts containing slices were flat mounted onto glass slides in VECTASHIELD® antifade mounting medium (H-1000-10, Vector Laboratories). Images were acquired on a SP8 Leica confocal microscope (Leica Microsystems, Mannheim, Germany) and spinning disk confocal microscope (Nikon CrestOptics X-Light V3, Japan).

### Spheroid growth assay

1500 ONS-76 cells per well were seeded in 96-well, round bottom Ultra Low Attachment Plate **(#**7007, Corning). After 48 h, the cells were treated with a range of famlasertib concentrations and corresponding DMSO (vehicle controls) dilutions. Image stacks over the entire growth clusters were acquired using a 5x air objective with an Operetta automated microscope (Revvity, Inc., Waltham, MA, USA) 24, 48, 72 and 96 h after seeding (0, 24 and 48 h after compound treatments). One image per stack in optimal focus was used for segmentation as well as for area and roundness analysis using Harmony 4.9 software. 12 wells were analyzed per condition.

### RNA interference

MDCK-II cells at approximately 60% confluency were transfected with Dharmafect 4 Transfection Reagent (T-2005-01, Horizon) and 10 nM siRNAs following the manufacturer’s instruction. siRNAs used are listed in Supplementary Table 8. Non-targeting control siRNA was used as a negative control. After 48 h, cells were fixed and processed for immunofluorescence analysis.

### Immunoblotting (IB)

Cells were lysed in RIPA buffer with protease (cOmplete™, Roche) and phosphatase (PhosSTOP™, Roche) inhibitors and cleared by centrifugation. Protein concentration was measured using the Pierce™ BCA assay (Thermo Fisher). Proteins were separated on 4–20% Mini-PROTEAN® TGX gels (Bio-Rad) and transferred to 0.2 µm nitrocellulose membranes. After 1 h blocking with 5% milk, membranes were incubated with primary antibodies overnight at 4 °C, followed by HRP-linked secondary antibodies and detection with IB Substrate or SuperSignal™ West Femto (Thermo Fisher). Band intensity was quantified using Image Lab 5.2.1 and ImageJ/Fiji.

### Real-time PCR /qRT-PCR

Total RNA was isolated using the RNeasy Mini Kit (Qiagen). cDNA was synthesized from 1 µg RNA with the High-Capacity cDNA Reverse Transcription Kit (Applied Biosystems). qRT-PCR was performed using TaqMan® Gene Expression Master Mix on a QuantStudio 7 Pro system. Predesigned TaqMan® assays (Thermo Fisher) were used (see supplementary Table 8). Gene expression was calculated using the 2^−ΔCt method normalized to GAPDH.

### Immunofluorescence analysis (IFA)

7×10³ wild-type or LA-EGFP mCherry-Nuc9 MB cells in 200 µl were seeded per well in ibidi 8-well plates and incubated overnight. Cells were fixed with 4% PFA, washed, permeabilized with 0.1% Triton X-100, and blocked with 5% FBS in PBS. Primary antibodies were applied overnight at 4 °C, followed by PBS washes and DAPI staining (1:5000). Samples were mounted with VECTASHIELD and imaged on Nikon Ti2 or Leica SP8. *TJP1 quantification:* Junctional signals in D425 cells were measured using 100×100 pixel ROIs at cell-cell contacts; in MDCK-II cells, 10-pixel-wide lines were drawn along TJP1-positive junctions. Mean pixel values and 95% confidence intervals were calculated.

### MTT assay

MTT (Sigma-Aldrich) was used to assess cell viability. Log-phase cells (5–8×10³/well) were seeded in 96-well plates and cultured 24 h at 37 °C, 5% CO₂. Cells were treated with compounds in complete medium for 18–24 h in the dark, washed, and incubated with 0.5 mg/ml MTT for 2 h. After removing the MTT solution, 100 µl DMSO was added, and absorbance at 570 nm was measured. Data represent mean ± SEM of ≥3 independent experiments; DMSO-treated cells served as controls.

### Scanning electron microscopy

1×10⁴ cells were seeded on collagen I–coated coverslips in 24-well plates and treated with DMSO or 0.5 µM famlasertib for 24 h at 37 °C, 5% CO₂. Cells were fixed with 2.5% glutaraldehyde in 0.1 M cacodylate buffer, rinsed with PBS, and post-fixed in 1% OsO₄/PBS. Samples were dehydrated through ethanol series, treated with hexamethyldisilazane, air-dried, mounted on aluminum stubs, sputter-coated with 4 nm platinum, and imaged using a Zeiss GeminiSEM 450.

### Phospho-proteomics

Serum starved D425 were pretreated for 4 h with DMSO (control) or 0.25 µM famlasertib. Half of the samples were stimulated with bFGF (50 ng/ml, 5 min). Cells were then lyzed on ice in 150 µL high-SDS RIPA containing 4% w/v SDS, 1x cOmplete protease inhibitors, 1x Halt Phosphatase Inhibitor Single-Use Cocktail (ThermoFisher 78428). Lysates were transferred to pre-cooled Eppendorf tubes and snap-frozen in liquid nitrogen. Samples were stored at –80 °C for later processing. Each condition was performed in technical triplicates and exactly as described in ^26^.

### Data availability

Proteomics data have been deposited to the ProteomeXchange Consortium via PRIDE (dataset PXD062602).

### UW228-GCaMP6s Cell Line Generation and In Vitro Ca²⁺ Imaging

UW228-GCaMP6s cells were generated by lentiviral transduction of UW228 wild-type cells with pLVX-puro GCaMP6s (Addgene) followed by puromycin selection. For imaging, 4×10³ cells were seeded in ibidi µ-Slide 8-well plates and cultured for 72 h in DMEM containing 10% FBS and 1% penicillin–streptomycin. Cells were treated with 1 µM famlasertib for 24 h. Before imaging, cultures were incubated for 10 min in artificial cerebrospinal fluid (ACSF); treated groups received drug-containing ACSF. GCaMP6 imaging was performed on a Nikon Eclipse Ti2 microscope (20× objective, 488 nm excitation, 200 ms exposure). Ca²⁺ activity was recorded for 20 min at 2 s intervals. Fluorescence changes were analyzed using a custom Python pipeline and AQuA2 (MATLAB) to quantify Ca²⁺ peak amplitude (ΔF/F), event number, area, and duration.

### Zebrafish quantitative morphometric screening

Embryos were obtained by natural spawning and maintained at 28.5 °C in E3 medium. At 2 days post fertilization (dpf), dechorionated embryos were exposed to famlasertib (1–20 µM; n=25/group) or controls (E3 or 0.01% DMSO) to determine LC50. At 5 dpf, larvae were imaged using a Vertebrate Automated Screening Technology (VAST) BioImager^TM^ platform for morphometric analysis. A ResNet50-based model trained on 2250 annotated images of body area, eye size, yolk area, and swim-bladder inflation was used to quantify these structures in all the experimental images.

### Zebrafish xenografts

LA-EGFP mCherry-Nuc9 medulloblastoma cells (ONS-76, D425) were cultured at 34 °C. Cells were collected and resuspended (4×10⁶ cells/40 µl) in 0.05% polyvinylpyrrolidone. Approximately 300 tumor cells were co-injected with 5 µM famlasertib or 0.01% DMSO into the hindbrain ventricle of 48 hours post fertilization (hpf) pigment-deficient Casper embryos. Tumor-bearing larvae were anesthetized, positioned in agarose molds in 96-well plates, and imaged within 2 h post-injection (hpi).

### Imaging and treatment

Maximum intensity projections of 5 µm stacks were acquired using an Olympus IXplore SpinSR10 spinning-disk confocal microscope (20× objective). After baseline imaging (2 hpi), larvae were maintained in E3 containing 0.01% DMSO or 2.5–5 µM famlasertib at 34 °C and re-imaged at 24 and 96 hpi. Fluorescent tumors were extracted and analyzed using a custom Python pipeline to quantified longitudinal morphometric variables.

## Results

### Famlasertib is not cytotoxic and causes no developmental aberrations

Famlasertib targets the whole MAP4 family of Ser/Thr kinases ^21^. We therefore assessed *MAP4K2-7* expression in SHH (DAOY, UW228 and ONS-76) and group 3 (Grp3, D425, HD-MB03 and D283) MB cell models by qRT-PCR (Fig. 1a), excluding *MAP4K1*, due to low expression in MB ^27^. *MAP4K4* was highly expressed across all models, with the highest levels in HD-MB03 (Fig. 1a; Fig. S1a). Analysis of differential expression across human lifespan ^28^ revealed that *MAP4K4* expression peaks during early development and declines to a stable lower level after birth, whereas *MAP4K3* and *MAP4K5* remain relatively constant in the cerebellum (Fig. 1b; Fig. S1b,c).

**Figure 1.**
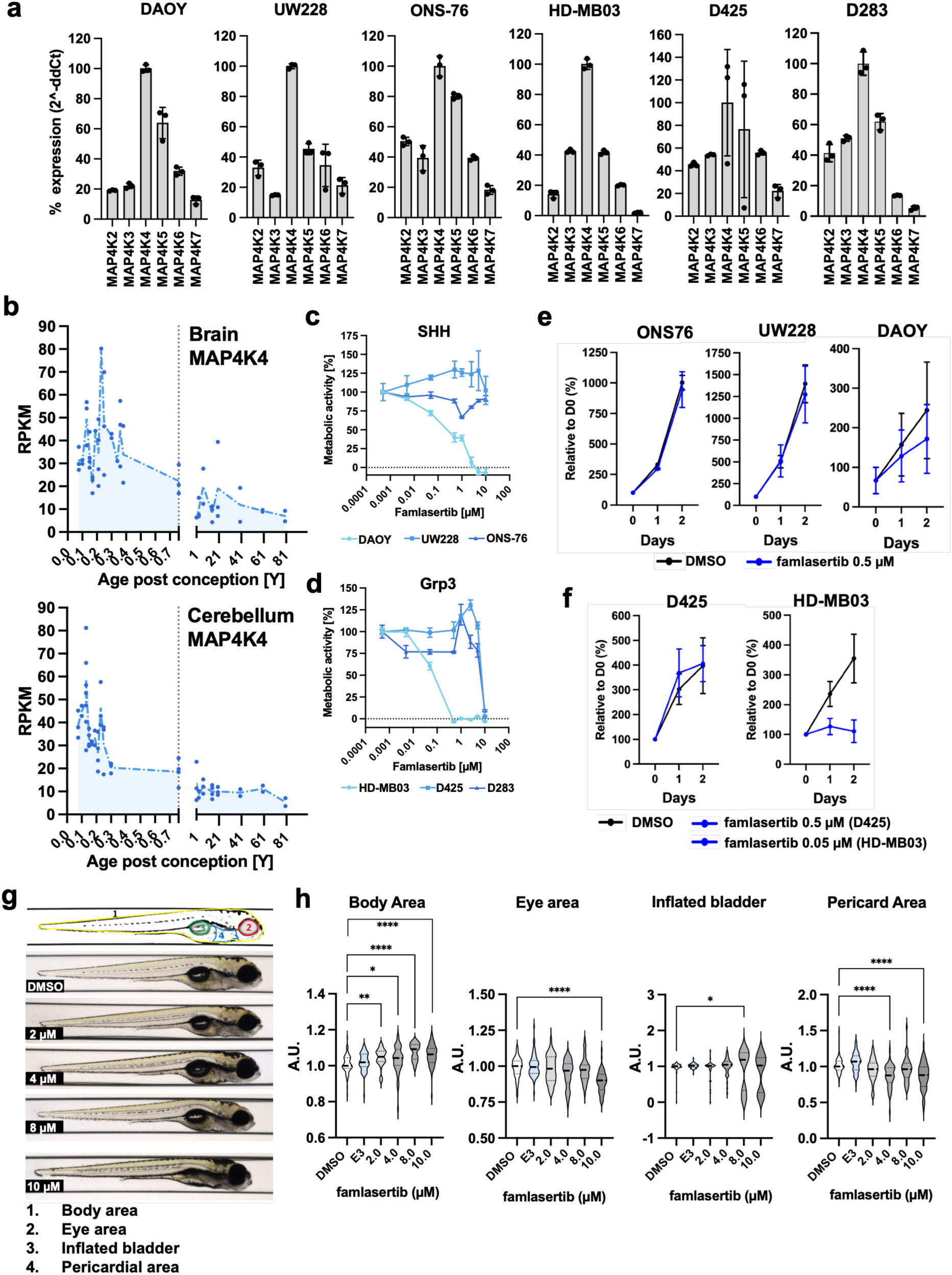
Suppressing MAP4K activity does not cause overt developmental toxicity. **a)** qRT–PCR analysis of MAP4K2–7 expression across six medulloblastoma (MB) cell models. GAPDH-normalized MAP4K mRNA expression levels are shown for three SHH (DAOY, UW228 and ONS-76) and three Group 3 (HD-MB03, D425 and D283) MB cell lines. Bar plots show mean and SD of three technical replicas. **b)** MAP4K4 expression in bulk RNA-seq datasets across the human lifespan in brain (top) and cerebellum (bottom). The x-axis indicates age in years; the vertical dashed line marks birth. The y-axis shows expression values as reads per kilobase per million mapped reads (RPKM). **c,d)** MTT assays assessing the effects of famlasertib on metabolic activity in three SHH (c) and three Group 3 (d) cell lines. MTT activity was normalized to the lowest concentration tested (0.0005 µM). Means and SD from three biological replicas are shown. **e,f)** Cell counts of SHH (e) and Group 3 (f) cell lines following 24 and 48 h of famlasertib exposure (0.5 µM). Means and SD from three technical replicas are shown. **g)** Representative VAST images of dpf 4 zebrafish larvae after 48 h of famlasertib treatment. **h)** Quantification of morphological changes during zebrafish larval development. Statistical significance for all panels: *P < 0.05, **P < 0.01, ***P < 0.001, \*\*\**P < 0.0001.* Number of analyzed larvae: DMSO (68), E3 (64), number of larvae in famlasertib treatments: 2 µM (41), 4 µM (44), 8 µM (40), 10 µM (59).

Famlasertib decreased metabolic activity in DAOY cells, measured as NAD(P)H-dependent oxidoreductase activity, while UW228 and ONS-76 cells remained unaffected (Fig. 1c). We observed the highest sensitivity to famlasertib in the Grp3 model HD-MB03, unlike D425 and D283, where an inhibitory effect was only observed at 10 µM (Fig. 1d). Famlasertib did not affect cell proliferation in ONS-76, UW228, or D425 cells (Fig. 1e,f), but markedly reduced growth rate of HD-MB03 (Fig. 1f) and induced pronounced morphological changes (Fig. S1d).

To assess potential developmental toxicity, 2-day-post-fertilization (dpf) zebrafish larvae were treated with 1 to 10 µM famlasertib for 3 days (Fig. 1g). Survival was 100% across all concentrations, with no visible malformations. Quantitative morphometric analysis revealed no major differences between control and treated groups (Fig. 1h), although some minor statistically significant differences were noted in eye-size at 10 µM and pericardic area at 4 and 8 µM. Overall, these data confirm the high tolerability of famlasertib ^29^ and reveal no overt developmental toxicity.

### Tumor cell invasion is effectively repressed by famlasertib

MAP4K4 promotes growth factor-driven invasive motility in MB ^14,15^. Using the spheroid invasion assay (SIA ^22^), we compared the effects of famlasertib and the unrelated molecule GNE-495 at 1 and 5 µM. Both compounds blocked bFGF-induced invasion, with famlasertib showing greater efficacy at 1 µM (Fig. 2a, S2a). Famlasertib’s inhibition of basal and bFGF-induced invasion was dose-dependent (Fig. 2b,c, S2b). We confirmed famlasertib inhibition of bFGF-induced invasion in a second SHH-MB cell model, UW228 (Fig. 2d). ONS-76 cells invade collagen I independent of exogenous growth factors, and famlasertib similarly suppressed ONS-76 invasion in a dose-dependent manner, albeit requiring a higher IC_50_, due to a transiently increased invasiveness at lower famlasertib concentrations (Fig. 2e).

**Figure 2.**
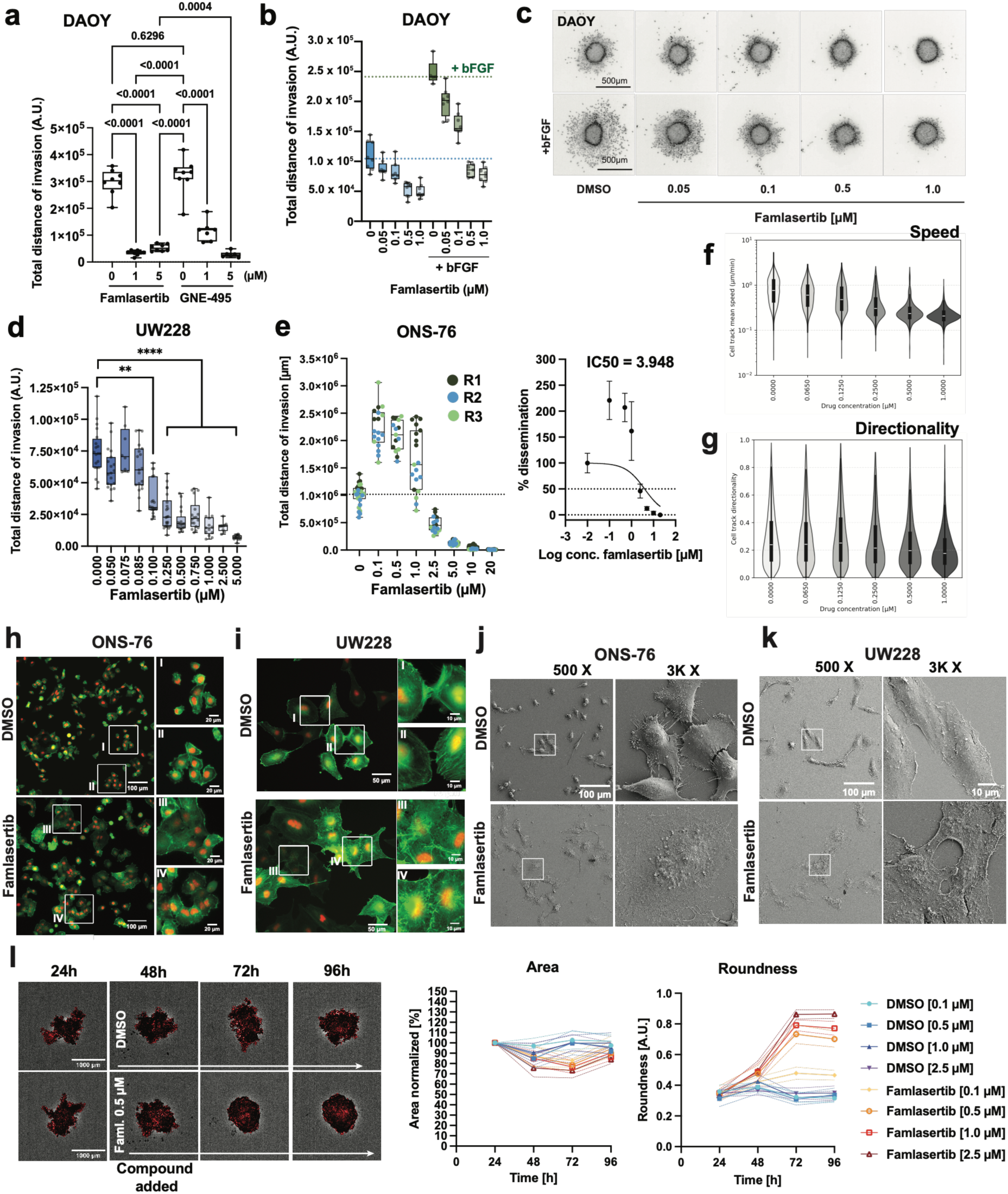
MAP4K inhibition suppresses invasive motility and promotes cell clustering. **a)** Spheroid invasion assay (SIA) of bFGF-stimulated DAOY cells (100 ng/ml in SFM), comparing the effects of famlasertib and GNE-495. DMSO served as a solvent control. Box plots show total invasion distance, calculated as the sum of each cell’s invasion distances in each spheroid. Each dot represents one independent spheroid of approximately 2500 cells (1-Way Anova, lognormal distribution, F = 107). **b)** SIA as described in (a) to assess the dose–response effects of famlasertib in the absence and presence of bFGF. Each dot represents one independent spheroid. **c)** Representative inverted grayscale images of Hoechst-stained nuclei in collagen I–embedded spheroids at the experimental endpoint shown in (b). **d)** SIA of bFGF-stimulated UW228 cells (100 ng/ml in SFM) treated with increasing concentrations of famlasertib. Pooled data from three biological replicates are shown. Each dot represents one spheroid. **e)** SIA of ONS-76 cells in SFM, treated with increasing concentrations of famlasertib. Spheroids from three independent repeats (R1-R3) are combined in box plot. **f, g)** Violin plots of mean track speed (f) and directionality (g) of ONS-76 cells treated with increasing concentrations of famlasertib. Zero concentration refers to DMSO controls. Total number of tracked objects were approximately 20000, distributed across 60 wells, with a minimum of 1250 tracks and 6 wells, and a maximum of 7000 tracks and 18 wells per condition. **h, i)**. Immunofluorescence analysis (IFA) of ONS-76 (h) and UW228 (i) cells expressing LA–EGFP and mCherryNuc following 24 h exposure to DMSO or 0.5 µM famlasertib. White boxes (I–IV) indicate regions shown at 4× magnification to the right of each panel. **j, k)** Scanning electron microscopy (SEM) images of ONS-76 (j) and UW228 (k) cells treated with DMSO or 0.5 µM famlasertib for 24 h. White boxes indicate regions shown at 6× higher magnification to the right of each panel. **l)** Spheroid growth assay. Left: representative bright field images with segmentation marks of ONS-76 cells in suspension culture exposed to DMSO or 0.5 µM famlasertib. Time stamps are hours after seeding. DMSO or compound were added 48 h after seeding. Right: Relative area (normalized to DMSO control, 24h) and roundness of ONS-76 cells in suspension culture exposed to DMSO or 0.5 µM famlasertib for 24 – 96 h.

To determine whether reduced invasiveness reflects impaired cell motility, we tracked speed and directionality of migrating cells. ONS-76 cells, which migrate consistently in 2D without stimulation (Movie 1), showed a dose-dependent decrease in speed upon famlasertib treatment, with 1 µM sufficient to reduce track mean speed to baseline (Fig. 2f, Movie 2). UW228 cells displayed a similar trend (Movies 3 and 4), though with a lower speed (Fig. S2c). Reduced migration was accompanied by decreased directionality of migration in both ONS-76 (Fig. 2g) and UW228 (Fig. S2c), indicating that famlasertib impairs both the speed and directionality control of MB cell motility.

### Famlasertib treatment increases cell clustering and altered intracellular Ca^2+^ wave dynamics

Famlasertib treatment enhances cell-cell interactions ^30^, a phenotype confirmed in ONS-76 (Fig. 2h, S2d) and UW228 (Fig. 2i, S2e) cells expressing LA-EGFP, which cluster tightly under treatment. To morphologically examine these effects, we utilized scanning electron microscopy (SEM). The morphologies of the cell models differ markedly, likely influenced by their substrate adhesion properties (Fig. 2j,k), ranging from highly adherent (e.g., UW228 and ONS-76) to poorly adherent cells (e.g., D283 and D425, S2E). SEM imaging confirmed famlasertib-induced cell flattening, increased spreading, and increased cell-cell adhesion across all cell lines except for D283 (Fig. S2f). Increased cell clustering and cluster compaction was also observed in suspension cultures of ONS-76 cells, where MAP4K inhibition was associated with pronounced spheroid formation and spheroid rounding (Fig. 2l). The increased cell-cell adherence caused by famlasertib could affect cell-to-cell communication by influencing intercellular calcium waves (ICW) through enabling Ca^2+^ or Ca^2+^ messenger transfer. We therefore explored whether the increased cell clustering after MAP4K inhibition is associated with altered [Ca^2+^]_i_ dynamics in living cells. We generated UW228 cells expressing the [Ca^2+^]_i_ sensor GCaMP6s and monitored changes in fluorescence using live cell microscopy imaging and AQuA analysis (Fig. S3a,b). In UW228 cells, increased cell-cell adhesion was associated with a moderately lower number of [Ca^2+^]_i_ peaks (DMSO: 7.26 peaks/20 mins; famlasertib: 4.77 peaks/20 mins, p=0.0017, t-test), but with no change in average peak amplitude (DMSO: 0.52 fold-change; famlasertib: 0.55 fold-change, p=0.73, t-test). Furthermore, famlasertib caused [Ca^2+^]_i_ events that simultaneously occurred or propagated across nearby cells and displayed prolonged activity duration (Fig. S3c,d). Collectively, these data demonstrate consistently increased cell clustering as a consequence of famlasertib treatment, which may be associated with altered cell-to-cell communication across junctional complexes.

### Phosphoproteomic profiling reveals novel potential functions associated with MAP4K activity

To identify proteins and kinase pathways targeted by MAP4K inhibition, we performed phosphoproteomic analysis following acute treatment (4 h) with a low concentration (0.25 µM) of famlasertib. We compared phosphosite changes upon 5 min stimulation with bFGF (50 ng/mL) in D425 group 3 MB cells, a readily bFGF-inducible cell line ^26^, in the presence or absence of 0.25 µM famlasertib (Fig. S4a). Differential phosphorylation was defined by FDR ≤ 0.05 and log₂FC ≥ 0.6 or ≤ −0.6. bFGF stimulation increased phosphorylation of 106 proteins and decreased 10 in DMSO control cells (^26^, Supplementary Table 1). In the presence of famlasertib, bFGF increased 128 phosphosites in a total of 110 proteins and decreased 15 phosphosites in a total of 14 proteins (Fig. 3a, Supplementary Table 2). 12 of the proteins differentially regulated in the presence of famlasertib are kinases, all of them with increased phosphorylation (Fig. 2a, lower), suggesting that repressing MAP4K functions by famlasertib causes rewiring of FGFR signaling. Without growth factor stimulation, famlasertib decreased phosphorylation of 145 phosphosites in a total of 122 proteins and increased 48 in a total of 43 proteins, including 11 kinases (Fig. 3b; Supplementary Table 3); MAP4K4 was the only kinase displaying both increased and decreased phosphosites, indicating on-target activity of famlasertib. In bFGF-activated state (Fig. 3c; Supplementary Table 4), famlasertib decreased phosphorylation of 154 phosphosites in a total of 127 proteins and increased 35 in a total of 31 proteins, including 11 kinases. Famlasertib together with bFGF stimulation caused phosphorylation of MAP4K4 (S710, S715), and of MAP4K5 (T168) (Fig. 3c; Supplementary Table 4). Famlasertib most strongly reduced phosphorylation of BLTP2 (T1269), OTUD7B (T547), SLC39A (T540), and Haspin (S147), while increasing ZNF507 (S195) (Fig. 3d,e). Context-specific effects were evident upon famlasertib treatment, with reduced NF-X1 (S49/S50) and MIGA2 (S726) phosphorylation in starved, and reduced PPM1H (S221) phosphorylation in bFGF stimulated cells (Fig. 3f). Together, these findings point towards MAP4K-dependent modulation of developmentally regulated transcriptional modulators, lipid transporters, and a mitochondrial fusion regulator.

**Figure 3.**
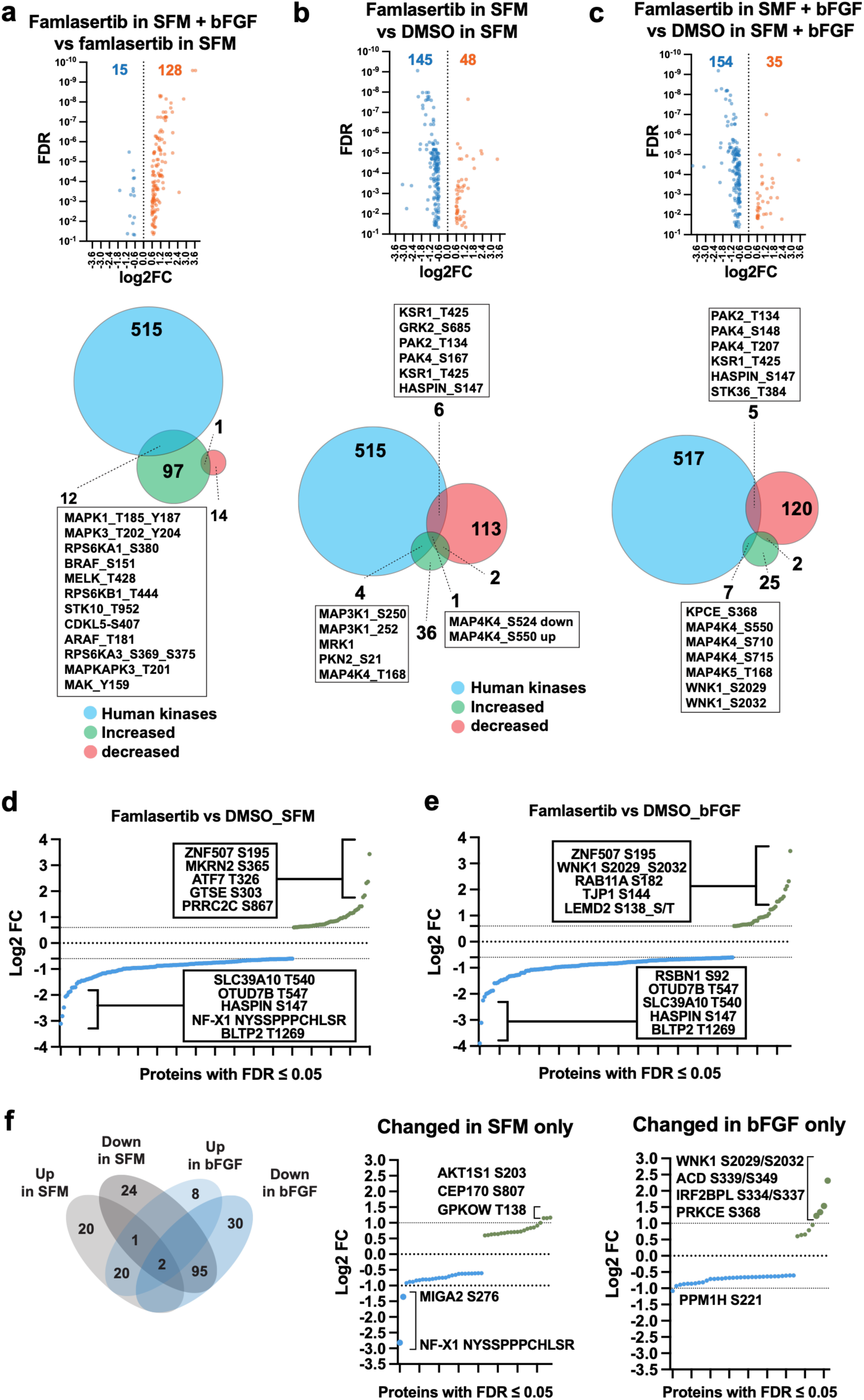
Phosphoproteomic analysis of the effects of famlasertib on global protein phosphorylation. a–c) Volcano plots illustrating phosphosite alterations following famlasertib treatment. The number of significantly upregulated (red) and downregulated (blue) phosphosites is indicated. The proportion of affected kinases relative to the human kinome (n = 526) is shown in Venn diagrams, with circle sizes proportional to the number of altered proteins. **d, e)** Top five downregulated and upregulated phosphosites after famlasertib treatment (0.25 µM) in serum-free medium (SFM; d) or SFM supplemented with 50 ng/ml bFGF (e). **f)** Left: Venn diagram of significantly up- or downregulated phosphosites after famlasertib treatment in either SFM or SFM + 50 ng/ml bFGF (log₂FC ≥ 0.6 or ≤ –0.6; FDR ≤ 0.05). Right: Top phosphosites selectively altered by famlasertib in SFM alone or in SFM supplemented with 50 ng/ml bFGF.

### MAP4K inhibition by famlasertib promotes PXN Y118 phosphorylation via PTK6

To predict relevant kinase activities based on the altered phosphorylation sites detected, we performed Kinase-Substrate Enrichment analysis (KSEA). We compared famlasertib- to DMSO-treated cells, without (Fig. 4a, Supplementary Table 5) or with bFGF stimulation (Fig. 4b, Supplementary Table 6), using an FDR cut-off of <0.1 or lower, along with a Z-score threshold of ≤ -2.4 for downregulated kinases and ≥ 2.4 for upregulated kinases. The 26 predicted up or down regulated kinase activities cover a broad spectrum of biological activities, including autophagy, cell adhesion, proliferation, survival, cell growth, protein synthesis as well as cell adhesion and motility. The predicted activation of PTK6, PAK2 and RPS6KB1 and inhibition of MAPK15, PRKD1, PBK (Fig. 4a,b) are independent of bFGF stimulation (Supplementary Table 7). We next compared the KSEA-predicted altered kinases to proteins (Fig. S4b) or mRNA (Fig. S4c) differentially expressed in medulloblastoma using the Clinical Proteomic Tumor Analysis Consortium (CPTAC) database ^27^. Notably, PDZ-binding kinase (PBK), predicted to be inhibited by famlasertib, was the only kinase among these candidates consistently upregulated in medulloblastoma at both the mRNA and protein levels (Fig. 4c). PBK is a pro-proliferative kinase upregulated in Grp 3 MB ^31^, suggesting that differential PBK dependency may underlie the selective sensitivity of HDM-MB03 (Fig. 1d, f).

**Figure 4.**
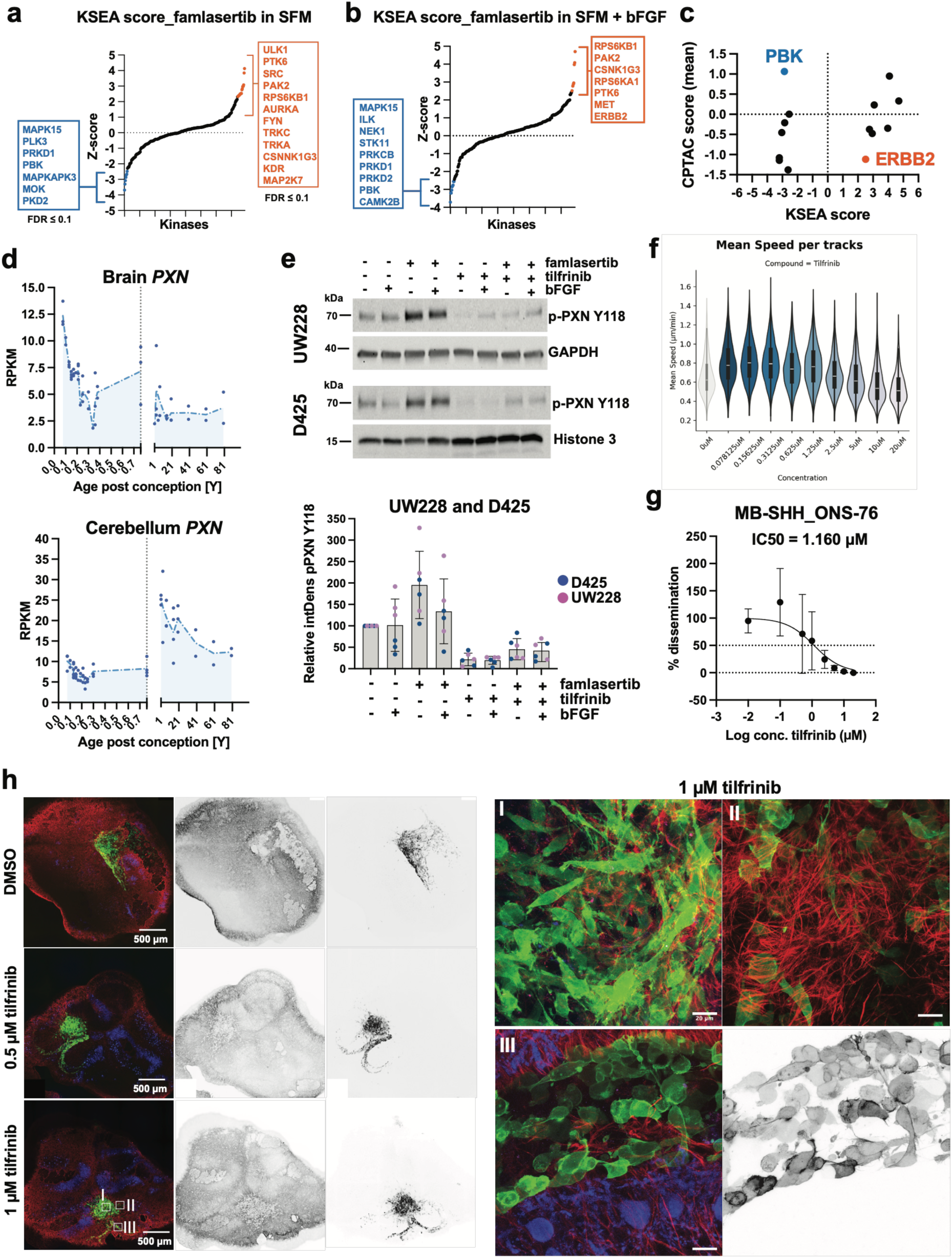
PXN Y118 is a MAP4K-regulated phosphosite that is repressed by tilfrinib. a, **b)** Kinase–substrate enrichment analysis (KSEA) scores predicting the effects of famlasertib in serum-free medium (SFM; A) or SFM supplemented with bFGF (B). An FDR cutoff of <0.1 was applied, together with a Z-score threshold of ≤ −2.4 for downregulated kinases and ≥ 2.4 for upregulated kinases. **c)** Integration of KSEA-predicted kinase activity scores with CPTAC protein expression data predicts repression of PBK by famlasertib, a kinase upregulated in MB tumors. **d)** Paxillin (PXN) expression across the human lifespan based on bulk RNA-seq datasets from brain (top) and cerebellum (bottom). The x-axis denotes age in years; the vertical dashed line indicates birth. The y-axis shows expression values in reads per kilobase per million mapped reads (RPKM). **e)** Upper: Representative immunoblot (IB) analysis of PXN Y118 phosphorylation in D425 and UW228 cells. Lower: Quantification of four independent IB experiments performed in D425 and UW228 cells. Each dot represents the quantification of an independent experiment. **f)** Violin plots showing track mean speed of ONS-76 cells treated with increasing concentrations of tilfrinib. On average, 2881 (min 1143, max 4168) tracks were analyzed per condition. **g)** Spheroid invasion assay (SIA)-based determination of the tilfrinib IC₅₀ for bFGF-independent invasiveness in ONS-76 cells. **h)** Evaluation of tilfrinib effects on ONS-76 cell invasion in organotypic cerebellar slice culture. Immunofluorescence analysis of ONS-76 spheroids implanted into cerebellar slices and treated with 0.5 and 1 µM tilfrinib for five days. Green: LA–EGFP; red: GFAP. I, II and III show magnifications of representative areas highlight with white squares. I: region of tumor cell cluster core, II: invasion front of individually invading cells, III: stream of collectively invading cells. Black and white image shows inverted grey-scale image F-actin cytoskeleton (LA-eGFP channel).

Paxillin (PXN) Y118, a known substrate of SRC kinases and PTK6, showed significantly increased phosphorylation following famlasertib treatment in serum-starved cells (Supplementary Table 3). *PXN* is a developmentally regulated gene, displaying increased expression in the cerebellum after birth (Fig. 4d). We confirmed the predicted increase in PXN-pY118 after MAP4K inhibition by immunoblotting in D425 and UW228 cells (Fig. 4e) and found that PXN-pY118 was reduced to baseline by co-treatment with the PTK6 inhibitor tilfrinib ^32^; tilfrinib also suppressed basal PXN-Y118 phosphorylation in both cell lines. In control cells, PXN-pY118 predominantly colocalizes with F-actin–rich lamellipodial protrusions (Fig. S4d). Famlasertib treatment promoted a loss of large F-actin–rich lamellipodia and increased cell–cell contacts. Under these conditions, PXN-pY118 was detected in cortical F-actin–containing filamentous protrusions. Tilfrinib treatment had little effect on lamellipodia and PXN-pY118 localization and only partially restored lamellipodial localization of PXN-pY118 in famlasertib–treated cells. Tilfrinib treatment at effective concentration did not affect single cell motility of ONS-76 cells and a dose-dependent reduction in track mean speed was only observed above 5 µM concentration (Fig. 4f). In contrast, a dose-dependent decrease in collagen I invasion of ONS-76 cells, with a calculated IC₅₀ of 1.16 µM (Fig. 4g), was observed. Surprisingly, however, we could not confirm reduced invasion of tilfrinib-treated cultures in the tissue context (Fig. 4h), where we still observed single and collectively invading cells at a concentration that blocks PXN-pY118 and invasion.

In conclusion, the observed phosphoproteomic alterations indicate substantial rewiring of cellular signaling by famlasertib. One consequence related to cytoskeletal control is enhanced PXN-pY118 phosphorylation, which can be reversed by the PTK6 inhibitor tilfrinib.

### Famlasertib target kinases are involved in cell-cell interaction signaling

The phosphoproteomic analysis revealed decreased phosphorylation of talin (T144), afadin (T566), delta-catenin (T177, T916), MAGI1 (S1261/5), and TJP1 (T589, T1142, T1167), and increased phosphorylation of TJP1 (S144, T1124) upon famlasertib treatment (Fig. 5a, Supplementary Tables 3, 4, Fig. S5a), which provides some candidate proteins potentially involved in increased cell-cell adhesion. Among these proteins, TJP1 exhibited the most significant changes, with several phosphosites affected (Fig. 5b), and famlasertib-treated UW228 (Fig. 5c) and ONS-76 (Fig. S5b) displayed TJP1 enrichment in cell-cell contacts. These contacts displayed a jagged appearance, with the TJP1 staining highlighting structures aligned with filamentous actin and stress fibers. In D425 cells, which grow as cell clusters, famlasertib treatment similarly induced the accumulation of TJP1 in cell-cell contacts, evident as distinct peaks of fluorescence between individual cells (Fig. 5d). The increased accumulation of TJP1 in regions of cell-cell contacts after famlasertib treatment was also apparent when the cells were embedded in organotypic cerebellum slices (Fig. 5e,f). bFGF stimulation did not affect TJP1 distribution in the tumor tissue, while it caused a significant reduction in astrocytic processes.

**Figure 5.**
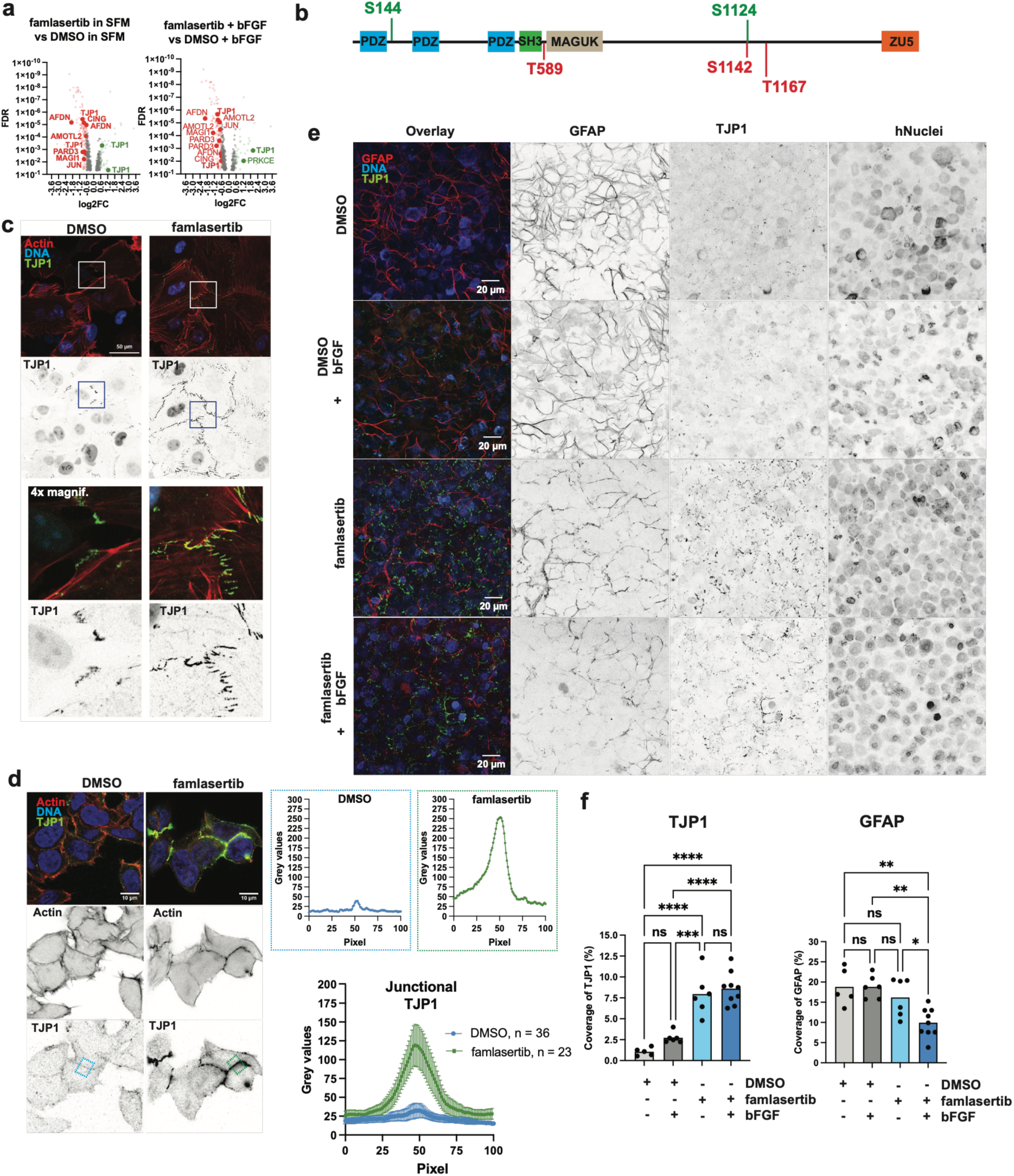
Famlasertib-dependent changes in TJP1 phosphorylation are associated with enhanced localization of TJP1 at cell–cell contacts. **a)** Volcano plots highlighting phosphosites significantly altered upon famlasertib treatment. Junctional proteins and proteins involved in the regulation of cell–cell contacts with altered phosphosites are indicated. Significance was defined as FDR ≤ 0.05 and a log₂ fold change (log₂FC) ≥ 0.6 (upregulated) or ≤ −0.6 (downregulated). **b)** Schematic representation of the TJP1 protein structure and phosphorylation sites altered in the presence of famlasertib. Upregulated sites are shown in green and downregulated sites in red. **c)** IFA of TJP1 localization in UW228 cells cultured in the absence or presence of 0.5 µM famlasertib for 24 h. Representative inverted grayscale images of TJP1 are shown to enhance contrast. Boxes indicate regions displayed below at 4× magnification. Scale bar, 50 µm. **d)** IFA of TJP1 localization in D425 cells treated as in (c). Representative inverted grayscale images of F-actin and TJP1 are shown for improved contrast. Dotted squares mark representative 100 × 100 pixels cell-cell contact regions (DMSO: n = 36, famlasertib: n = 26) used to quantify TJP1 signal intensity at cell–cell contacts in DMSO-treated (blue) and famlasertib-treated (green) cells. Quantification of gray value intensities is shown to the right of the IFA panels. **e)** Maximum intensity projections of confocal IFA of TJP1 localization in D425 cells implanted into organotypic cerebellar slice cultures (OCSCs) and maintained for five days with or without 1 µM famlasertib, in the absence or presence of bFGF. **f)** Quantification of GFAP and TJP1 coverage under the indicated treatment conditions (One-way ANOVA, F-values (1.53 (GFAP) and 53.17 (TJP1), N=3 independent slice per conditions; with 2-4 regions of interest analyzed per replicate).

In MDCK-II cells, where TJP1 localizes robustly to cell-cell contacts (Fig. 6a. S5c), famlasertib caused cell compaction, phenotypically resembling TJP1 KO cells (Fig. S5c,d), an increase in absolute apical junctional density of TJP1 (Fig. 6b), and cytosolic re-distribution at basal cell regions (Fig. 6a, S5d). These alterations are associated with increased basal F-actin and stress fibers (Fig. 6a). Famlasertib caused cell compaction and a concomitantly reduced cell perimeter (Fig. 6c), but relative cortical TJP1 mean intensity in these cells remained markedly higher compared to control cells (Fig. 6d).

**Figure 6.**
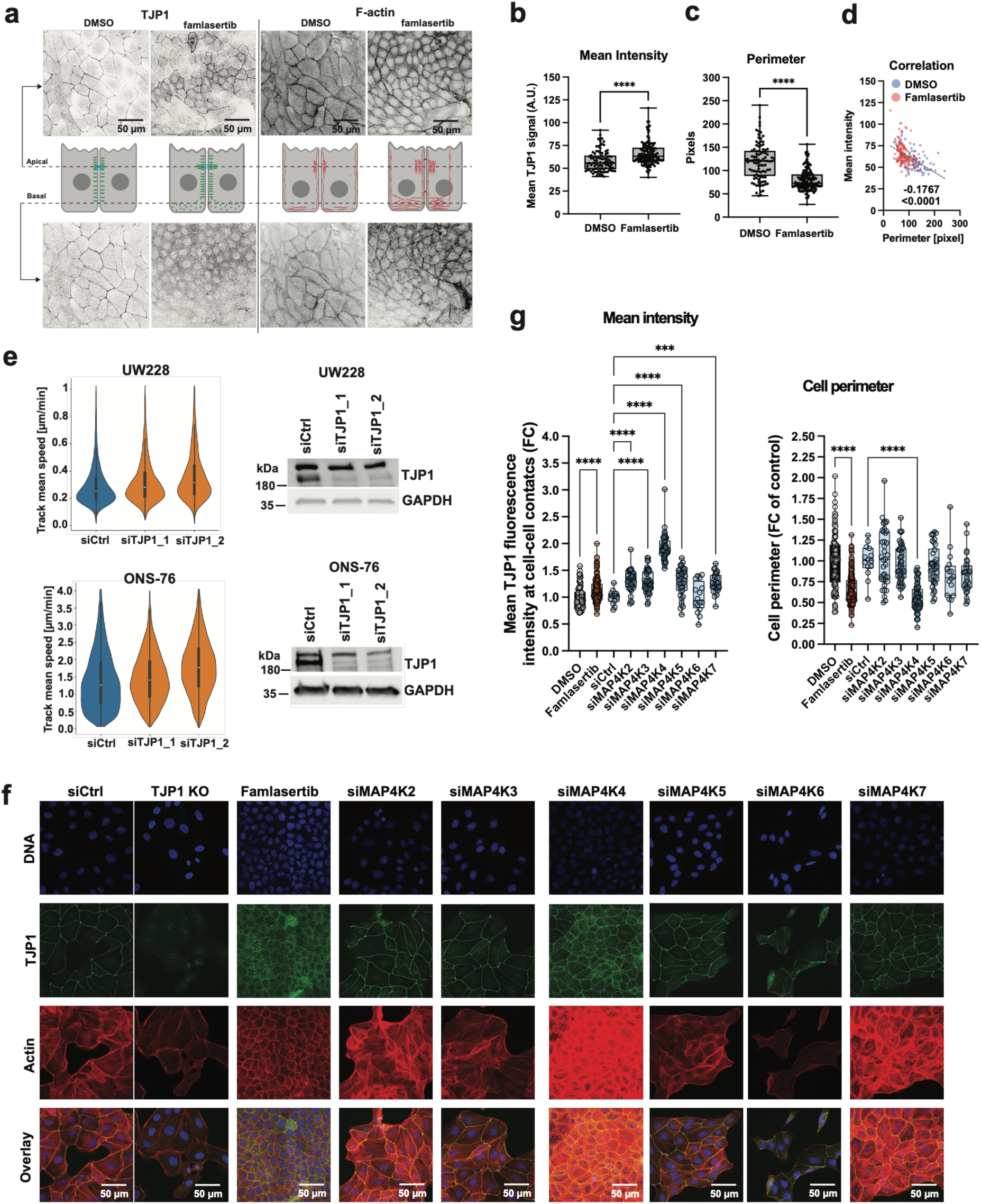
Famlasertib promotes accumulation of TJP1 in apical junctions. **a)** Representative confocal microscopy images of TJP1 and F-actin in basal and apical confocal microscopy sections of MDCK cells treated with DMSO or 0.5 µM famlasertib for 24 h. The schematic indicates the approximate positions of the cross-sections and the corresponding localization patterns of TJP1 and F-actin. **b)** Quantification of mean TJP1 fluorescence intensity at cell–cell contacts in DMSO-treated control cells and cells exposed to 0.5 µM famlasertib for 24 h, shown as fold change relative to DMSO controls. Statistical significance was determined using unpaired t test with Welch’s correction; p < 0.05 is considered significant. Center line represents mean and whiskers extend to minimum and maximum values, including individual data points (DMSO: n=96 cells; famlasertib: n=138 cells). **c)** Quantification of cell perimeter, presented as fold change relative to DMSO controls. Statistical significance was determined using unpaired t test with Welch’s correction; p < 0.05 is considered significant. Center line represents mean and whiskers extend to minimum and maximum values, including individual data points (DMSO: n=96 cells; famlasertib: n=138 cells). **d)** Ratio of mean TJP1 intensity to cell perimeter, demonstrating a negative correlation between junctional TJP1 intensity and perimeter length. The dotted square highlights cells with comparable perimeters. **e)** siRNA-mediated depletion of TJP1 using two independent siRNAs in UW228 and ONS-76 cells. Left: Violin plots showing track mean speed of siCtrl- and siTJP1-treated UW228 and ONS-76 cells. Right: Corresponding immunoblot controls confirming efficient TJP1 depletion in both cell lines. 300 – 800 cells were anlyzed per condition (mean: 500 cells). **f)** Immunofluorescence analysis of MDCK cells transfected with siCtrl or siRNAs targeting individual MAP4Ks. Famlasertib treatment served as a control. **g)** Quantification of mean TJP1 fluorescence intensity at cell–cell contacts and cell perimeter measurements, normalized to control conditions. Statistical significance was determined using ordinary one-way ANOVA with Sidak’s multiple comparisons test; p < 0.05 is considered significant. Center line represents mean and whiskers extend to minimum and maximum values, including individual data points (DMSO: n=96 cells; famlasertib: n=138 cells; siCtrl: n=14 cells; siMAP4K2: n=38 cells; siMAP4K3: n=48 cells; siMAP4K4: n=64 cells; siMAP4K5: n=38 cells; siMAP4K6: n=16 cells; siMAP4K7: n=30 cells).

The depletion of TJP1 in UW228 and ONS-76 cells increased track mean speed significantly, suggesting that TJP1 restricts single cell motility of MB tumor cells (Fig. 6e). In an approach to identify the MAP4K associated with altered TJP1 distribution, we used siRNAs to deplete MAP4K2-7 in MDCK-II cells (Fig. 6f). Only the depletion of MAP4K4 causes a phenotype similar to that observed after famlasertib treatment, characterized by increased junctional TJP1 and a reduction in cell perimeter (Fig. 6g).

In summary, famlasertib alters phosphorylation of key cell–cell contact regulators, particularly TJP1, promoting its junctional accumulation, enhanced adhesion, and compaction. Similar effects upon MAP4K4 depletion implicate this kinase in TJP1 regulation, while loss of TJP1 increases 2D cell motility.

### Famlasertib prevents tissue invasion of MB tumor cells

ONS-76 cells effectively invade cultured organotypic cerebellar slices (OCSCs) ^33^. To assess the migrastatic impact of famlasertib in the physiologically relevant tissue context, we implanted ONS-76 tumor spheroids in OCSCs and treated them either with DMSO or 1 µM famlasertib for 5 days. This dosage corresponds to the IC25 in the ONS-76 3D invasion assay (Fig. 2e) and approximately half of the maximal brain concentration achieved *in vivo* after 10 mg/kg intraperitoneal administration ^21^. At the endpoint, slices were fixed and tumor growth and invasion analyzed by confocal microscopy. Control tumor cells invaded cerebellar tissue efficiently, spreading up to 1.5 mm from the implantation site (Fig. S6a). bFGF stimulation further enhanced dispersion, resulting in near-complete dissociation of cells, with only a few cells loosely connected by filamentous structures. Although 1 µM famlasertib did not fully block tissue invasion (Fig. S5a), it shifted invasion from predominantly individual to more collective modes and increased cell–cell adhesion both with and without bFGF (Fig. S6b). Increasing famlasertib to 2.5 and 5.0 µM—approximately the IC50 and equivalent to one- and twofold brain exposure after 10 mg/kg dosing—resulted in a near-complete inhibition of invasion, with no distantly disseminating cells observed (Fig. 7a, S6c). High-resolution imaging furthermore showed that most invading cells in DMSO controls remained connected by few long processes, whereas famlasertib-treatment caused pronounced morphological alterations and the extensive formation of thin cellular protrusions (Fig. 7b).

**Figure 7.**
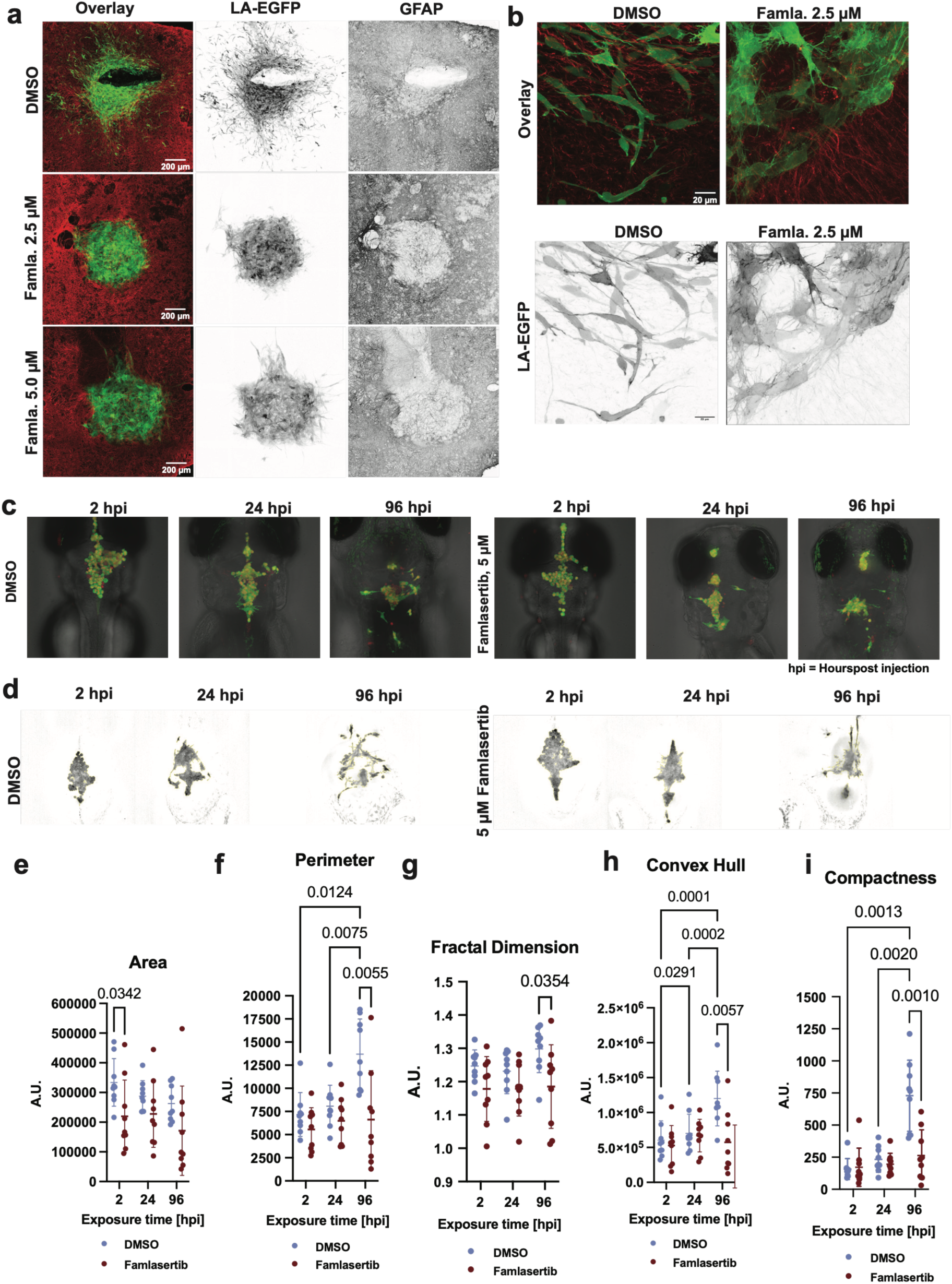
Famlasertib suppresses invasion and tumor cell dissemination in tissue context and in vivo in zebrafish. **a)** Maximum intensity projections of confocal IFA of organotypic cerebellar slice cultures implanted with ONS-76 spheroids and treated for 5 days with famlasertib (2.5 or 5.0 µM). Inverted grayscale images of LA–EGFP (tumor cells) and GFAP (glial cells within the tumor microenvironment) are shown to enhance contrast. Green: LA– EGFP; red: GFAP. **b)** Higher-magnification views of the invasion fronts in DMSO- and famlasertib-treated cultures shown in (a). **c)** Multichannel confocal imaging with brightfield anatomical context of orthotopically xenotransplanted ONS-76 cells in zebrafish larvae brain. Representative images from one DMSO-treated (control) and one 5 µM famlasertib-treated larva at 2, 24, and 96 hpi. Tumor cells are shown in green (LA–EGFP) and nuclei in red (mCherryNuc). **d)** Grayscale images of LA–EGFP fluorescence in orthotopically xenotransplanted ONS-76 cells at 2, 24, and 96 hours post injection (hpi) from one representative larvae treated with DMSO and another treated with 5uM famlasertib. **e–h)** Quantification of tumor-associated parameters based on LA–EGFP fluorescence in DMSO- and famlasertib-treated zebrafish larvae at 2, 24, and 96 hpi. The experiment was independently repeated twice with consistent results; data from one representative experiment is shown. Statistical significance was determined using two-way ANOVA with Tukey’s multiple comparisons test (paired design), with p < 0.05 considered significant. Data are presented as median (center line) with quartile range (whiskers) and individual data points (DMSO: n= 9 larvae; famlasertib: n= 9 larvae).

### Famlasertib reduces tumor growth and complexity in zebrafish medulloblastoma xenograft model

To investigate the *in vivo* efficacy of famlasertib on tumor cell growth and dissemination, ONS-76 cells expressing LA-EGFP and nuclear mCherry were orthotopically injected into the brains of Casper mutant zebrafish larvae in solutions containing either DMSO or 5 µM famlasertib. Larvae bearing microtumors were imaged at baseline (2 hours post injection, 2 hpi) and subsequently maintained under continuous drug exposure by supplementing the medium with DMSO or 5 µM famlasertib. Imaging was repeated at 24 and 96 hpi (Fig. 7c,d S7), and tumor-associated parameters quantified (Fig. 7e-i). Acute exposure to 5 µM famlasertib resulted in more compact tumor cell clusters compared with DMSO controls, as evidenced by a reduction in tumor area (p=0.0342) as early as 2 hpi (Fig. 7e). The immediate cell compaction *in vivo* is consistent with the compaction observed in 3D cell clusters *in vitro* (Fig. 2l). The trend toward reduced area in famlasertib-treated groups persisted over subsequent days, although differences compared to controls did not reach statistical significance. ONS-76 xenografts in DMSO-treated larvae displayed progressive dissemination into the surrounding brain parenchyma over four days, reflected by a significant increase in tumor perimeter between 2 and 96 hpi (p = 0.0124) and between 24 and 96 hpi (p=0.0075) (Fig. 7f). In larvae exposed to famlasertib, tumor spread was attenuated, and no increase in perimeter between 24 and 96 hpi was noted. Moreover, tumor perimeter in control larvae was significantly larger at endpoint (p=0.0055). Consistent with an invasion repressed state, famlasertib-treated tumors displayed fewer F-actin protrusions and a significant reduction in fractal dimension at 96 hpi (p = 0.0354, (Fig. 7g)), convex hull complexity (p = 0.0057, (Fig. 7h)) and compactness (p = 0.0010, (Fig. 7i)) . Collectively, these parameters reflect tumor irregularity and invasive behavior ^34^.

Together, these findings demonstrate that famlasertib represses the acquisition of invasive morphological features *in vivo*, which is associated with reduced dissemination of the tumor cells.

## Discussion

Migrastatic compounds could offer a strategy to limit local invasion and metastasis ^17,35,36^, a critical need in pediatric brain tumors where complete resection is often unfeasible, and treatment escalation causes severe long-term effects. Here, we demonstrate the migrastatic potential of the neuroprotective MAP4K inhibitor famlasertib *in vitro* and *in vivo*, define downstream signaling pathways, and identify TJP1 as a novel MAP4K effector.

MAP4Ks regulate diverse processes, including motility, invasion, and growth control ^37^, and the effects of famlasertib thus likely reflect inhibition of multiple MAP4Ks and their downstream effectors. One of these, MAP4K4, is highly expressed in MB and promotes growth factor–driven invasion via integrin activation and phosphorylation of cytoskeletal regulators such as PKC-θ and VASP ^14,15^. MAP4K4 has primarily been linked to motility through regulation of F-actin and cell–matrix adhesion. However, famlasertib-treated MB cells showed increased cell–cell adhesion and junctional accumulation of TJP1, indicating that additional mechanisms including cell-cell adhesion signaling are regulated by MAP4K4 or the other MAP4K family members. The jagged TJP1 distribution resembles a previously described MAP4K4-dependent phenotype with reduced junctional tension and restricted collective migration ^38^. Consistently, MAP4K4 was found to localize to adherens junctions, suggesting that it may regulate TJP1 phosphorylation and stability locally ^39^. Loss of TJP1/2 promotes cell compaction, reduced epithelial sheet migration, and actomyosin remodeling, pointing towards negative mechanical feedback control, orchestrated at tight junctions (TJs) to prevent excessive contraction and pulling ^39^. Similarly, MAP4K4 depletion in MDCK-II cells induced TJP1 relocalization, compaction, and increased stress fiber formation, supporting a role for MAP4K4 in (TJ) regulation. TJP1 localization and function are regulated by actin dynamics and site-specific phosphorylation, thereby controlling phase separation mechanisms required for tight junction assembly ^40^. Among the famlasertib-sensitive sites, T589—near the SH3 and MAGUK domains—may influence TJP1 conformation or phase behavior, whereas regions around S1124 and T1142 are linked to F-actin interactions. These findings suggest that MAP4Ks regulate TJP1 via site-specific phosphorylation. However, whether changes at T589, T1142, and T1167 reflect direct kinase inhibition or secondary effects of altered cytoskeletal organization remains unclear. Notably, S1142 shows context-dependent regulation upon famlasertib treatment, increasing without bFGF but decreasing upon stimulation, pointing to an indirect mechanism. Overall, our data link differential TJP1 phosphorylation to its localization, warranting further mechanistic investigation.

One consequence of pharmacological MAP4K inhibition in MB cells is increased PXN pY118. KSEA identified PTK6 as a possible upstream regulator, and PTK6 inhibition in famlasertib-treated cells reversed PXN Y118 phosphorylation. *PXN* expression rises postnatally in the cerebellum ^28^, which may reflect a role in axonal pathfinding ^41^, but remains low in MB at both mRNA and protein levels ^27^, arguing against a major role in tumor growth. Although PXN pY118 is generally linked to increased motility, its effects are context-dependent ^42^: *in vitro,* reduced phosphorylation impairs migration, whereas *in vivo* it can enhance it. PTK6 inhibition with tilfrinib did not affect invasion in the tissue model, indicating that PTK6 is not involved in MB invasion in the tissue context.

Famlasertib markedly inhibited tissue invasion of the highly invasive MB cell line ONS-76, and reduce spreading of the orthotopically implanted tumor cells in zebrafish larvae *in vivo,* demonstrating its migrastatic potential in a physiologically-relevant context. Reduced invasion was accompanied by a pronounced morphological shift, with cells exhibiting numerous filamentous protrusions. The molecular basis of this transformation remains unclear but may involve the altered phosphorylation of transcriptional regulators, such as NFX1, ZNF507, or ATF7 we observed. However, the functional relevance of the affected phosphosites in these proteins is unknown, and their contribution remains speculative. Alternatively, transcription-independent mechanisms may be involved, including ER-associated lipid transfer proteins such as MIGA2 and BLTP2, which link the ER and mitochondria ^43,44^. Notably, BLTP2 regulates plasma membrane fluidity by increasing phosphatidylethanolamine levels and promoting cancer cell dissemination ^44^, whereas a cancer-related role for MIGA2 has not yet been established.

In conclusion, our study identifies MAP4K signaling as a therapeutically targetable regulatory process of invasion in medulloblastoma and establishes famlasertib as a promising migrastatic agent that restrains tumor cell dissemination without causing developmental toxicity. Beyond its established roles in cytoskeletal regulation, MAP4K inhibition reveals an additional layer of control over cell–cell adhesion, highlighting junctional remodeling as a potential mechanism limiting invasion. While the precise molecular pathways linking MAP4Ks to TJP1 phosphorylation and cytoskeletal dynamics require further investigation, our findings provide a framework for targeting invasive behavior in MB and support the clinical exploration of migrastatic strategies to complement current therapies.

## Acknowledgments

We thank Prof. Alf Honigmann for generously providing the MDCK-II TJP1 WT and KO cells. This project was funded by projects from the Swiss National Science Foundation (Sinergia_CRSII5_202245/1, SNF_310030_188793), the Stiftung für Wissenschaftliche Forschung an der Universität Zürich (STWF-22-005), and Childhood Cancer Research Foundation Switzerland. Mass-spectrometry was performed at the Functional Genomic Center Zürich and imaging was performed with equipment maintained by the Center for Microscopy and Image Analysis, University of Zurich. We thank those two facilities for their support.

## Author contributions

**M.T.S.:** Methodology, Investigation, Formal analysis (Figures 2d, i, 4g, 6a–d, f, g, S5c, d), Visualization (figure preparation and assembly)

**M-S.L.:** Conceptualization (slice culture model), Methodology, Investigation, Formal analysis (Figures 5e, f, 6a, b, S3a-d, S5b, S6a,c), Visualization (figure preparation and assembly)

**S.Y.:** Investigation, Formal analysis (Figures 1c, e, 2a–c, h, 5c, d, S1d, S2a, b, d)

**V.A.:** Conceptualization (in vivo study), Methodology, Investigation, Formal analysis, Data curation (zebrafish larvae in vivo study and bioimaging data), Visualization (figure preparation and assembly)

**A.A.:** Methodology (*in vivo* zebrafish bioimage analysis), Formal analysis (zebrafish bioimaging data, Figures 1, 7)

**B.C.:** Methodology (single cell tracking, imaging), Formal analysis (single-cell motility quantification), Investigation (in vitro bioimage data acquisition and analysis)

**D.V.:** Investigation, Formal analysis (Figures 1a, 2 j, k, 6 e, f, S1a, S2f, S4e)

**L.L.K.:** Investigation, Formal analysis (Figure 3, Figure 4a, b)

**D. H.:** Investigation (Figure S6b)

**T.B.:** Investigation (Figure 4e, S4d)

**A.W.:** Investigation (Figure 4f)

**S.C.F.N.:** Conceptualization (in vivo study), Formal analysis, Data curation, Resources, Funding acquisition, Writing – Review & Editing

**M.B.:** Conceptualization, Supervision, Resources, Funding acquisition, Formal analysis, Visualization (figure preparation and assembly), Writing – Original Draft

## Statements

- The authors declare no competing interests.
- All animal experiments were performed in accordance with the Swiss Federal Act on Animal Protection (TSchG) and the Swiss Animal Protection Ordinance (TSchV), and were approved by the Cantonal Veterinary Office of Zürich, license numbers: ZH079/2023 (mice) and ZH073/2024 (zebrafish larvae).
- During the preparation of this work, the author(s) used Claude.app for proofreading the figure legends in order to increase clarity and consistence. After using this tool, the author(s) reviewed and edited the content as needed and take(s) full responsibility for the content of the published article.

---

**Movie 1:** ONS-76 LA-eGFP cells in complete growth medium exposed to DMSO. Example tracking analysis movie showing detected objects and corresponding tracks fading over time color coded according to track ID (different color indicates different object). Blinking red circles indicates bad tracks (or fragments) not fulfiling the exclusion criteria for further analysis (60 min duration). Time stamp in hours:minutes.

**Movie 2:** ONS-76 LA-eGFP cells in complete growth medium exposed to 1 µM famlasertib. Example tracking analysis movie showing detected objects and corresponding tracks fading over time color coded according to track ID (different color indicates different object). Blinking red circles indicates bad tracks (or fragments) not fulfiling the exclusion criteria for further analysis (60 min duration). Time stamp in hours:minutes.

**Movie 3:** UW228 LA-eGFP cells in complete growth medium exposed to DMSO. Example tracking analysis movie showing detected objects and corresponding tracks fading over time color coded according to track ID (different color indicates different object). Blinking red circles indicates bad tracks (or fragments) not fulfiling the exclusion criteria for further analysis (60 min duration). Time stamp in hours:minutes.

**Movie 4:** ONS-76 LA-eGFP cells in complete growth medium exposed to 1 µM famlasertib. Example tracking analysis movie showing detected objects and corresponding tracks fading over time color coded according to track ID (different color indicates different object). Blinking red circles indicates bad tracks (or fragments) not fulfiling the exclusion criteria for further analysis (60 min duration). Time stamp in hours:minutes.

**Supplementary Table 1:** Comparison of a phosphosite alterations in DMSO control D425 cells + bFGF compared to DMSO control cells – bFGF (FDR ≤ 0.05 and log₂FC ≥ 0.6 or ≤ −0.6). Comparison: DMSO + bFGF vs DMSO – bFGF. Treatments: Starvation:18h, bFGF: 50 ng/mL, 5 min

**Supplementary Table 2:** Significantly (FDR ≤ 0.05 and log₂FC ≥ 0.6 or ≤ −0.6) altered phosphosites of the comparison of famlasertib-treated D425 cells + bFGF compared to famlasertib-treated D425 cells - bFGF. Comparison: Famla. + bFGF vs Famla. – bFGF. Treatments: Starvation:18h, famlasertib: 4h, 250 µM, bFGF: 50 ng/mL, 5 min

**Supplementary Table 3:** Significantly (FDR ≤ 0.05 and log₂FC ≥ 0.6 or ≤ −0.6) altered phosphosites of the comparison of famlasertib-treated D425 cells - bFGF compared to DMSO-treated D425 cells - bFGF. Comparison: Famla. – bFGF vs DMSO – bFGF. Treatments: Starvation:18h, famlasertib: 250 µM, 4h

**Supplementary Table 4:** Significantly (FDR ≤ 0.05 and log₂FC ≥ 0.6 or ≤ −0.6) altered phosphosites of the comparison of famlasertib-treated D425 cells + bFGF compared to DMSO-treated D425 cells + bFGF. Comparison: Famla. + bFGF vs DMSO + bFGF. Treatments: Starvation:18h, famlasertib: 250 µM, 4h, bFGF: 50 ng/mL, 5 min

**Supplementary Table 5:** Kinase Substrate Enrichment Analysis (KSEA)-predicted activated or repressed kinases (FDR ≤ 0.1 and z-score ≥ 2.4 or ≤ −2.4) from the comparison of famlasertib-treated D425 cells - bFGF compared to DMSO-treated D425 cells - bFGF. Comparison: Famla. – bFGF vs DMSO – bFGF. Treatments: Starvation: 18h, famlasertib: 250 µM, 4h

**Supplementary Table 6:** Kinase Substrate Enrichment Analysis (KSEA)-predicted activated or repressed kinases (FDR ≤ 0.1 and z-score ≥ 2.4 or ≤ −2.4) from the comparison of famlasertib-treated D425 cells + bFGF compared to DMSO-treated D425 cells + bFGF. Comparison: Famla. + bFGF vs DMSO + bFGF. Treatments: Starvation: 18h, famlasertib: 250 µM, bFGF: 50 ng/mL, 5 min 4h.

**Supplementary Table 7:** Kinase Substrate Enrichment Analysis (KSEA)-predicted activated or repressed kinases (FDR ≤ 0.1 and z-score ≥ 2.4 or ≤ −2.4) from the comparison of famlasertib-treated D425 cells + bFGF compared to famla-treated D425 cells - bFGF. Comparison: Famla. + bFGF vs Famla. - bFGF. Treatments: Starvation: 18h, famlasertib: 250 µM, bFGF: 50 ng/mL, 5 min 4h

**Supplementary Table 8:** RT-qPCR primers and siRNAs used

