## Supplementary figures for "MAP4K inhibition by the CNS penetrant inhibitor famlasertib restrains medulloblastoma dissemination without developmental toxicity"

#### Figure S1

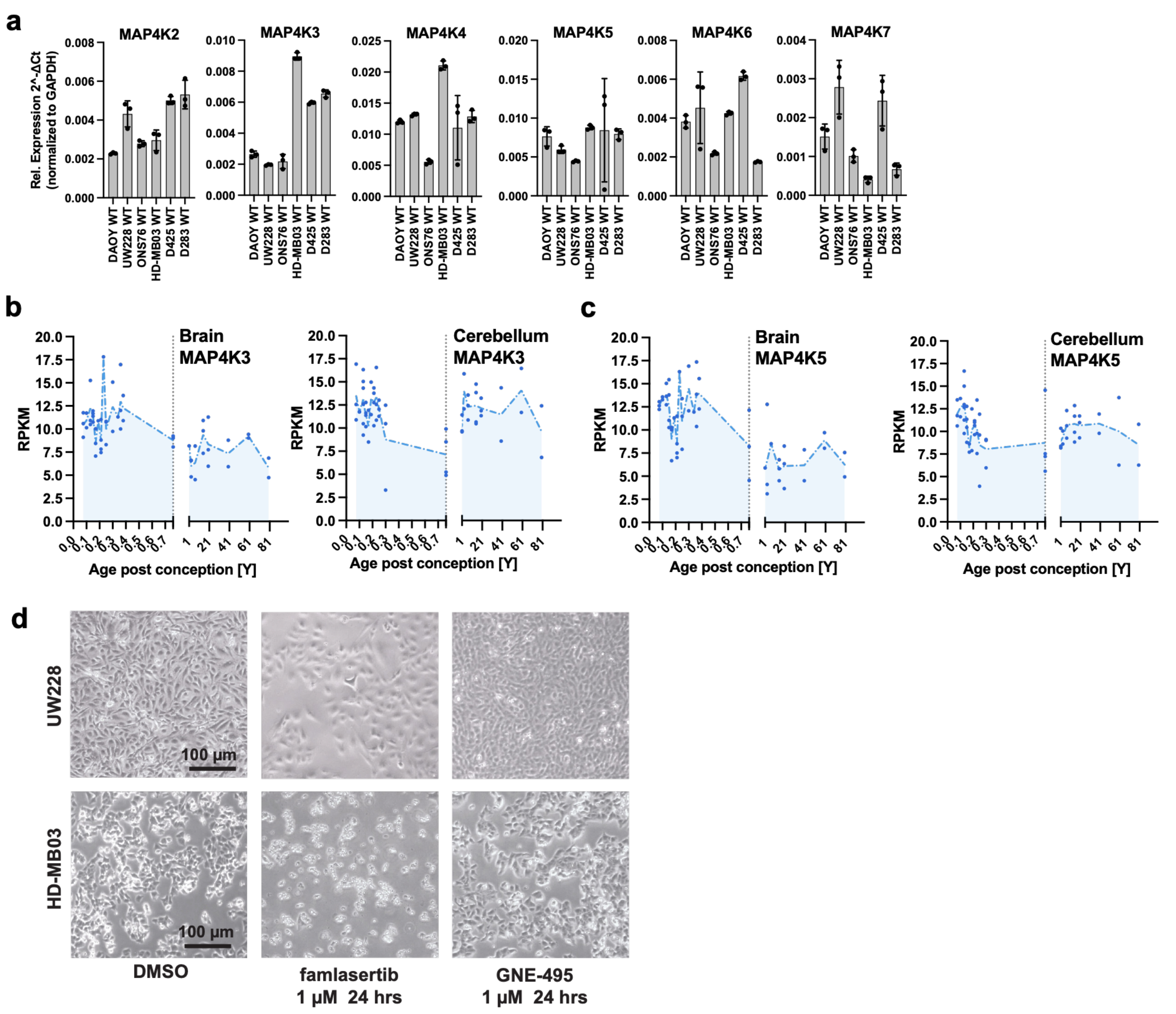


**Supplementary Figure S1**

**a)** Comparative analysis of relative MAP4K mRNA expression levels across three SHH (DAOY, UW228 and ONS-76) and three Group 3 (HD-MB03, D425 and D283) MB cell lines, determined by GAPDH-normalized qRT–PCR. **b, c)** MAP4K3 (b) and MAP4K5 (c) expression across the human lifespan in brain (left) and cerebellum (right), assessed by bulk RNA-seq. The x-axis indicates age in years; the vertical dashed line marks birth. The y-axis shows expression values as reads per kilobase per million mapped reads (RPKM). **d)** Representative bright-field images of UW228 and HD-MB03 cells after 24 h exposure to the indicated treatments.

Figure S2

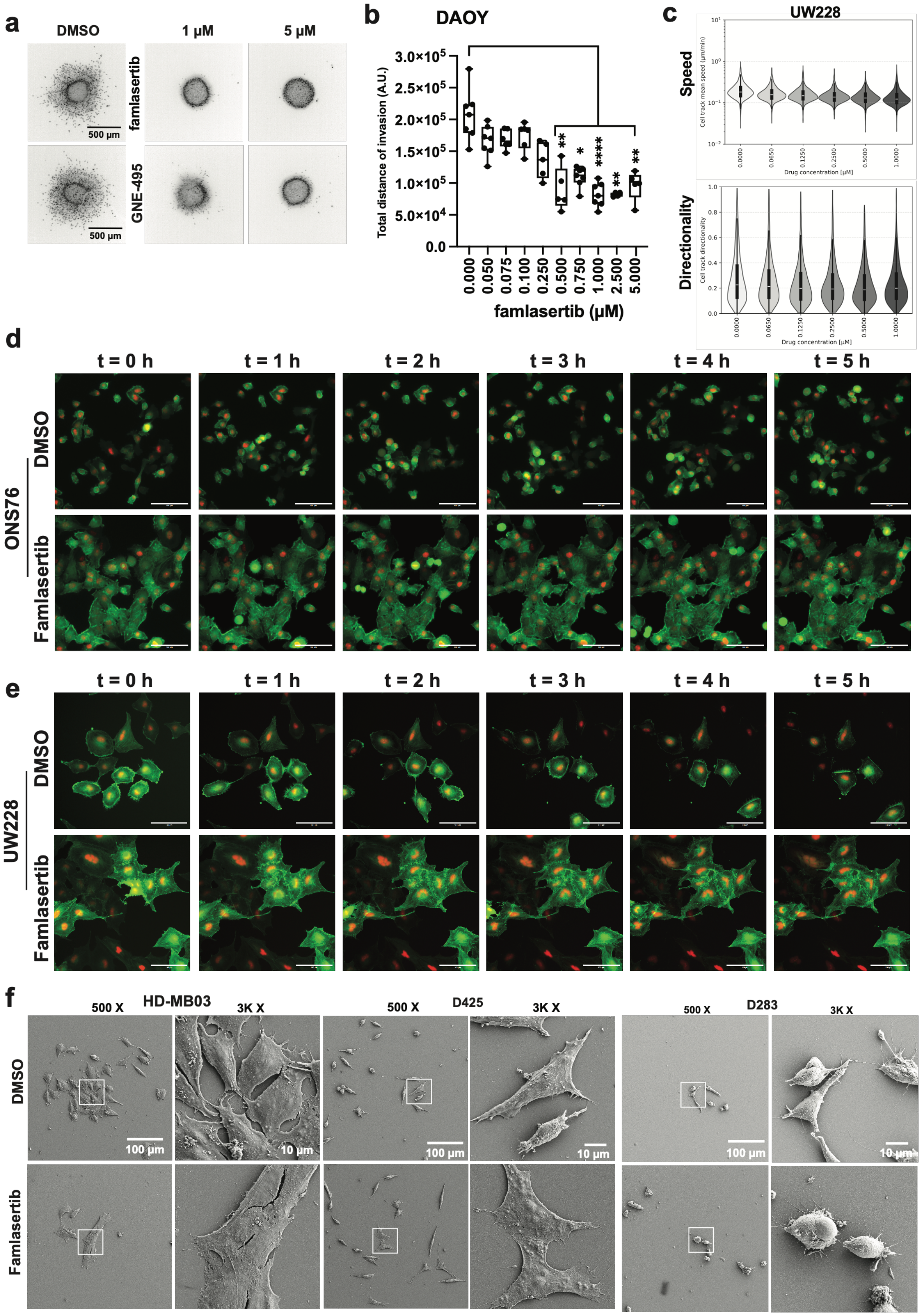
a
DMSO
1 µM
5 µM
famlasertib
GNE-495
500 µm
500 µm
b
DAOY
Total distance of invasion (A.U.)
famlasertib (µM)
c
UW228
Speed
Directionality
Drug concentration [µM]
d
t = 0 h
t = 1 h
t = 2 h
t = 3 h
t = 4 h
t = 5 h
ONS76
DMSO
Famlasertib
e
UW228
f
HD-MB03
D425
D283
500 X
3K X
100 µm
10 µm

**Supplementary Figure S2**

**a)** Representative inverted grayscale images of Hoechst-stained nuclei in collagen I–embedded spheroids at the experimental endpoint shown in (Figure 2a). **b)** Spheroid invasion assay (SIA) comparing the effects of increasing concentrations of famlasertib on bFGF-induced invasiveness in DAOY cells. DMSO served as a solvent control. Box plots show total invasion distance, calculated as the sum of invasion distances measured for each individual cell. Each dot represents one independent spheroid. **c)** Violin plots of mean track speed (f) and directionality (g) of UW228 cells treated with increasing concentrations of famlasertib. Zero concentration refers to DMSO controls. Total number of tracked objects were approximately 10000, distributed across 60 wells, with a minimum of 1250 tracks and 6 wells, and a maximum of 7000 tracks and 18 wells per condition. **d, e)** Still images of ONS-76 (c) and UW228 (d) cells expressing LA–EGFP and mCherryNuc following 24 h exposure to DMSO or 0.5 µM famlasertib. Scale bar: 100 µm. **f)** Scanning electron microscopy (SEM) images of HD-MB03, D425 and D283 cells treated with DMSO or 0.5 µM famlasertib for 24 h. White boxes indicate regions shown at 6× higher magnification to the right of each panel.

#### Figure S3

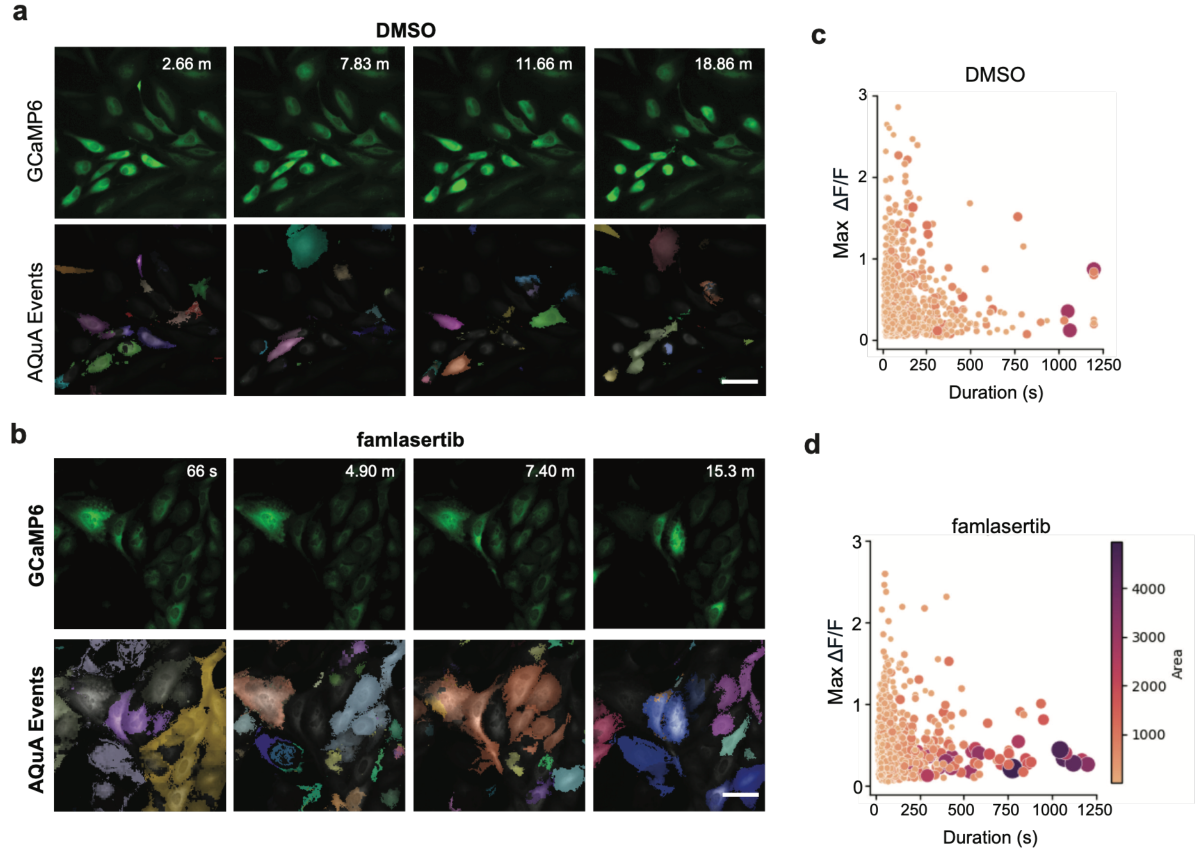


**Supplementary Figure S3**

**a, b)** Images illustrating the fluorescence change of the GCaMP6-$Ca^{2+}$ signal (top) and colour-coded, AQuA extracted calcium events (bottom) across different time points in DMSO control (a) and famlasertib (0.5 µM, (b) treated UW228 cells. **c, d)** $Ca^{2+}$ features of individual events in Max DF/F, duration and event area in DMSO control (c) and famlasertib (d) groups. Dots: individual $Ca^{2+}$ events from n = 4 of each group, respectively (Total events in DMSO control were 1435 events; in famlasertib were 1082 events).

#### Figure S4

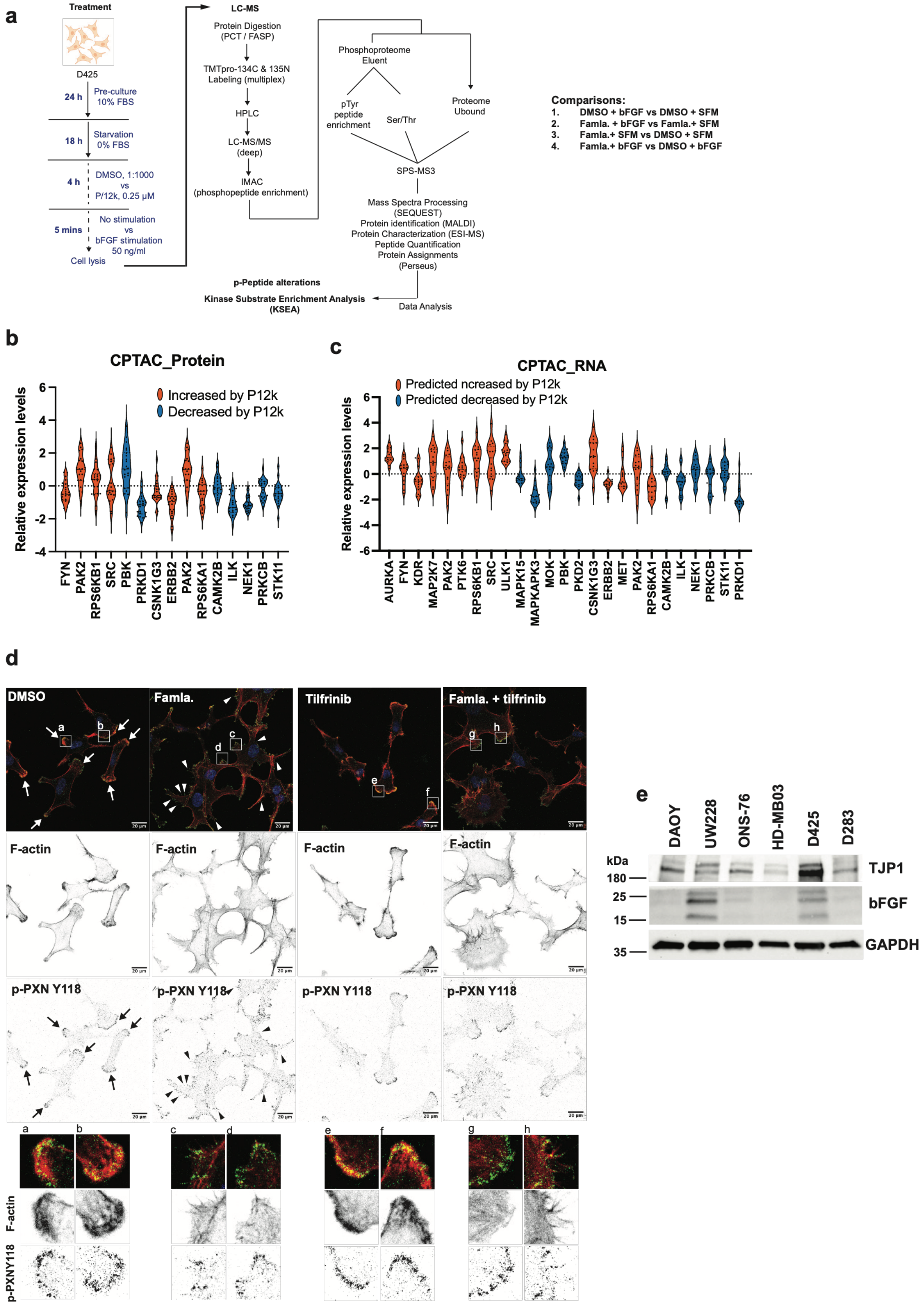

a
Treatment
D425
24 h
Pre-culture 10% FBS
18 h
Starvation 0% FBS
4 h
DMSO, 1:1000 vs P/12k, 0.25 μM
5 mins
No stimulation vs bFGF stimulation 50 ng/ml
Cell lysis
LC-MS
Protein Digestion (PCT / FASP)
TMTpro-134C & 135N Labeling (multiplex)
HPLC
LC-MS/MS (deep)
IMAC (phosphopeptide enrichment)
Phosphoproteome Eluent
pTyr peptide enrichment
Ser/Thr
Proteome Ubound
SPS-MS3
Mass Spectra Processing (SEQUEST)
Protein identification (MALDI)
Protein Characterization (ESI-MS)
Peptide Quantification
Protein Assignments (Perseus)
Data Analysis
p-Peptide alterations
Kinase Substrate Enrichment Analysis (KSEA)
Comparisons:
1. DMSO + bFGF vs DMSO + SFM
2. Famla. + bFGF vs Famla.+ SFM
3. Famla.+ SFM vs DMSO + SFM
4. Famla.+ bFGF vs DMSO + bFGF
b
CPTAC_Protein
Increased by P12k
Decreased by P12k
Relative expression levels
FYN PAK2 RPS6KB1 SRC PBK PRKD1 CSNK1G3 ERBB2 PAK2 RPS6KA1 CAMK2B ILK NEK1 PRKCB STK11
c
CPTAC_RNA
Predicted ncreased by P12k
Predicted decreased by P12k
Relative expression levels
AURKA FYN KDR MAP2K7 PAK2 PTK6 RPS6KB1 SRC ULK1 MAPK15 MAPKAPK3 MOK PBK PKD2 CSNK1G3 ERBB2 MET PAK2 RPS6KA1 CAMK2B ILK NEK1 PRKCB STK11 PRKD1
d
DMSO
Famla.
Tilfrinib
Famla. + tilfrinib
F-actin
p-PXN Y118
a b c d e f g h
p-PXNY118
e
DAOY UW228 ONS-76 HD-MB03 D425 D283
kDa
180
25
15
35
TJP1
bFGF
GAPDH

**Supplementary Figure S4**

**a)** Schematic overview of the phosphoproteomic workflow and the comparisons performed to identify treatment-dependent alterations. **b)** Correlation analysis between proteins harboring significantly altered phosphosites and corresponding protein expression scores derived from the Clinical Proteomic Tumor Analysis Consortium (CPTAC) dataset. **c)** Correlation analysis between proteins with significantly altered phosphosites and corresponding mRNA expression levels obtained from the CPTAC dataset. **d)** Confocal immunofluorescence analysis (IFA) of p-PXN (Y118) in UW228 cells treated with famlasertib, tilfrinib, or their combination. Inverted grayscale images of F-actin and p-PXN (Y118) are shown to enhance contrast. Regions a–h are displayed below the main panels at 4× magnification and highlight the distribution of p-PXN (Y118) within lamellipodia and cortical regions enriched in F-actin–positive protrusions. **e)** Immunoblot (IB) analysis of TJP1 and bFGF expression across different MB cell models, with GAPDH serving as a loading control.

### Figure S5

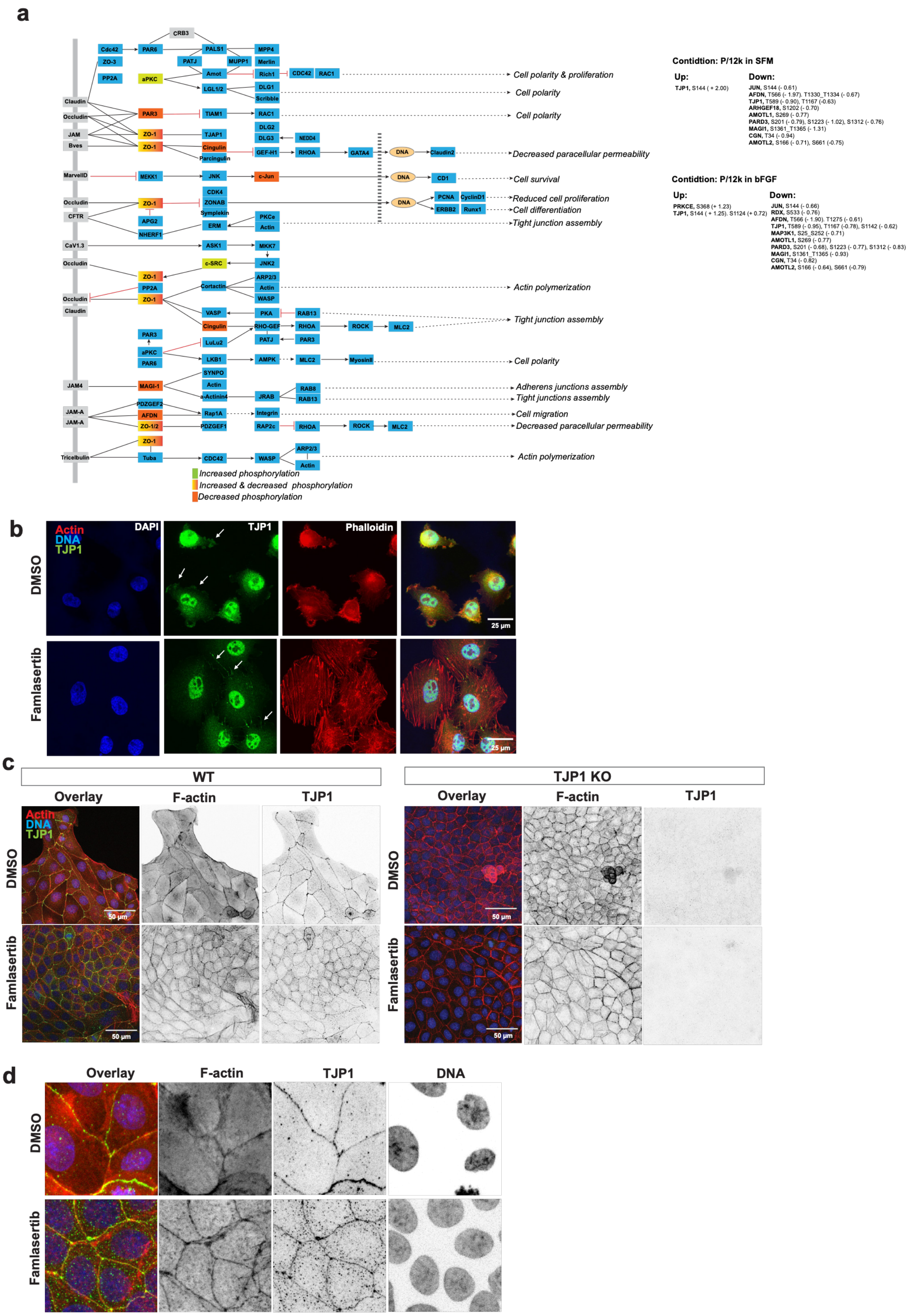
a
CRB3
Cdc42
PAR6
PALS1
MPP4
ZO-3
PATJ
MUPP1
Merlin
PP2A
aPKC
Amot
Rich1
CDC42
RAC1
Cell polarity & proliferation
LGL1/2
DLG1
Scribble
Cell polarity
Claudin
Occludin
PAR3
TIAM1
RAC1
Cell polarity
JAM
ZO-1
TJAP1
DLG2
DLG3
NEDD4
Bves
ZO-1
Cingulin
GEF-H1
RHOA
GATA4
DNA
Claudin2
Decreased paracellular permeability
Parcingulin
MarvelD
MEKK1
JNK
c-Jun
DNA
CD1
Cell survival
CDK4
Occludin
ZO-1
ZONAB
DNA
PCNA
CyclinD1
Reduced cell proliferation
CFTR
Symplekin
ERBB2
Runx1
Cell differentiation
APG2
PKCe
ERM
Actin
Tight junction assembly
NHERF1
CaV1.3
ASK1
MKK7
Occludin
c-SRC
JNK2
ZO-1
ARP2/3
PP2A
Cortactin
Actin
Actin polymerization
Occludin
ZO-1
WASP
Claudin
VASP
PKA
RAB13
Tight junction assembly
Cingulin
RHO-GEF
RHOA
ROCK
MLC2
PATJ
PAR3
PAR3
aPKC
LuLu2
PAR6
LKB1
AMPK
MLC2
Myosinll
Cell polarity
SYNPO
JAM4
MAGI-1
Actin
RAB8
Adherens junctions assembly
a-Actinin4
JRAB
RAB13
Tight junctions assembly
PDZGEF2
JAM-A
AFDN
Rap1A
Integrin
Cell migration
JAM-A
ZO-1/2
PDZGEF1
RAP2c
RHOA
ROCK
MLC2
Decreased paracellular permeability
ZO-1
ARP2/3
Tricelbulin
Tuba
CDC42
WASP
Actin
Actin polymerization
Increased phosphorylation
Increased & decreased phosphorylation
Decreased phosphorylation
Contidtion: P/12k in SFM
Up:
TJP1, S144 ( + 2.00)
Down:
JUN, S144 (- 0.61)
AFDN, T566 (- 1.97). T1330_T1334 (- 0.67)
TJP1, T589 (- 0.90), T1167 (-0.63)
ARHGEF18, S1202 (- 0.70)
AMOTL1, S269 (- 0.77)
PARD3, S201 (- 0.79), S1223 (- 1.02), S1312 (- 0.76)
MAGI1, S1361_T1365 (- 1.31)
CGN, T34 (- 0.94)
AMOTL2, S166 (- 0.71), S661 (-0.75)
Contidtion: P/12k in bFGF
Up:
PRKCE, S368 (+ 1.23)
TJP1, S144 ( + 1.25). S1124 (+ 0.72)
Down:
JUN, S144 (- 0.66)
RDX, S533 (- 0.76)
AFDN, T566 (- 1.90). T1275 (- 0.61)
TJP1, T589 (- 0.95), T1167 (-0.78), S1142 (- 0.62)
MAP3K1, S25_S252 (- 0.71)
AMOTL1, S269 (- 0.77)
PARD3, S201 (- 0.68), S1223 (- 0.77), S1312 (- 0.83)
MAGI1, S1361_T1365 (- 0.93)
CGN, T34 (- 0.82)
AMOTL2, S166 (- 0.64), S661 (-0.79)
b
Actin
DNA
TJP1
DAPI
TJP1
Phalloidin
DMSO
Famlasertib
25 μm
25 μm
c
WT
TJP1 KO
Overlay
F-actin
TJP1
Overlay
F-actin
TJP1
Actin
DNA
TJP1
DMSO
Famlasertib
50 μm
50 μm
50 μm
50 μm
d
Overlay
F-actin
TJP1
DNA
DMSO
Famlasertib

**Supplementary Figure S5**

**a)** Schematic overview of integral membrane proteins of cell junctions (grey boxes), junction-associated proteins, and components of connected signaling pathways (blue boxes). Junction-associated proteins exhibiting increased or decreased phosphorylation upon famlasertib treatment are color coded. Significantly altered phosphosites in junction-associated proteins are shown on the right. **b)** Maximum intensity projections of confocal IFA of ONS-76 cells treated with DMSO or 1 µM famlasertib for 24 h. **c)** Maximum intensity projections of confocal IFA of MDCK wild-type (WT) and MDCK TJP1 knockout (KO) cells treated with DMSO or 1 µM famlasertib for 24 h. **d)** Maximum intensity projections of confocal IFA of MDCK wild-type (WT) cells treated with DMSO or 1 µM famlasertib for 24 h.

#### Figure S6

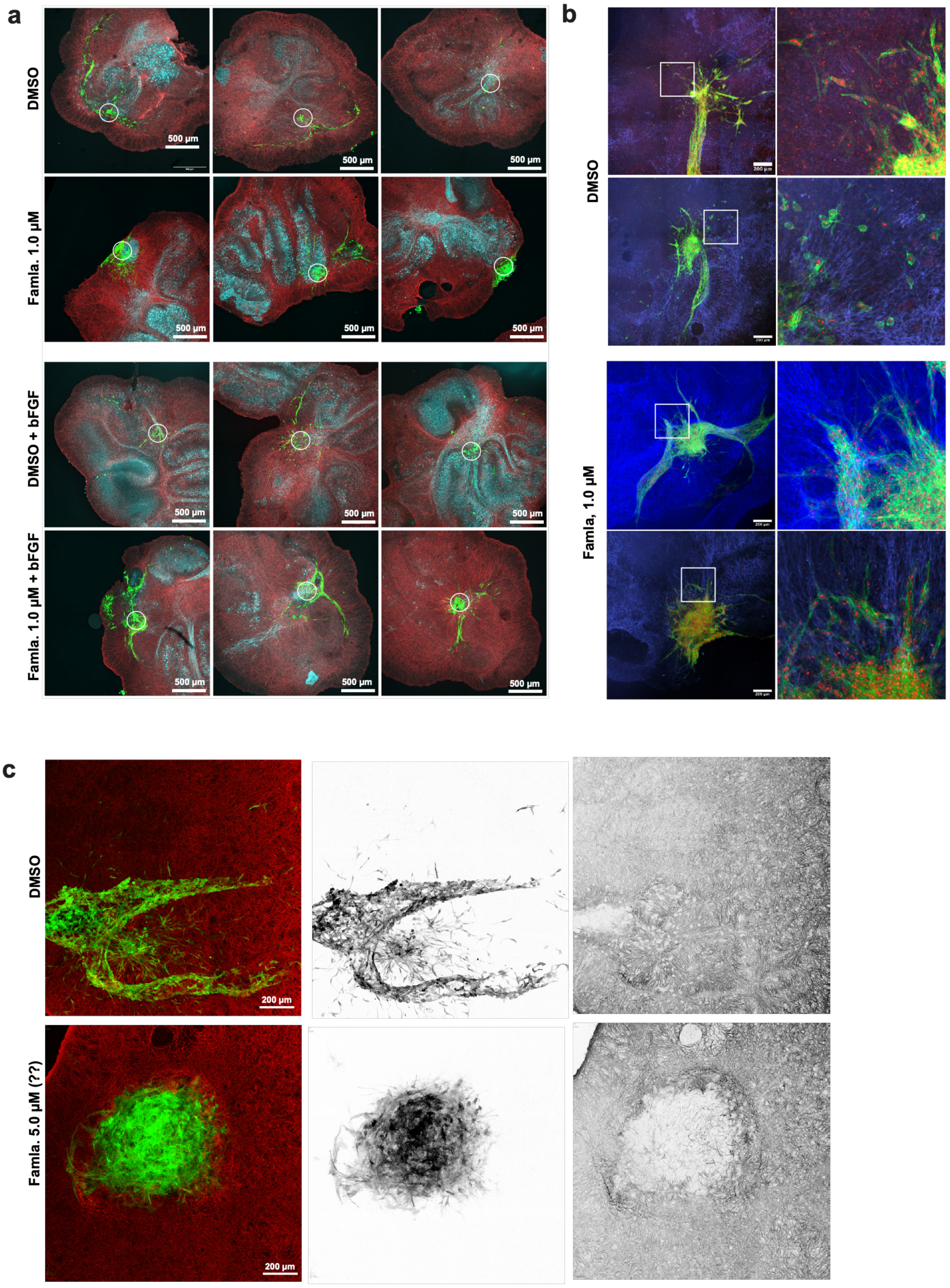


**Supplementary Figure S6**

**a)** Maximum intensity projections of confocal IFA of organotypic cerebellar slice cultures implanted with ONS-76 spheroids and treated for 5 days with famlasertib (1 µM), -/+ bFGF (50 ng/ml). Green: LA-EFP, cyan: calbindin (Purkinje cells), red: GFAP (Glial cells. **b)** Maximum intensity projections of confocal IFA of organotypic cerebellar slice cultures implanted with ONS-76 spheroids and treated for 5 days with famlasertib (1 µM). Green: LA-EFP, Blue: GFAP (Glial cells), red: human nuclei. **c)** Maximum intensity projections of confocal IFA of organotypic cerebellar slice cultures implanted with ONS-76 spheroids and treated for 5 days with famlasertib (5 μM). Inverted grayscale images of LA–EGFP (tumor cells) and GFAP (glial cells within the tumor microenvironment) are shown to enhance contrast. Green: LA–EGFP; red: GFAP.

**Figure S7**

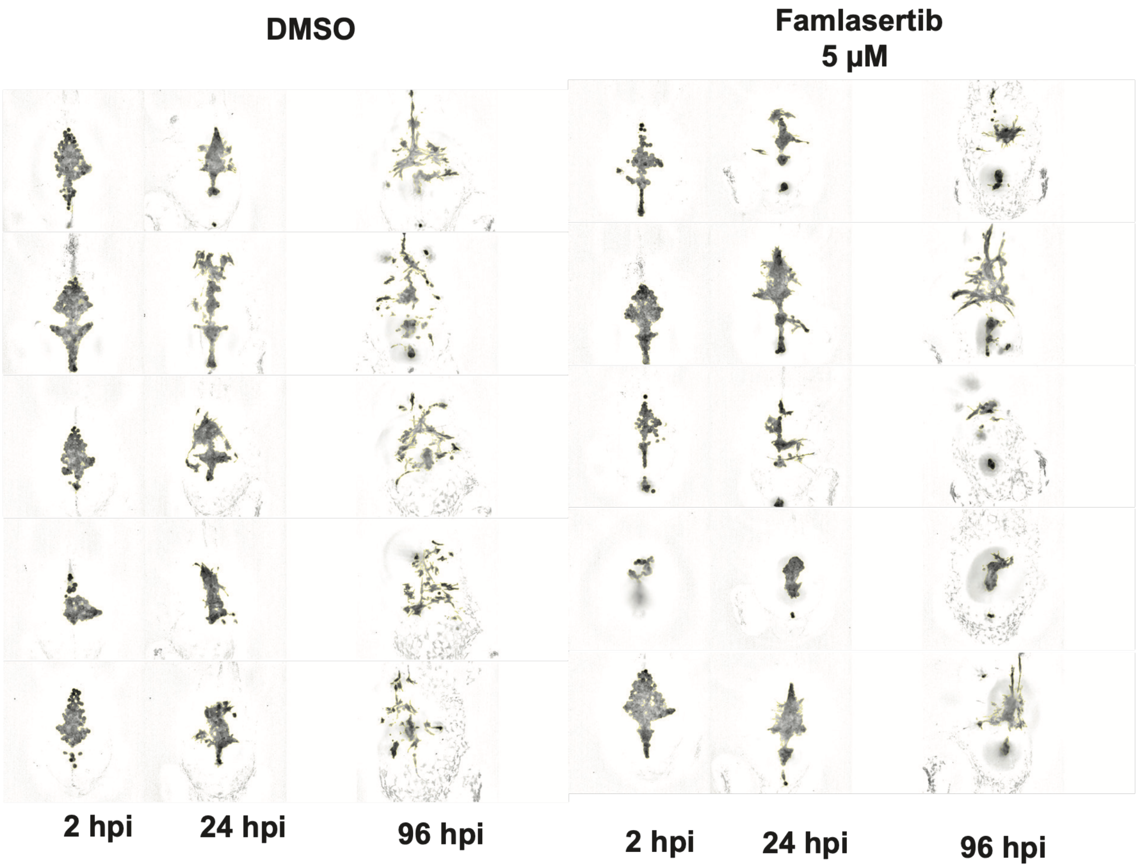


**Supplementary Figure S7**

Grayscale fluorescence imaging with tumor segmentation highlighting tumor progression. Representative live images of LA–EGFP signal at 2, 24, and 96 hpi in larvae treated with DMSO (n = 5) or famlasertib (n = 5). Tumor segmentation shown with a yellow line.
