## Supplementary table 1 for "MAP4K inhibition by the CNS penetrant inhibitor famlasertib restrains medulloblastoma dissemination without developmental toxicity"

| Master_id_first | PsiteInProtein | diff | FDR |
|---|---|---|---|
| P84098 | S189 | -1.601285935 | 5.34E-04 |
| O15318 | S157 | -1.099746365 | 1.48E-06 |
| Q9NRR4 | S217_S221 | -1.084820763 | 0.053353531 |
| P11388 | S1377 | -0.906680414 | 4.94E-04 |
| Q9UHB7 | SSPRPTAEK | -0.790739696 | 0.008201779 |
| Q9UET6 | S328 | -0.787862448 | 0.020411503 |
| Q96RK0 | S436 | -0.725815166 | 0.03996628 |
| Q9UQ35 | S222 | -0.647954614 | 9.28E-04 |
| Q9UGJ0 | S71 | -0.623054246 | 1.05E-04 |
| Q12830 | S1310 | -0.61198994 | 1.64E-04 |
| O75152 | S758 | 0.600581432 | 1.08E-04 |
| Q9NW75 | S129 | 0.603346356 | 0.025396568 |
| O95239 | S801 | 0.608402004 | 7.23E-04 |
| Q9BW71 | S269 | 0.612583742 | 0.015086008 |
| P23588 | S230 | 0.617812509 | 0.002774465 |
| Q2LD37 | S4124 | 0.620432603 | 0.002919652 |
| Q96RK0 | S173 | 0.621975348 | 0.00325839 |
| Q07352 | S92 | 0.622425044 | 0.001706091 |
| P47974 | S123 | 0.625429989 | 5.13E-04 |
| Q5T3J3 | T389 | 0.626906056 | 8.35E-04 |
| Q9C0B5 | S585 | 0.630533702 | 1.62E-05 |
| Q14980 | S2047 | 0.631751766 | 0.030072139 |
| P50548 | T526 | 0.632349172 | 6.68E-05 |
| O14686 | S2726 | 0.634112153 | 0.003166261 |
| Q86SQ0 | S212 | 0.634453806 | 0.025782168 |
| O43765 | S305 | 0.636390615 | 7.41E-04 |
| Q14687 | S140 | 0.647659414 | 0.042091275 |
| P29966 | T143_S/T | 0.650991426 | 0.005211092 |
| Q7Z333 | S1366 | 0.654912206 | 0.002210316 |
| Q9Y618 | T1891 | 0.655328979 | 5.93E-05 |
| O94854 | S578 | 0.662045104 | 7.44E-04 |
| Q15691 | S155 | 0.667176558 | 1.60E-05 |
| Q5TCQ9 | S225 | 0.668281716 | 0.003311399 |
| Q9UPP1 | S1021 | 0.671320468 | 0.003214663 |
| P15822 | T815 | 0.677893992 | 0.032612101 |
| Q9HCH0 | S1295 | 0.678682301 | 0.017665654 |
| Q9NZJ0 | S557 | 0.680741726 | 0.00131733 |
| Q9UKA4 | S433 | 0.687767512 | 0.01083684 |
| Q9Y6Y0 | T330 | 0.688945771 | 7.36E-06 |
| Q9NPR2 | S830 | 0.690385117 | 2.79E-05 |
| Q09666 | S5110 | 0.691529262 | 0.007446374 |
| Q6PKG0 | T376 | 0.693030694 | 1.05E-05 |
| O43395 | T172 | 0.706961962 | 9.69E-04 |
| Q8TF72 | S285 | 0.715513557 | 5.41E-04 |
| Q9UBH6 | S676 | 0.731256273 | 6.05E-04 |
| A6H8Y1 | S100 | 0.742657482 | 8.49E-04 |
| Q9UGU5 | S197 | 0.742826356 | 2.09E-04 |
| P0CAP2 | S179 | 0.744699706 | 0.04100589 |
| P55196 | T1352 | 0.744821371 | 1.62E-05 |
| Q9C0A6 | RSSQAGDIAAEK | 0.747945513 | 0.011160465 |
| Q86SQ0 | KSSISSISGRDDLMDYHR | 0.755708695 | 0.001935992 |
| Q5SW79 | S1160_S1165 | 0.760147348 | 3.02E-06 |
| Q5VZL5 | S1181 | 0.760613334 | 0.00649768 |
| P20794 | Y159 | 0.76152212 | 0.0092259 |

| | | | |
|---|---|---|---|
| Q9C0B5 | S409 | 0.764737033 | 4.32E-06 |
| P15056 | S429 | 0.765237995 | 2.56E-04 |
| P19634 | S703 | 0.765531499 | 0.003366383 |
| Q8IY17 | S372 | 0.768346763 | 0.001729054 |
| P30260 | S364 | 0.77178388 | 9.03E-04 |
| P51812 | S369_S375 | 0.77284535 | 7.10E-06 |
| O76039 | S407 | 0.784799619 | 1.06E-06 |
| Q5T5P2 | S474 | 0.805447723 | 9.92E-06 |
| Q765P7 | S601 | 0.814822492 | 1.78E-06 |
| Q6EEV4 | LRAPSSR | 0.816925808 | 7.16E-04 |
| P49327 | S831 | 0.830290237 | 0.010818611 |
| Q07889 | S1134 | 0.831732501 | 8.35E-05 |
| Q2LD37 | S3653 | 0.844740204 | 2.72E-05 |
| Q02880 | S1466_S1476 | 0.860430005 | 0.012477168 |
| Q9Y6J0 | T1927 | 0.865531602 | 0.001441422 |
| Q14160 | T1342_S1348 | 0.873953214 | 5.69E-04 |
| P09884 | T219 | 0.88199901 | 2.99E-07 |
| P10398 | T181 | 0.882434715 | 2.01E-05 |
| O94804 | T952 | 0.904594386 | 2.13E-05 |
| Q7L804 | S388 | 0.905983072 | 1.61E-05 |
| O60934 | T497 | 0.90645826 | 2.06E-04 |
| O14924 | S172 | 0.907228951 | 4.45E-04 |
| P47974 | S125 | 0.908337559 | 8.78E-04 |
| Q14934 | S272 | 0.927758926 | 0.001247339 |
| O15357 | T958 | 0.92990858 | 1.88E-05 |
| P78536 | S819 | 0.937365009 | 0.04174181 |
| Q9BY44 | S526 | 0.94387581 | 2.47E-04 |
| Q5VT52 | S581 | 0.950257319 | 2.67E-04 |
| P35658 | S430_S433 | 0.967176553 | 4.29E-06 |
| O43379 | S49 | 0.978846214 | 2.35E-05 |
| Q5TKA1 | S65 | 0.987942275 | 2.09E-04 |
| O75151 | S1056_S/T | 0.99804248 | 0.00353474 |
| Q8N5U6 | S128 | 1.001813291 | 5.35E-07 |
| Q32P44 | S156_S | 1.022239865 | 0.017095907 |
| Q9BY84 | S501 | 1.029517509 | 0.036189437 |
| Q8NG27 | S265 | 1.045478351 | 9.74E-05 |
| P41162 | TESSPGSR | 1.056013312 | 0.001798125 |
| Q9UKJ3 | T293 | 1.060143047 | 2.70E-07 |
| P53367 | T361 | 1.077303313 | 9.25E-06 |
| Q8TF72 | S1173 | 1.083112256 | 5.18E-06 |
| O43295 | S858 | 1.087697647 | 1.45E-04 |
| Q6ZSZ5 | T1345 | 1.090282824 | 1.99E-08 |
| Q92609 | S558_T/S | 1.091240871 | 6.04E-06 |
| Q6Y7W6 | S26_S30 | 1.110548473 | 1.99E-08 |
| P23443 | T444 | 1.111601402 | 9.88E-07 |
| Q9UHI6 | S500 | 1.113680257 | 1.99E-06 |
| Q9NUL3 | S440 | 1.114938608 | 6.24E-06 |
| Q6ZRI6 | T364 | 1.118541397 | 4.77E-07 |
| P78536 | T735 | 1.122244157 | 6.92E-09 |
| Q8N5C8 | S645 | 1.140803328 | 5.37E-04 |
| O14974 | S507 | 1.145085025 | 5.42E-06 |
| Q86SQ0 | S415 | 1.153948522 | 4.25E-05 |
| Q15418 | S363_S369 | 1.158902881 | 2.70E-07 |
| Q9HCM7 | S487 | 1.15983313 | 5.65E-05 |
| Q7RTP6 | RKTSQSEEEEAPR | 1.160156936 | 8.13E-06 |
| Q92995 | S40 | 1.171179592 | 5.40E-05 |

| | | | |
|---|---|---|---|
| Q15418 | S732 | 1.174621426 | 2.06E-04 |
| Q8TF72 | S1279 | 1.1846569 | 1.08E-04 |
| P30307 | T48 | 1.211040686 | 2.99E-07 |
| Q9UJY4 | S400 | 1.22029939 | 9.03E-04 |
| Q14680 | T428 | 1.282646332 | 5.06E-08 |
| Q8TF72 | S816 | 1.341870967 | 2.70E-07 |
| Q15418 | S380 | 1.341896167 | 5.35E-07 |
| P15056 | S151 | 1.355292094 | 2.00E-07 |
| Q8IWC1 | S524 | 1.362640637 | 2.70E-07 |
| Q15154 | S1283 | 1.378095071 | 3.91E-05 |
| P12270 | S2155 | 1.426634142 | 3.58E-08 |
| Q7Z5J4 | T1068 | 1.433040868 | 1.88E-07 |
| Q5SW79 | S1160 | 1.495906705 | 6.04E-08 |
| P17544 | T326 | 1.513289333 | 6.89E-04 |
| Q8TF72 | S403 | 1.543404534 | 7.41E-07 |
| Q9UQC2 | S543 | 1.573349361 | 1.13E-07 |
| P12270 | T2116 | 1.586348409 | 1.28E-07 |
| P27361 | Y204 | 1.676090111 | 2.70E-07 |
| P28482 | T185 | 1.695007092 | 9.67E-08 |
| Q9UJY4 | S177 | 1.816177669 | 1.62E-05 |
| Q9UDY2 | S163 | 1.88550839 | 9.88E-07 |
| Q9NZN5 | T703 | 2.151653105 | 0.00171603 |
| P28482 | Y187 | 2.434053769 | 5.06E-08 |
| P27361 | T202_Y204 | 2.990318021 | 1.86E-09 |
| P28482 | T185_Y187 | 3.267982367 | 1.86E-09 |
