## Supplementary table 2 for "MAP4K inhibition by the CNS penetrant inhibitor famlasertib restrains medulloblastoma dissemination without developmental toxicity"

| Master_id_first | PsiteInProtein | diff | FDR |
|---|---|---|---|
| P84098 | S189 | -1.670489281 | 2.81E-04 |
| Q9NRR4 | S217_S221 | -1.123886299 | 0.042133206 |
| O15318 | S157 | -1.011674643 | 3.33E-06 |
| Q96RK0 | S436 | -0.955248207 | 0.005313429 |
| P11388 | S1377 | -0.890785582 | 4.64E-04 |
| Q9UHB7 | SSPRPTAEK | -0.744292381 | 0.012496476 |
| Q86YS7 | S295 | -0.724145059 | 2.27E-04 |
| Q12830 | S1310 | -0.720303255 | 2.75E-05 |
| Q9UQ35 | S222 | -0.714191545 | 3.12E-04 |
| Q58WW2 | S847_S850 | -0.710412426 | 0.045456295 |
| P55327 | S36 | -0.702963265 | 6.88E-05 |
| O95359 | S2317 | -0.650783117 | 5.09E-04 |
| Q9UGJ0 | S71 | -0.644264742 | 6.58E-05 |
| Q9UPW6 | S39 | -0.64230502 | 0.006363026 |
| Q96T58 | T1619 | -0.628494198 | 0.048388526 |
| Q5VZ89 | S1225 | 0.601622749 | 0.001690327 |
| A6H8Y1 | S100 | 0.605046658 | 0.004766805 |
| O43765 | S305 | 0.605124332 | 9.75E-04 |
| P50747 | S79 | 0.6067396 | 0.003477569 |
| Q9UBH6 | S676 | 0.612103292 | 0.002709452 |
| Q8NFC6 | S1077 | 0.612352517 | 1.69E-05 |
| Q9BW71 | S269 | 0.616715941 | 0.013969911 |
| P20794 | Y159 | 0.621540647 | 0.035205852 |
| Q8IWN7 | S1671 | 0.621928479 | 0.004060354 |
| Q16644 | T201 | 0.636610904 | 0.001132634 |
| Q9NW75 | S129 | 0.638526346 | 0.016179326 |
| P15056 | S429 | 0.642108443 | 0.001173429 |
| O43395 | T172 | 0.64835402 | 0.001919793 |
| P50548 | T526 | 0.64887295 | 4.81E-05 |
| Q2NKX8 | FDASTPKNDISPP | 0.649344065 | 9.14E-04 |
| Q8NDI1 | S649 | 0.656249797 | 2.60E-04 |
| Q86VW2 | S556_T565_S/T | 0.658502784 | 7.94E-04 |
| Q96D71 | S709 | 0.658641067 | 1.08E-04 |
| Q9Y3R0 | S996 | 0.660215311 | 0.003485813 |
| O75152 | S758 | 0.672977667 | 3.11E-05 |
| Q7LBC6 | T1270 | 0.673014802 | 4.55E-04 |
| Q5T3J3 | T389 | 0.677200574 | 3.26E-04 |
| Q8TF72 | S285 | 0.690682971 | 6.32E-04 |
| Q7Z4S6 | S853 | 0.691495771 | 1.70E-04 |
| P55196 | T1352 | 0.694573587 | 3.27E-05 |
| O75151 | S1056_S/T | 0.696891464 | 0.043890913 |
| Q6PKG0 | T376 | 0.697518943 | 8.85E-06 |
| Q2LD37 | S4124 | 0.698401708 | 8.41E-04 |
| Q5T5P2 | S474 | 0.699718392 | 4.38E-05 |
| Q8IY17 | S372 | 0.700691502 | 0.003477569 |
| Q8IWB9 | S270_S/T | 0.706474401 | 0.008155201 |
| P23588 | S230 | 0.70734361 | 6.87E-04 |
| Q9NZJ0 | S557 | 0.713796214 | 7.21E-04 |
| Q5U5Q3 | S537_S/T | 0.714020432 | 0.0123582 |
| Q9P266 | S1194 | 0.714525412 | 1.13E-04 |
| Q9Y6Y0 | T330 | 0.719599582 | 3.82E-06 |
| O94854 | S578 | 0.727203002 | 2.54E-04 |
| O75151 | S625 | 0.727555284 | 0.017760793 |
| Q9HCH0 | S1295 | 0.736508525 | 0.009349702 |

| | | | |
|---|---|---|---|
| Q9P2B2 | S875 | 0.740392655 | 0.037535384 |
| Q5SW79 | S1160_S1165 | 0.743910193 | 3.32E-06 |
| Q15691 | S155 | 0.745540061 | 3.82E-06 |
| Q96RK0 | S173 | 0.74998776 | 5.09E-04 |
| Q9UGU5 | S197 | 0.759878196 | 1.55E-04 |
| Q9C0B5 | S409 | 0.774603299 | 3.25E-06 |
| Q5SW79 | T174 | 0.776978059 | 0.019969504 |
| Q86SQ0 | SISSISGRDDLMDY | 0.778628242 | 0.001297773 |
| Q6EEV4 | LRAPSSR | 0.780686954 | 8.99E-04 |
| Q9UKA4 | S433 | 0.794625722 | 0.003326958 |
| Q8WXD9 | S826 | 0.798325002 | 5.05E-05 |
| O43379 | S49 | 0.803109157 | 1.84E-04 |
| Q9BY44 | S526 | 0.808955982 | 9.14E-04 |
| Q07889 | S1134 | 0.811811249 | 9.68E-05 |
| Q9UPP1 | S1021 | 0.820420955 | 4.39E-04 |
| Q9H6F5 | S217 | 0.821211657 | 0.002724479 |
| Q9Y6X9 | S739_S743 | 0.825922634 | 0.006955041 |
| P0CAP2 | S179 | 0.829747668 | 0.019228924 |
| P09884 | T219 | 0.836571736 | 5.24E-07 |
| O14686 | S2726 | 0.863333634 | 1.55E-04 |
| Q9UPN7 | S726 | 0.867783888 | 0.006027765 |
| Q2LD37 | S3653 | 0.884869812 | 1.54E-05 |
| P51812 | S369_S375 | 0.887843423 | 1.33E-06 |
| O60934 | T497 | 0.892160715 | 2.16E-04 |
| Q14934 | S272 | 0.892917386 | 0.001536624 |
| Q765P7 | S601 | 0.900876186 | 5.38E-07 |
| Q14160 | T1342_S1348 | 0.930576518 | 2.55E-04 |
| O15357 | T958 | 0.933498436 | 1.68E-05 |
| P10398 | T181 | 0.955592066 | 7.81E-06 |
| Q5TKA1 | S65 | 0.961395971 | 2.39E-04 |
| Q86SQ0 | S212 | 0.961723677 | 7.87E-04 |
| Q5VT52 | S581 | 0.965151101 | 2.09E-04 |
| Q9UKJ3 | T293 | 0.969989181 | 6.66E-07 |
| Q8N5U6 | S128 | 0.974062795 | 6.66E-07 |
| O76039 | S407 | 1.001394554 | 5.85E-08 |
| P41162 | TESSPGSR | 1.003331325 | 0.002537304 |
| P47974 | S125 | 1.020107506 | 2.47E-04 |
| Q8TF72 | S1173 | 1.02690173 | 7.86E-06 |
| P53367 | T361 | 1.030749738 | 1.37E-05 |
| Q8NG27 | S265 | 1.040476507 | 9.36E-05 |
| P30260 | S364 | 1.048750617 | 3.85E-05 |
| Q8N5C8 | S645 | 1.049056194 | 9.50E-04 |
| Q7L804 | S388 | 1.090364347 | 1.88E-06 |
| O94804 | T952 | 1.091401297 | 2.53E-06 |
| Q9UHI6 | S500 | 1.101983264 | 2.09E-06 |
| O14924 | S172 | 1.113490681 | 4.74E-05 |
| P23443 | T444 | 1.11543865 | 8.48E-07 |
| P35658 | S430_S433 | 1.123403476 | 6.66E-07 |
| Q02880 | S1466_S1476 | 1.131573865 | 0.001148546 |
| Q6ZRI6 | T364 | 1.151053588 | 3.20E-07 |
| P78536 | T735 | 1.151662174 | 4.76E-09 |
| Q9HCM7 | S487 | 1.155552962 | 5.32E-05 |
| Q09666 | S5110 | 1.161055922 | 5.05E-05 |
| Q6ZSZ5 | T1345 | 1.18162819 | 5.50E-09 |
| Q9NUL3 | S440 | 1.199038899 | 2.53E-06 |
| O14974 | S507 | 1.199310364 | 2.83E-06 |

| | | | |
|---|---|---|---|
| Q14680 | T428 | 1.208454976 | 5.85E-08 |
| Q7RTP6 | RKTSQSEEEEAPI | 1.209598439 | 4.24E-06 |
| Q9Y6J0 | T1927 | 1.213465802 | 4.64E-05 |
| P30307 | T48 | 1.229095945 | 2.56E-07 |
| Q6Y7W6 | S26_S30 | 1.241505956 | 5.01E-09 |
| Q92995 | S40 | 1.24511433 | 2.72E-05 |
| Q8TF72 | S1279 | 1.270024865 | 4.88E-05 |
| Q92609 | S558_T/S | 1.294311834 | 7.57E-07 |
| Q86SQ0 | S415 | 1.311617773 | 9.80E-06 |
| Q15154 | S1283 | 1.320951525 | 5.59E-05 |
| Q8IWC1 | S524 | 1.337802706 | 3.14E-07 |
| P49327 | S831 | 1.362757505 | 1.14E-04 |
| Q15418 | S363_S369 | 1.387172446 | 3.23E-08 |
| Q9UJY4 | S400 | 1.396751421 | 2.24E-04 |
| Q5SW79 | S1160 | 1.419707148 | 6.33E-08 |
| Q8TF72 | S403 | 1.475842451 | 1.11E-06 |
| O43295 | S858 | 1.484343041 | 4.24E-06 |
| Q15418 | S732 | 1.528840505 | 1.08E-05 |
| Q8TF72 | S816 | 1.53438654 | 5.69E-08 |
| Q9UQC2 | S543 | 1.544255432 | 8.54E-08 |
| Q7Z5J4 | T1068 | 1.551286529 | 5.69E-08 |
| P15056 | S151 | 1.558659437 | 3.23E-08 |
| P12270 | S2155 | 1.581266963 | 7.02E-09 |
| Q15418 | S380 | 1.622483945 | 5.85E-08 |
| P28482 | T185 | 1.908205633 | 1.99E-08 |
| P12270 | T2116 | 1.935996532 | 1.13E-08 |
| P27361 | Y204 | 2.028335793 | 3.23E-08 |
| Q9UJY4 | S177 | 2.051775366 | 3.67E-06 |
| Q9UDY2 | S163 | 2.053682259 | 3.52E-07 |
| Q9NZN5 | T703 | 2.496942393 | 3.48E-04 |
| P28482 | Y187 | 2.826457168 | 7.02E-09 |
| P27361 | T202_Y204 | 3.452269656 | 2.60E-10 |
| P28482 | T185_Y187 | 3.653298586 | 2.60E-10 |
