## Supplementary table 3 for "MAP4K inhibition by the CNS penetrant inhibitor famlasertib restrains medulloblastoma dissemination without developmental toxicity"

| Master_id_first | PsiteInProtein | diff | FDR |
|---|---|---|---|
| Q14667 | T1269 | -3.108141116 | 3.64E-04 |
| Q12986 | NYSSPPPCHLSR | -2.817871942 | 0.005560233 |
| Q8TF76 | S147 | -2.481805893 | 4.08E-04 |
| Q6GQQ9 | T547 | -2.066624751 | 8.80E-10 |
| Q9ULF5 | T540 | -2.012049272 | 1.68E-08 |
| P55196 | T566 | -1.904127051 | 6.70E-06 |
| Q96B97 | T354 | -1.864841922 | 3.65E-06 |
| P55010 | PPENSDSGTGKK | -1.724806875 | 1.05E-08 |
| Q9Y490 | T144 | -1.708470042 | 4.50E-08 |
| Q14008 | T569 | -1.701629405 | 6.30E-08 |
| O60716 | T916 | -1.564088716 | 3.96E-07 |
| Q96PY5 | T467 | -1.504990335 | 2.55E-08 |
| O60716 | T177 | -1.497759427 | 1.31E-07 |
| Q9HBI1 | T28 | -1.449898751 | 6.70E-06 |
| Q9HAU0 | S855_T868 | -1.448978882 | 1.05E-08 |
| Q96PY5 | T450 | -1.439540452 | 3.99E-06 |
| Q5VWQ0 | S92 | -1.401106769 | 0.005932397 |
| Q9UBT7 | S538 | -1.388417033 | 2.33E-08 |
| P55010 | T153 | -1.37072482 | 1.68E-08 |
| Q7L4E1 | S276 | -1.358098142 | 0.002070713 |
| O15355 | S527 | -1.320544943 | 1.05E-08 |
| Q8NHQ8 | PFQSGSLKRPGS | -1.299050452 | 2.26E-08 |
| Q9HAU0 | T868 | -1.279379983 | 1.68E-08 |
| Q5HYJ3 | S227 | -1.247831063 | 2.98E-06 |
| Q6WCQ1 | S269 | -1.221617265 | 4.64E-04 |
| Q8TEH3 | T923 | -1.203429085 | 3.19E-05 |
| Q9BYN0 | T8 | -1.167116736 | 4.96E-05 |
| Q15058 | S272 | -1.165446493 | 9.74E-06 |
| Q86SQ0 | T898 | -1.160435859 | 3.96E-07 |
| Q6IQ23 | T472 | -1.133086137 | 1.82E-04 |
| Q9NV70 | T445 | -1.123674943 | 2.54E-08 |
| Q5T5P2 | T1760 | -1.120340964 | 1.85E-05 |
| Q04721 | S2093 | -1.09596714 | 1.22E-06 |
| O75146 | S1045 | -1.094187269 | 1.09E-05 |
| Q15059 | S542 | -1.091349078 | 1.98E-05 |
| Q8IVL1 | S1215 | -1.0731331 | 2.03E-05 |
| Q6R327 | T1332 | -1.041692126 | 1.05E-05 |
| O75122 | S360 | -1.033839303 | 9.18E-06 |
| Q99959 | T118 | -1.026275863 | 3.22E-06 |
| Q7Z2Z1 | S1045 | -0.991101049 | 0.009564698 |
| Q16181 | T426 | -0.987232757 | 3.65E-06 |
| Q9UQC2 | T385 | -0.98559854 | 1.40E-05 |
| Q6ZT07 | S267 | -0.983842651 | 1.18E-04 |
| Q8IVT5 | T425 | -0.983087558 | 2.19E-07 |
| Q9NXV6 | S151 | -0.971312438 | 3.96E-07 |
| Q9HAU0 | T438 | -0.966190493 | 2.76E-07 |
| P23508 | S681 | -0.963527723 | 6.81E-06 |
| Q9BTT6 | T474 | -0.962871739 | 2.06E-04 |
| Q92556 | S344 | -0.958543954 | 5.66E-08 |
| Q9Y448 | T153 | -0.958136504 | 1.49E-05 |
| Q7Z460 | S1091 | -0.956730721 | 7.10E-06 |
| Q07157 | T589 | -0.948760819 | 3.65E-06 |
| Q15811 | S559 | -0.946586015 | 0.001867503 |
| Q96QZ7 | S1361_T1365 | -0.93753858 | 0.001590938 |

| | | | |
|---|---|---|---|
| Q13435 | S883 | -0.935934566 | 6.70E-06 |
| Q9HAU0 | S382 | -0.93276788 | 1.82E-04 |
| Q5T5P2 | S1084 | -0.925159608 | 4.06E-05 |
| Q99501 | T376 | -0.924542471 | 2.52E-04 |
| Q9BQ39 | HHYDSDEKSETR | -0.91727353 | 0.024921716 |
| P84098 | S189 | -0.907679996 | 0.033723097 |
| Q96MU7 | T148 | -0.878887347 | 0.037829431 |
| Q9UBP0 | S268 | -0.87864525 | 3.19E-05 |
| P38432 | S271 | -0.877882717 | 2.36E-05 |
| Q12923 | T320 | -0.875361226 | 6.47E-05 |
| Q9UHV9 | S14 | -0.867535695 | 0.01846624 |
| Q9C0D5 | S214 | -0.864768775 | 0.003689508 |
| Q6NSI4 | S39 | -0.864481975 | 8.28E-04 |
| Q13177 | T134 | -0.861904323 | 1.98E-05 |
| Q6ZVF9 | S526 | -0.858466865 | 4.26E-04 |
| Q9Y2I1 | S467 | -0.853827107 | 1.98E-05 |
| O60716 | T59 | -0.851516547 | 2.50E-06 |
| Q9H0B6 | S539 | -0.850899358 | 6.01E-05 |
| Q9UNL2 | S105 | -0.849638511 | 7.10E-06 |
| Q9UQB8 | T340 | -0.841768662 | 1.98E-05 |
| Q96F86 | S161 | -0.835192327 | 0.027103985 |
| Q86VW2 | S490 | -0.834038972 | 9.40E-05 |
| Q6UB98 | S1143 | -0.833942879 | 1.09E-05 |
| Q13136 | T1159 | -0.829502195 | 0.003240581 |
| Q8TEW0 | S1312 | -0.827854685 | 0.005932397 |
| Q9P2M7 | T34 | -0.820859584 | 7.22E-06 |
| Q7Z4S6 | S853 | -0.807412946 | 2.71E-05 |
| O95490 | YSSGTQSR | -0.806154668 | 2.29E-04 |
| P35658 | S1023 | -0.803412654 | 0.036413724 |
| P53801 | Y174 | -0.802538375 | 0.026440093 |
| Q15181 | S250 | -0.791148671 | 0.012922115 |
| Q9Y2J4 | S661 | -0.788388981 | 6.70E-06 |
| P35611 | S135 | -0.784012053 | 5.13E-05 |
| Q9Y5A9 | S394 | -0.783342619 | 8.07E-05 |
| Q07157 | T1167 | -0.782506262 | 0.001519252 |
| Q04637 | S704 | -0.779217567 | 4.92E-05 |
| Q8NEY1 | T942 | -0.775370459 | 8.50E-06 |
| Q8TEW0 | S1223 | -0.773775296 | 0.007082264 |
| P35568 | S794 | -0.773038275 | 1.40E-05 |
| O14646 | S1677_S | -0.770864253 | 0.001041057 |
| Q5SYE7 | S662 | -0.765006156 | 6.44E-05 |
| P35241 | S533 | -0.758105065 | 0.032165945 |
| Q8NHQ8 | S387 | -0.753901899 | 1.58E-05 |
| P49006 | S119_S/T | -0.750664741 | 0.008382431 |
| Q76L83 | SSLTQEEAPVSW | -0.747317101 | 1.76E-04 |
| O43491 | SQPPPAAESQSSL | -0.744424857 | 0.022572892 |
| O00192 | S203 | -0.743203841 | 7.96E-05 |
| Q96T37 | S58 | -0.741300775 | 4.06E-05 |
| Q9C0D5 | T687 | -0.741016038 | 3.56E-05 |
| Q92609 | S522 | -0.732813312 | 4.87E-04 |
| Q8NFC6 | S1017 | -0.731311408 | 3.50E-05 |
| P35711 | S424 | -0.72351618 | 3.65E-04 |
| P20290 | T82 | -0.722701934 | 3.56E-04 |
| O60749 | S226 | -0.713993177 | 9.74E-06 |
| P53367 | S132 | -0.713359032 | 0.001745391 |
| Q9HAU0 | T460 | -0.709642163 | 1.58E-05 |

| | | | |
|---|---|---|---|
| O43491 | SSDPEEEKGSQP | -0.70374801 | 3.83E-05 |
| Q6ZVF9 | S460 | -0.697277545 | 3.22E-06 |
| Q9Y4D1 | T675 | -0.692897914 | 2.10E-04 |
| Q8TEW0 | S201 | -0.675576314 | 0.001127271 |
| P25098 | S685 | -0.674065557 | 0.015128319 |
| Q92551 | S197 | -0.673421144 | 3.19E-04 |
| Q4VC05 | S207 | -0.666631503 | 1.07E-05 |
| Q9Y4K1 | S410 | -0.66589918 | 5.95E-04 |
| P05412 | S63 | -0.661607607 | 0.002195488 |
| Q04637 | S1209 | -0.658995596 | 1.98E-05 |
| Q12767 | S668 | -0.658440952 | 3.13E-04 |
| O60716 | S230 | -0.658377386 | 1.58E-05 |
| Q9UHY8 | S65 | -0.65650915 | 4.96E-05 |
| Q93052 | S609 | -0.651039959 | 0.001759769 |
| Q96F81 | S1350 | -0.646940701 | 6.59E-05 |
| Q9Y2J4 | S166 | -0.642205639 | 8.55E-05 |
| Q6DN90 | T901 | -0.634836063 | 0.006958044 |
| O96013 | T207 | -0.634461386 | 2.03E-05 |
| Q5VZ46 | S518 | -0.62945503 | 0.045452161 |
| Q01082 | T2187 | -0.628269648 | 0.008566303 |
| P49815 | GPLPSSSPRSPS | -0.625526601 | 0.005117384 |
| Q6R327 | T1103 | -0.623297144 | 5.30E-04 |
| P06730 | S209 | -0.623207968 | 2.01E-04 |
| Q9UQB8 | T360 | -0.622140051 | 6.14E-04 |
| Q8WUH6 | S77 | -0.621976805 | 0.003452289 |
| Q9UIW2 | T1615 | -0.619758382 | 8.28E-04 |
| Q96CW1 | S236 | -0.618851269 | 5.86E-05 |
| Q96NL6 | S656 | -0.61873091 | 0.015050147 |
| Q04637 | S1097 | -0.618253492 | 1.41E-04 |
| P55196 | S1275 | -0.616017966 | 1.07E-05 |
| P13073 | S56 | -0.613448061 | 0.001514523 |
| O95819 | T834 | -0.610065352 | 0.018095709 |
| Q9Y5S2 | S524 | -0.604367641 | 0.001354977 |
| P60468 | S49 | -0.603939988 | 2.71E-05 |
| Q9BRK4 | S570 | -0.602871106 | 3.37E-05 |
| Q8IZP0 | T196 | 0.600332281 | 0.004231049 |
| P62753 | S235_S236 | 0.607886321 | 0.015647504 |
| Q16513 | S21 | 0.611142248 | 6.67E-04 |
| Q96T88 | ASATSSSQRDW | 0.621837233 | 0.004516747 |
| Q9UGH3 | S639_S/T | 0.63015119 | 0.021939966 |
| Q03111 | S209 | 0.632054136 | 0.00974682 |
| Q04637 | SSLSRER | 0.639810501 | 0.001966976 |
| Q14241 | S322 | 0.640312185 | 0.009564698 |
| Q9Y6G9 | T408 | 0.643604984 | 0.002710202 |
| P62753 | S236_S240 | 0.644408149 | 0.017151429 |
| Q5VSY0 | S360 | 0.653167884 | 0.006739153 |
| O15040 | S408 | 0.655953223 | 0.008208954 |
| Q9H4M7 | T296 | 0.667004206 | 0.029726219 |
| P27448 | SVSSSQKQR | 0.673577433 | 0.001071305 |
| Q13233 | S252 | 0.695203491 | 3.53E-06 |
| P49023 | Y118 | 0.698254049 | 0.007966093 |
| P17096 | S9 | 0.708053041 | 0.006462963 |
| Q13233 | S250_S252 | 0.711410445 | 3.94E-04 |
| P62753 | S244_S247 | 0.717066182 | 0.001757042 |
| Q07157 | S1124 | 0.720974107 | 4.76E-04 |
| O75694 | S992 | 0.758641103 | 2.52E-04 |

| | | | |
|---|---|---|---|
| Q9Y4I1 | S600 | 0.77882986 | 7.64E-04 |
| Q05D32 | S9 | 0.809866775 | 1.98E-05 |
| Q9UPW6 | S39 | 0.82928638 | 5.08E-04 |
| Q5QJE6 | S245 | 0.831169048 | 0.001645587 |
| Q9Y2I9 | S787 | 0.836173338 | 0.019605992 |
| Q8TBZ3 | S347 | 0.851679974 | 0.013861972 |
| O95819 | S710_S715 | 0.865250429 | 5.62E-06 |
| O60238 | LEHVPSSSSIHNG | 0.91091649 | 0.017864016 |
| O95819 | S550 | 0.956989147 | 1.98E-05 |
| O15550 | S829 | 0.966406805 | 2.10E-04 |
| Q99502 | S314 | 0.999118334 | 0.036974231 |
| P15927 | S174 | 0.999549223 | 7.64E-04 |
| Q9H9D4 | S346 | 1.052553265 | 4.68E-04 |
| Q8NC56 | S138_S/T | 1.0911335 | 0.013560244 |
| Q92917 | T138 | 1.14968258 | 0.027583976 |
| Q5SW79 | S807 | 1.150341015 | 3.94E-04 |
| Q96B36 | S203 | 1.168131014 | 0.007942145 |
| Q07157 | S144 | 1.248523138 | 0.045642212 |
| Q9NS37 | S50 | 1.28098435 | 2.01E-04 |
| O14668 | S158 | 1.321636135 | 1.40E-05 |
| Q9Y4K4 | T168 | 1.411432511 | 2.26E-08 |
| P62491 | S182 | 1.41861915 | 5.52E-04 |
| Q9Y520 | S867 | 1.77222641 | 1.98E-05 |
| Q9NYZ3 | S303 | 1.838108759 | 5.70E-05 |
| P17544 | T326 | 2.311651391 | 7.84E-06 |
| Q9H000 | S365 | 2.366027162 | 1.09E-05 |
| Q8TCN5 | S195 | 3.429181246 | 2.03E-05 |
