## Supplementary table 4 for "MAP4K inhibition by the CNS penetrant inhibitor famlasertib restrains medulloblastoma dissemination without developmental toxicity"

| Master_id_first | PsiteInProtein | diff | FDR |
|---|---|---|---|
| Q14667 | T1269 | -3.897270029 | 3.64E-05 |
| Q8TF76 | S147 | -3.111764272 | 4.15E-05 |
| Q9ULF5 | T540 | -2.253257277 | 6.40E-09 |
| Q6GQQ9 | T547 | -2.109511673 | 6.52E-10 |
| Q5VWQ0 | S92 | -1.995271816 | 2.46E-04 |
| P55196 | T566 | -1.968164062 | 4.51E-06 |
| Q96B97 | T354 | -1.928075585 | 2.65E-06 |
| P55010 | PPENSDSGTGKK | -1.890567278 | 5.18E-09 |
| Q7Z2Z1 | S1045 | -1.879787361 | 2.68E-05 |
| Q96PY5 | T467 | -1.592323263 | 1.71E-08 |
| Q9Y490 | T144 | -1.589549959 | 1.25E-07 |
| Q9UBT7 | S538 | -1.569057463 | 6.72E-09 |
| P55010 | T153 | -1.513847557 | 6.40E-09 |
| Q9HAU0 | S855_T868 | -1.47647664 | 6.40E-09 |
| Q14008 | T569 | -1.471143207 | 3.65E-07 |
| O60716 | T177 | -1.440794187 | 2.26E-07 |
| Q9HBI1 | T28 | -1.427617944 | 7.30E-06 |
| Q96PY5 | T450 | -1.371990208 | 6.38E-06 |
| O60716 | T916 | -1.323012224 | 2.95E-06 |
| Q96QZ7 | S1361_T1365 | -1.313458116 | 6.04E-05 |
| Q8NHQ8 | PFQSGSLKRPGS | -1.310406961 | 2.00E-08 |
| Q5T5P2 | T1760 | -1.306459221 | 4.51E-06 |
| Q8TEA8 | S197 | -1.283689603 | 0.001067563 |
| Q15058 | S272 | -1.279639101 | 4.39E-06 |
| O15355 | S527 | -1.264191852 | 1.08E-08 |
| Q9HAU0 | T868 | -1.193405765 | 2.85E-08 |
| Q5HYJ3 | S227 | -1.192708922 | 4.51E-06 |
| Q86SQ0 | T898 | -1.187717161 | 3.16E-07 |
| O75146 | S1045 | -1.121503779 | 8.85E-06 |
| Q8TEH3 | T923 | -1.11237875 | 6.90E-05 |
| Q7Z460 | S1091 | -1.110170243 | 1.76E-06 |
| Q9ULR3 | S221 | -1.079141106 | 0.006498753 |
| P23508 | S681 | -1.074085351 | 2.42E-06 |
| O75122 | S360 | -1.068998189 | 6.65E-06 |
| Q8IVL1 | S1215 | -1.068792407 | 2.28E-05 |
| Q04721 | S2093 | -1.063341282 | 1.84E-06 |
| Q99501 | T376 | -1.056038761 | 6.47E-05 |
| Q9UQC2 | T385 | -1.052492214 | 7.30E-06 |
| Q9NXV6 | S151 | -1.050736564 | 1.70E-07 |
| Q6R327 | T1332 | -1.037484133 | 1.02E-05 |
| Q9BYN0 | T8 | -1.025650001 | 1.82E-04 |
| Q8TEW0 | S1223 | -1.023219543 | 6.20E-04 |
| Q9NV70 | T445 | -1.022142923 | 9.74E-08 |
| Q6IQ23 | T472 | -1.021085233 | 4.72E-04 |
| Q5T5P2 | S1084 | -1.00510416 | 1.82E-05 |
| Q9H0B6 | S539 | -0.987443983 | 1.37E-05 |
| P35711 | S424 | -0.979899755 | 1.77E-05 |
| P84098 | S189 | -0.976883342 | 0.023719 |
| Q15811 | S559 | -0.970358982 | 0.001537497 |
| Q8IVT5 | T425 | -0.961983734 | 2.97E-07 |
| Q9HAU0 | T438 | -0.959172131 | 3.05E-07 |
| Q16181 | T426 | -0.955707187 | 5.13E-06 |
| Q9Y448 | T153 | -0.947167731 | 1.59E-05 |
| O95490 | YSSGTQSR | -0.946952527 | 4.30E-05 |

| | | | |
|---|---|---|---|
| Q15059 | S542 | -0.946496005 | 8.15E-05 |
| Q9UHV9 | S14 | -0.942353834 | 0.010655382 |
| Q99959 | T118 | -0.939904591 | 7.00E-06 |
| Q9P2M7 | T34 | -0.936722866 | 2.11E-06 |
| Q9C0D5 | S214 | -0.933637823 | 0.001972168 |
| Q9Y4B5 | S1514 | -0.927535416 | 0.006076837 |
| Q13177 | T134 | -0.92139234 | 1.04E-05 |
| Q07157 | T589 | -0.902328878 | 5.87E-06 |
| Q92556 | S344 | -0.898392825 | 1.41E-07 |
| Q6ZT07 | S267 | -0.894707058 | 2.91E-04 |
| Q9UQB8 | T340 | -0.892680192 | 1.04E-05 |
| Q9BSI4 | S330 | -0.890230362 | 0.034674629 |
| Q93052 | S609 | -0.876407012 | 1.00E-04 |
| Q6ZVF9 | S526 | -0.872847549 | 3.52E-04 |
| P18615 | S37 | -0.870512116 | 0.028637832 |
| Q6WCQ1 | S269 | -0.867872713 | 0.008995812 |
| Q9HAU0 | S382 | -0.864535207 | 3.52E-04 |
| A4D1S0 | S195 | -0.861523201 | 7.56E-04 |
| Q96N67 | T142 | -0.85877586 | 0.030832558 |
| Q9H4Z2 | S1127 | -0.856998314 | 0.03835815 |
| P35568 | S794 | -0.856465658 | 5.70E-06 |
| Q6DN90 | T901 | -0.855058703 | 5.05E-04 |
| Q76L83 | SSLTQEEAPVSW | -0.852763373 | 4.21E-05 |
| P23508 | S199 | -0.848640753 | 9.01E-06 |
| Q6UB98 | S1143 | -0.837025928 | 1.02E-05 |
| Q86VW2 | S490 | -0.835969095 | 9.01E-05 |
| Q13435 | S883 | -0.8343459 | 1.71E-05 |
| P20290 | T82 | -0.83358665 | 8.45E-05 |
| Q8NEY1 | T942 | -0.830926515 | 4.51E-06 |
| Q9UNL2 | S105 | -0.829593223 | 8.57E-06 |
| Q9P219 | S1778 | -0.82545321 | 4.06E-05 |
| Q9NRX4 | S29 | -0.817038922 | 0.022520551 |
| P38432 | S271 | -0.813559077 | 5.11E-05 |
| O60716 | S230 | -0.802950619 | 2.58E-06 |
| Q9Y2I1 | S467 | -0.800224515 | 3.64E-05 |
| Q8TEW0 | S201 | -0.792081101 | 2.46E-04 |
| Q9UQB8 | T360 | -0.786145467 | 6.13E-05 |
| Q04637 | S704 | -0.783825075 | 4.35E-05 |
| Q96F81 | S1350 | -0.776997278 | 1.06E-05 |
| Q8IY63 | S269 | -0.77478621 | 2.65E-05 |
| Q9Y5A9 | S394 | -0.773942719 | 9.01E-05 |
| Q6NSI4 | S39 | -0.771838078 | 0.002282837 |
| Q8TEW0 | S1312 | -0.761767497 | 0.011294537 |
| Q9NRP7 | T384 | -0.760455562 | 0.013264442 |
| Q9Y2J4 | S661 | -0.75789742 | 8.86E-06 |
| Q9C0D5 | T687 | -0.752860288 | 3.06E-05 |
| O60716 | T59 | -0.750640791 | 7.33E-06 |
| Q9Y4K1 | S410 | -0.748125423 | 1.96E-04 |
| Q8NFC6 | S1017 | -0.73202413 | 3.41E-05 |
| P49815 | GPLPSSSPRSPS | -0.720835337 | 0.001572531 |
| Q9BTT6 | T474 | -0.71998672 | 0.002848963 |
| Q5VZ46 | S518 | -0.719936059 | 0.022520551 |
| O14974 | S422 | -0.715750657 | 4.72E-04 |
| P41236 | S20 | -0.714688072 | 1.02E-05 |
| Q9Y4D1 | T675 | -0.70906047 | 1.60E-04 |
| Q9Y2J4 | S166 | -0.705924438 | 3.24E-05 |

| | | | |
|---|---|---|---|
| P18615 | QSSSSTTSQGGV | -0.70273981 | 2.69E-04 |
| Q6ZSZ5 | S1202 | -0.702104325 | 0.001594534 |
| P53367 | S132 | -0.701585265 | 0.001989616 |
| P60468 | S49 | -0.697456691 | 7.30E-06 |
| O43491 | SQPPPAAESQSSL | -0.688381156 | 0.041474631 |
| Q96T37 | S58 | -0.68619611 | 8.82E-05 |
| O96013 | S148 | -0.683971169 | 3.45E-04 |
| Q9UPP1 | S804 | -0.681036532 | 0.009272342 |
| Q8WUH6 | S77 | -0.679184437 | 0.001637079 |
| O43776 | S88 | -0.677849431 | 0.001299208 |
| O96013 | T207 | -0.676564287 | 1.08E-05 |
| O43491 | SSDPEEEKGSQPI | -0.676340086 | 5.70E-05 |
| P55196 | T1330_T1334 | -0.670656679 | 0.002604849 |
| Q12923 | T320 | -0.668314074 | 9.48E-04 |
| P35611 | S135 | -0.6682231 | 2.57E-04 |
| Q9UHY8 | S65 | -0.666392622 | 4.15E-05 |
| Q9H6R4 | S283 | -0.664763593 | 0.001972528 |
| Q9UF83 | S186 | -0.661393355 | 2.32E-04 |
| P15260 | T295 | -0.659943068 | 0.001696665 |
| Q4VC05 | S207 | -0.658444155 | 1.08E-05 |
| Q8WXE0 | S358 | -0.656217622 | 0.00149773 |
| O14639 | S655 | -0.65159831 | 3.10E-04 |
| Q6ZVF9 | S460 | -0.650198949 | 5.87E-06 |
| Q8NHQ8 | S387 | -0.648917915 | 6.51E-05 |
| P53985 | S213 | -0.64690587 | 2.39E-05 |
| Q5SYE7 | S662 | -0.64513933 | 3.45E-04 |
| Q9H2J7 | S699_S701 | -0.644416747 | 1.80E-04 |
| Q14978 | S456 | -0.636762257 | 1.17E-04 |
| Q9NR09 | GSSSLDR | -0.636063853 | 2.56E-05 |
| Q04637 | S1097 | -0.635633379 | 9.92E-05 |
| O75445 | T5111 | -0.634266373 | 0.030462239 |
| Q5T5U3 | S623 | -0.632042907 | 1.28E-04 |
| Q92551 | S197 | -0.63020549 | 5.94E-04 |
| O60749 | S226 | -0.629576875 | 3.06E-05 |
| Q04637 | S1209 | -0.628358396 | 3.22E-05 |
| Q07157 | T1167 | -0.628243663 | 0.009268279 |
| Q6R327 | T1103 | -0.623624282 | 5.41E-04 |
| Q92609 | S522 | -0.619334482 | 0.002277158 |
| Q07157 | S1142 | -0.619043859 | 3.17E-04 |
| Q9UIW2 | T1615 | -0.617372501 | 8.93E-04 |
| P05412 | S63 | -0.613102856 | 0.004066491 |
| P78314 | S444 | -0.607607525 | 0.009557789 |
| Q13620 | S139_S/T | -0.604639262 | 0.002321885 |
| O75122 | S430 | -0.602617795 | 0.002346961 |
| Q96T88 | ASATSSSQRDW( | 0.603223813 | 0.005811869 |
| Q9BZ29 | S32 | 0.604027304 | 8.36E-04 |
| O15040 | S408 | 0.610483671 | 0.014512663 |
| Q03111 | S209 | 0.617393685 | 0.011936949 |
| O15018 | S1919 | 0.63992366 | 0.004237228 |
| Q5VSY0 | S360 | 0.651370468 | 0.006678167 |
| Q04637 | SSLSRER | 0.657206927 | 0.001575627 |
| Q9NY43 | GDREITSSRESPI | 0.658492797 | 0.015701416 |
| Q5QJE6 | S245 | 0.666274923 | 0.010150864 |
| P62753 | S244_S247 | 0.675984398 | 0.002826612 |
| P62753 | S236_S240 | 0.682638907 | 0.011913895 |
| P62753 | S235_S236 | 0.715086609 | 0.004434669 |

| | | | |
|---|---|---|---|
| P62753 | S236_T241_S242 | 0.786339302 | 0.00115455 |
| O95819 | S710_S715 | 0.800385279 | 1.02E-05 |
| Q9H9D4 | S346 | 0.804124558 | 0.004952525 |
| O95819 | S550 | 0.908459599 | 3.22E-05 |
| O15550 | S829 | 0.92871089 | 3.05E-04 |
| Q9Y4I1 | S600 | 0.938199861 | 1.25E-04 |
| Q8TAQ2 | S283_S286 | 0.947492692 | 0.042224676 |
| O14668 | S158 | 0.978060558 | 2.60E-04 |
| Q9NS37 | S50 | 0.988572724 | 0.002167879 |
| P17544 | T326 | 1.066893702 | 0.009552836 |
| Q02156 | S368 | 1.231544168 | 0.009362713 |
| Q9Y4K4 | T168 | 1.244510669 | 9.86E-08 |
| Q9Y520 | S867 | 1.280731448 | 5.02E-04 |
| Q9H1B7 | S334_S337 | 1.345975911 | 0.00245989 |
| Q96AP0 | S339_S349 | 1.53600724 | 0.008911557 |
| Q99502 | S314 | 1.552656189 | 0.001442827 |
| Q9H000 | S365 | 1.622970621 | 4.28E-04 |
| Q9NYZ3 | S303 | 1.658266895 | 1.58E-04 |
| Q8NC56 | S138_S/T | 1.727187343 | 2.60E-04 |
| Q07157 | S144 | 2.008249123 | 0.001446207 |
| P62491 | S182 | 2.129519249 | 1.02E-05 |
| Q9H4A3 | S2029_S2032 | 2.31598185 | 0.016742946 |
| Q8TCN5 | S195 | 3.477290925 | 1.88E-05 |
