## Supplementary table 5 for "MAP4K inhibition by the CNS penetrant inhibitor famlasertib restrains medulloblastoma dissemination without developmental toxicity"

| Kinase.Gene | mS | Enrichment | m | z.score | p.value | FDR |
|---|---|---|---|---|---|---|
| MAPK15 | -0.6616076 | -178.69973 | 1 | -3.6875303 | 0.00011322 | 0.00784052 |
| PLK3 | -0.3407844 | -92.045613 | 3 | -3.30708 | 0.00047137 | 0.02611389 |
| PRKD1 | -0.1446301 | -39.064477 | 14 | -3.0761794 | 0.00104836 | 0.04156525 |
| PBK | -0.3624304 | -97.892179 | 2 | -2.8698891 | 0.00205308 | 0.05687028 |
| MAPKAPK3 | -0.3434853 | -92.775116 | 2 | -2.72139 | 0.0032504 | 0.06925855 |
| MOK | -0.2366954 | -63.931248 | 4 | -2.6648448 | 0.00385119 | 0.07619861 |
| PRKD2 | -0.2547029 | -68.795064 | 3 | -2.4806958 | 0.00655631 | 0.09558412 |
| MAPKAPK2 | -0.1028408 | -27.77722 | 15 | -2.2870868 | 0.01109538 | 0.11820847 |
| PIM2 | -0.1596115 | -43.110947 | 6 | -2.2172271 | 0.01330379 | 0.1290034 |
| MKNK1 | -0.2784159 | -75.199918 | 2 | -2.211351 | 0.01350577 | 0.1290034 |
| MKNK2 | -0.2784159 | -75.199918 | 2 | -2.211351 | 0.01350577 | 0.1290034 |
| JAK2 | -0.2659476 | -71.832248 | 2 | -2.1136199 | 0.01727387 | 0.1500623 |
| CAMK2B | -0.0665905 | -17.986043 | 26 | -1.9865949 | 0.02348366 | 0.18157111 |
| CAMK2A | -0.0606061 | -16.369664 | 31 | -1.9845426 | 0.02359769 | 0.18157111 |
| DMPK | -0.0943838 | -25.492993 | 13 | -1.9601571 | 0.02498872 | 0.18707768 |
| STK11 | -0.1116528 | -30.157328 | 9 | -1.9180929 | 0.02754962 | 0.19567293 |
| PRKCB | -0.0168148 | -4.5416641 | 243 | -1.7726847 | 0.03814048 | 0.25154555 |
| PLK2 | -0.1256426 | -33.935974 | 6 | -1.7560491 | 0.03954001 | 0.25471125 |
| PRKCQ | -0.1491336 | -40.280881 | 4 | -1.6942096 | 0.04511274 | 0.27244938 |
| VRK1 | -0.1726333 | -46.628127 | 3 | -1.6928259 | 0.0452443 | 0.27244938 |
| BRAF | -0.1446349 | -39.065794 | 4 | -1.6443411 | 0.05005288 | 0.28884683 |
| RET | -0.1649674 | -44.557579 | 3 | -1.6192333 | 0.05269853 | 0.29703209 |
| CSNK2A2 | -0.0335072 | -9.0502772 | 61 | -1.6107611 | 0.0536159 | 0.29703209 |
| MAP2K2 | -0.0496253 | -13.403762 | 29 | -1.5917069 | 0.05572529 | 0.3026648 |
| NEK1 | -0.0501203 | -13.537459 | 27 | -1.5500959 | 0.06055926 | 0.30784897 |
| PRKACG | -0.0509793 | -13.769471 | 26 | -1.5453961 | 0.06112525 | 0.30784897 |
| CSNK2A1 | -0.0120791 | -3.2625658 | 269 | -1.4346151 | 0.0756984 | 0.36152512 |
| EEF2K | -0.1764055 | -47.647005 | 2 | -1.4117548 | 0.07901108 | 0.37095034 |
| PIM3 | -0.1071779 | -28.948662 | 5 | -1.3742021 | 0.08468948 | 0.38512029 |
| CHEK1 | -0.0359767 | -9.7172898 | 39 | -1.3734266 | 0.08480988 | 0.38512029 |
| GSK3B | -0.0150598 | -4.0676498 | 172 | -1.3638261 | 0.08631116 | 0.38561597 |
| PRKCD | -0.0307052 | -8.2934509 | 50 | -1.3484976 | 0.08874919 | 0.3902147 |
| ATM | -0.0260478 | -7.0355029 | 63 | -1.3087947 | 0.09530196 | 0.4051012 |
| ILK | -0.2309724 | -62.385482 | 1 | -1.3007025 | 0.09668016 | 0.4051012 |
| CDK19 | -0.2298413 | -62.079969 | 1 | -1.2944332 | 0.09775792 | 0.4051012 |
| MAPK7 | -0.0267114 | -7.2147367 | 54 | -1.2387358 | 0.10772167 | 0.42627005 |
| PRKACB | -0.0416016 | -11.236551 | 24 | -1.2301352 | 0.10932325 | 0.42651464 |
| CDKL5 | -0.1869539 | -50.496117 | 1 | -1.0567266 | 0.14531818 | 0.49719006 |
| MAPK14 | -0.0227493 | -6.144571 | 46 | -0.9943591 | 0.16002404 | 0.49719006 |
| MAP4K6 | -0.0831741 | -22.46526 | 4 | -0.9630383 | 0.16776416 | 0.49719006 |
| CAMK4 | -0.0370762 | -10.01425 | 18 | -0.9589133 | 0.16880121 | 0.49719006 |
| GRK5 | -0.0659846 | -17.822401 | 6 | -0.9461039 | 0.17204781 | 0.49719006 |
| PRKCI | -0.0227764 | -6.1518888 | 40 | -0.9281944 | 0.17665338 | 0.49719006 |
| WNK1 | -0.1138764 | -30.757921 | 2 | -0.9216271 | 0.17836156 | 0.49719006 |
| ADRBK2 | -0.0656288 | -17.726304 | 5 | -0.8592613 | 0.19509819 | 0.49719006 |
| MAPK1 | -0.0056091 | -1.5150106 | 272 | -0.8511624 | 0.19733956 | 0.49719006 |
| MARK2 | -0.0584393 | -15.784419 | 6 | -0.8436651 | 0.19942828 | 0.49719006 |
| PAK5 | -0.0540526 | -14.599558 | 6 | -0.7841084 | 0.21648827 | 0.49719006 |
| ROCK2 | -0.0201656 | -5.4467098 | 34 | -0.7713753 | 0.22024224 | 0.49719006 |
| PAK1 | -0.015383 | -4.1549266 | 50 | -0.7479898 | 0.22723316 | 0.49719006 |
| DAPK2 | -0.1300197 | -35.118235 | 1 | -0.7411646 | 0.22929681 | 0.49719006 |
| CDC42BPA | -0.0315213 | -8.5138755 | 14 | -0.7304822 | 0.23254774 | 0.49719006 |
| CDC7 | -0.0618075 | -16.694159 | 4 | -0.7261864 | 0.23386226 | 0.49719006 |
| PRKG1 | -0.0139892 | -3.7784699 | 52 | -0.7070965 | 0.23975325 | 0.49719006 |

| | | | | | | |
|---|---|---|---|---|---|---|
| STK10 | -0.0851352 | -22.994947 | 2 | -0.6963426 | 0.24310715 | 0.49719006 |
| TLK2 | -0.0469165 | -12.672123 | 6 | -0.6872263 | 0.24597007 | 0.49719006 |
| DYRK1A | -0.0492553 | -13.303833 | 5 | -0.6563351 | 0.25580428 | 0.49719006 |
| MAP2K1 | -0.0100217 | -2.7068548 | 69 | -0.6318563 | 0.26374041 | 0.49719006 |
| GRK7 | -0.0445977 | -12.045794 | 5 | -0.5986096 | 0.27471662 | 0.49719006 |
| LCK | -0.0434963 | -11.748326 | 5 | -0.5849602 | 0.27928723 | 0.49719006 |
| ADRBK1 | -0.010145 | -2.7401689 | 51 | -0.5481059 | 0.2918096 | 0.49719006 |
| PRKCZ | -0.0136721 | -3.6928376 | 32 | -0.5447518 | 0.29296211 | 0.49719006 |
| MAPK9 | -0.0114346 | -3.0884856 | 42 | -0.54372 | 0.29331709 | 0.49719006 |
| ACVR1B | -0.0938157 | -25.339557 | 1 | -0.5405012 | 0.29442573 | 0.49719006 |
| RPS6KA3 | -0.0146941 | -3.9688693 | 28 | -0.5395414 | 0.29475668 | 0.49719006 |
| PDHK3 | -0.0629366 | -16.999129 | 2 | -0.5223417 | 0.30071623 | 0.49719006 |
| PRKCG | -0.0076271 | -2.060087 | 68 | -0.5178171 | 0.30229294 | 0.49719006 |
| TLK1 | -0.0333542 | -9.0089513 | 6 | -0.5030978 | 0.30744777 | 0.49719006 |
| TGFBR2 | -0.0154226 | -4.1656219 | 20 | -0.4740518 | 0.31773149 | 0.49719006 |
| ARAF | -0.0113174 | -3.0568109 | 31 | -0.4635043 | 0.32150146 | 0.49719006 |
| MAPKAPK5 | -0.0282231 | -7.6230336 | 6 | -0.4334349 | 0.33234943 | 0.49719006 |
| LIMK1 | -0.0727275 | -19.643651 | 1 | -0.4236183 | 0.33592211 | 0.49719006 |
| LIMK2 | -0.0727275 | -19.643651 | 1 | -0.4236183 | 0.33592211 | 0.49719006 |
| TESK1 | -0.0727275 | -19.643651 | 1 | -0.4236183 | 0.33592211 | 0.49719006 |
| CLK4 | -0.0502672 | -13.577135 | 2 | -0.4230341 | 0.3361352 | 0.49719006 |
| ACVR2B | -0.0400317 | -10.812542 | 3 | -0.4198481 | 0.33729822 | 0.49719006 |
| CAMK2G | -0.0149769 | -4.0452406 | 16 | -0.4141237 | 0.33939177 | 0.49719006 |
| CLK3 | -0.0368166 | -9.9441248 | 3 | -0.3889823 | 0.34864462 | 0.49719006 |
| PAK6 | -0.0307542 | -8.3066993 | 4 | -0.3819565 | 0.35124683 | 0.49719006 |
| AAK1 | -0.0639264 | -17.266469 | 1 | -0.3748373 | 0.35389074 | 0.49719006 |
| IKBKB | -0.003371 | -0.9105006 | 86 | -0.3635671 | 0.35809065 | 0.49719006 |
| PRKCT | -0.0046168 | -1.2469994 | 49 | -0.322767 | 0.37343584 | 0.49719006 |
| CSNK2B | -0.035424 | -9.5679934 | 2 | -0.3066872 | 0.37954072 | 0.49719006 |
| MTOR | -0.0074791 | -2.0201051 | 24 | -0.3036098 | 0.38071257 | 0.49719006 |
| MAP3K8 | -0.0278207 | -7.5143478 | 3 | -0.3026218 | 0.38108906 | 0.49719006 |
| ATR | -0.0064625 | -1.7455258 | 27 | -0.2927491 | 0.38485697 | 0.49719006 |
| PKD1 | -0.0056451 | -1.5247476 | 31 | -0.2884609 | 0.38649697 | 0.49719006 |
| HIPK1 | -0.0298961 | -8.0749191 | 2 | -0.2633576 | 0.39613747 | 0.49719006 |
| RIPK1 | -0.0437574 | -11.81883 | 1 | -0.2630489 | 0.39625643 | 0.49719006 |
| MAP3K1 | -0.042233 | -11.407104 | 1 | -0.2546001 | 0.39951599 | 0.49719006 |
| WNK2 | -0.0399398 | -10.787711 | 1 | -0.2418899 | 0.40443276 | 0.49719006 |
| STK3 | -0.0383348 | -10.35419 | 1 | -0.2329938 | 0.40788312 | 0.49719006 |
| STK4 | -0.0383348 | -10.35419 | 1 | -0.2329938 | 0.40788312 | 0.49719006 |
| PRKG2 | -0.0033247 | -0.8979897 | 27 | -0.2023783 | 0.4198105 | 0.49719006 |
| PRKCE | -0.0007854 | -0.2121244 | 59 | -0.1910563 | 0.42424074 | 0.49719006 |
| MST1 | -0.0302231 | -8.1632361 | 1 | -0.1880343 | 0.42542489 | 0.49719006 |
| PDPK1 | -0.014831 | -4.0058555 | 3 | -0.177921 | 0.42939253 | 0.49719006 |
| WNK3 | -0.0273824 | -7.3959641 | 1 | -0.1722895 | 0.43160498 | 0.49719006 |
| DYRK2 | -0.0065795 | -1.7771122 | 8 | -0.1611857 | 0.43597356 | 0.49719006 |
| STK25 | -0.0239787 | -6.4766393 | 1 | -0.1534245 | 0.43903178 | 0.49719006 |
| CSNK1G2 | -0.0017403 | -0.4700472 | 19 | -0.1314911 | 0.44769343 | 0.49719006 |
| BCR | -0.0121103 | -3.2709818 | 2 | -0.1239455 | 0.4506792 | 0.49719006 |
| VRK2 | -0.0185222 | -5.0028459 | 1 | -0.1231815 | 0.4509817 | 0.49719006 |
| VRK3 | -0.0185222 | -5.0028459 | 1 | -0.1231815 | 0.4509817 | 0.49719006 |
| LMTK2 | -0.0183416 | -4.9540425 | 1 | -0.12218 | 0.45137823 | 0.49719006 |
| MAP2K5 | -0.0004259 | -0.1150241 | 18 | -0.0970753 | 0.4613333 | 0.49719006 |
| TRPM7 | -0.0029119 | -0.7865058 | 7 | -0.0969933 | 0.46136587 | 0.49719006 |
| CAMK1G | -0.0082563 | -2.2300288 | 2 | -0.0937367 | 0.46265917 | 0.49719006 |
| CAMK1 | -0.0006336 | -0.1711412 | 15 | -0.0930771 | 0.46292114 | 0.49719006 |
| CSNK1A1 | 0.00207123 | 0.55943796 | 76 | -0.0788138 | 0.46859038 | 0.49719006 |

| | | | | | | |
|---|---|---|---|---|---|---|
| ACTR2 | -0.0037968 | -1.0255045 | 3 | -0.0719916 | 0.47130428 | 0.49719006 |
| MAPK3 | 0.0030085 | 0.81259371 | 308 | -0.0674913 | 0.4730953 | 0.49719006 |
| BRSK1 | -0.0040581 | -1.0960797 | 2 | -0.060829 | 0.47574768 | 0.49719006 |
| BCKDK | 0.00011297 | 0.03051315 | 8 | -0.0562698 | 0.47756345 | 0.49719006 |
| PRKCH | 0.00221319 | 0.59778007 | 30 | -0.0452077 | 0.48197088 | 0.49719006 |
| MAP3K10 | -0.004381 | -1.1832922 | 1 | -0.0448023 | 0.48213246 | 0.49719006 |
| MAPK10 | 0.00244994 | 0.66172761 | 34 | -0.0404757 | 0.48385695 | 0.49719006 |
| TRPM6 | 0.00172137 | 0.46494195 | 12 | -0.0380347 | 0.48483002 | 0.49719006 |
| MAP3K5 | -0.0001318 | -0.0356037 | 3 | -0.036808 | 0.48531904 | 0.49719006 |
| MAP3K6 | -0.0001318 | -0.0356037 | 3 | -0.036808 | 0.48531904 | 0.49719006 |
| DYRK1B | -0.0028855 | -0.7793758 | 1 | -0.0365137 | 0.48543638 | 0.49719006 |
| CDK14 | 0.00013967 | 0.03772355 | 2 | -0.0279256 | 0.48886073 | 0.49719006 |
| CDK20 | 0.000431 | 0.11641279 | 1 | -0.0181317 | 0.49276691 | 0.49719006 |
| ICK | 0.0031412 | 0.84843713 | 11 | -0.0103152 | 0.49588491 | 0.49768159 |
| NEK11 | 0.00364056 | 0.98331138 | 4 | -0.0006849 | 0.49972676 | 0.49972676 |
| TAF1 | 0.00659676 | 1.78177898 | 1 | 0.0160425 | 0.49360024 | 0.49719006 |
| CAMK2D | 0.00432258 | 1.16752446 | 28 | 0.01819053 | 0.49274343 | 0.49719006 |
| PRKDC | 0.00412484 | 1.1141154 | 113 | 0.02489268 | 0.49007028 | 0.49719006 |
| GRK6 | 0.00841846 | 2.27382107 | 3 | 0.04527487 | 0.48194411 | 0.49719006 |
| GRK1 | 0.00707732 | 1.91157782 | 9 | 0.05611813 | 0.47762385 | 0.49719006 |
| PKD3 | 0.01043858 | 2.81945337 | 5 | 0.08348609 | 0.46673252 | 0.49719006 |
| CAMK1D | 0.01026628 | 2.77291522 | 6 | 0.0891152 | 0.46449518 | 0.49719006 |
| PAK7 | 0.01983819 | 5.35828143 | 1 | 0.08943416 | 0.46436844 | 0.49719006 |
| CSF1R | 0.01085475 | 2.93185914 | 6 | 0.09710448 | 0.46132171 | 0.49719006 |
| PIM1 | 0.00795951 | 2.14985793 | 18 | 0.10010796 | 0.46012931 | 0.49719006 |
| CLK2 | 0.00869002 | 2.34716875 | 17 | 0.11398157 | 0.4546262 | 0.49719006 |
| PLK4 | 0.00760712 | 2.05467881 | 30 | 0.11854112 | 0.45281946 | 0.49719006 |
| STK39 | 0.02103076 | 5.68039412 | 2 | 0.13582684 | 0.44597909 | 0.49719006 |
| MOS | 0.02959326 | 7.99311925 | 1 | 0.14350238 | 0.44294672 | 0.49719006 |
| IRAK1 | 0.03123417 | 8.43632758 | 1 | 0.15259724 | 0.43935795 | 0.49719006 |
| NEK6 | 0.03123417 | 8.43632758 | 1 | 0.15259724 | 0.43935795 | 0.49719006 |
| MAP2K3 | 0.00922135 | 2.49068033 | 25 | 0.15294761 | 0.43921979 | 0.49719006 |
| EPHA7 | 0.01760581 | 4.75531539 | 5 | 0.17231361 | 0.43159549 | 0.49719006 |
| TEC | 0.03791481 | 10.2407627 | 1 | 0.18962517 | 0.42480143 | 0.49719006 |
| MARK1 | 0.01091147 | 2.94717975 | 25 | 0.19978562 | 0.42082412 | 0.49719006 |
| NEK4 | 0.01713092 | 4.62704837 | 8 | 0.21051668 | 0.41663222 | 0.49719006 |
| MAP3K7 | 0.03087909 | 8.34041938 | 2 | 0.2130218 | 0.41565498 | 0.49719006 |
| BMX | 0.03196134 | 8.63273418 | 2 | 0.22150489 | 0.41234967 | 0.49719006 |
| PDK3 | 0.04368793 | 11.8000774 | 1 | 0.22162311 | 0.41230365 | 0.49719006 |
| PDK4 | 0.04368793 | 11.8000774 | 1 | 0.22162311 | 0.41230365 | 0.49719006 |
| PRKAA2 | 0.00832179 | 2.24771026 | 77 | 0.22467114 | 0.41111756 | 0.49719006 |
| TBK1 | 0.0338621 | 9.1461286 | 2 | 0.23640379 | 0.40655968 | 0.49719006 |
| PDHK4 | 0.01608819 | 4.34540785 | 12 | 0.23780877 | 0.40601471 | 0.49719006 |
| MST3 | 0.02871446 | 7.75575494 | 3 | 0.24011688 | 0.40511982 | 0.49719006 |
| CDK5 | 0.00783481 | 2.11617552 | 128 | 0.25913475 | 0.39776564 | 0.49719006 |
| LATS1 | 0.05210551 | 14.0736594 | 1 | 0.26827817 | 0.3942426 | 0.49719006 |
| BUB1 | 0.0383359 | 10.3544986 | 2 | 0.27147115 | 0.39301434 | 0.49719006 |
| CDK3 | 0.01640859 | 4.43194825 | 16 | 0.28170133 | 0.38908627 | 0.49719006 |
| EPHA5 | 0.03394583 | 9.1687437 | 3 | 0.29033813 | 0.38577879 | 0.49719006 |
| EPHA6 | 0.03394583 | 9.1687437 | 3 | 0.29033813 | 0.38577879 | 0.49719006 |
| CSNK1D | 0.00891866 | 2.40892341 | 102 | 0.29199515 | 0.38514516 | 0.49719006 |
| CHUK | 0.0093446 | 2.52396847 | 91 | 0.2983217 | 0.38272882 | 0.49719006 |
| EIF2AK2 | 0.01173002 | 3.16827002 | 45 | 0.29847489 | 0.38267037 | 0.49719006 |
| GSG2 | 0.04198822 | 11.3409888 | 2 | 0.30009948 | 0.38205064 | 0.49719006 |
| PNCK | 0.05952533 | 16.0777473 | 1 | 0.30940308 | 0.37850746 | 0.49719006 |
| KIT | 0.04379146 | 11.8280407 | 2 | 0.31423391 | 0.37667169 | 0.49719006 |

| | | | | | | |
|---|---|---|---|---|---|---|
| EPHA2 | 0.02287402 | 6.17825501 | 9 | 0.31878132 | 0.37494617 | 0.49719006 |
| CLK1 | 0.00843936 | 2.2794655 | 156 | 0.32792842 | 0.37148289 | 0.49719006 |
| CDK16 | 0.03790332 | 10.2376589 | 3 | 0.32833012 | 0.37133104 | 0.49719006 |
| EPHA1 | 0.02790428 | 7.53692519 | 6 | 0.32857712 | 0.37123767 | 0.49719006 |
| EPHA3 | 0.02790428 | 7.53692519 | 6 | 0.32857712 | 0.37123767 | 0.49719006 |
| EPHA4 | 0.02790428 | 7.53692519 | 6 | 0.32857712 | 0.37123767 | 0.49719006 |
| ROCK1 | 0.01068071 | 2.88485193 | 75 | 0.33496239 | 0.36882673 | 0.49719006 |
| PRKAA1 | 0.01354393 | 3.65820693 | 38 | 0.33625501 | 0.36833929 | 0.49719006 |
| CDK2 | 0.00724758 | 1.95756473 | 312 | 0.34708321 | 0.3642644 | 0.49719006 |
| SMG1 | 0.0480177 | 12.969545 | 2 | 0.34736081 | 0.36416014 | 0.49719006 |
| RPS6KA2 | 0.01036381 | 2.79925652 | 89 | 0.34831828 | 0.36380059 | 0.49719006 |
| TNK2 | 0.0669534 | 18.084065 | 1 | 0.35057375 | 0.36295408 | 0.49719006 |
| MAPK12 | 0.01350176 | 3.6468144 | 44 | 0.36027821 | 0.35931954 | 0.49719006 |
| PLK1 | 0.00942628 | 2.5460306 | 131 | 0.36311309 | 0.3582602 | 0.49719006 |
| BTK | 0.05161766 | 13.9418924 | 2 | 0.3755787 | 0.35361506 | 0.49719006 |
| DAPK3 | 0.0517968 | 13.990279 | 2 | 0.3769829 | 0.35309316 | 0.49719006 |
| PAK3 | 0.02486631 | 6.71637101 | 11 | 0.38904956 | 0.34861974 | 0.49719006 |
| PDGFRB | 0.03138209 | 8.47627846 | 7 | 0.40590338 | 0.3424068 | 0.49719006 |
| IRAK4 | 0.07781705 | 21.0183281 | 1 | 0.41078632 | 0.34061461 | 0.49719006 |
| FLT3 | 0.07939664 | 21.4449742 | 1 | 0.41954132 | 0.33741028 | 0.49719006 |
| MELK | 0.04774318 | 12.8953993 | 3 | 0.42279304 | 0.33622313 | 0.49719006 |
| OXSR1 | 0.05914721 | 15.9756175 | 2 | 0.43459819 | 0.33192706 | 0.49719006 |
| PDK2 | 0.02382922 | 6.43625538 | 16 | 0.44621896 | 0.32771955 | 0.49719006 |
| UHMK1 | 0.05077852 | 13.7152415 | 3 | 0.45193233 | 0.32565886 | 0.49719006 |
| RPS6KB2 | 0.02483635 | 6.70827972 | 15 | 0.45366894 | 0.32503356 | 0.49719006 |
| EPHB2 | 0.04530853 | 12.2378013 | 4 | 0.46121085 | 0.32232367 | 0.49719006 |
| MAP2K6 | 0.01729766 | 4.6720852 | 38 | 0.46450749 | 0.3211421 | 0.49719006 |
| IRR | 0.08898075 | 24.033635 | 1 | 0.47266196 | 0.31822718 | 0.49719006 |
| TXK | 0.08898075 | 24.033635 | 1 | 0.47266196 | 0.31822718 | 0.49719006 |
| TYK2 | 0.08898075 | 24.033635 | 1 | 0.47266196 | 0.31822718 | 0.49719006 |
| INSR | 0.04188302 | 11.3125739 | 5 | 0.47319511 | 0.31803699 | 0.49719006 |
| ITK | 0.04770456 | 12.8849665 | 4 | 0.48777118 | 0.31285597 | 0.49719006 |
| NME1 | 0.09226169 | 24.9198161 | 1 | 0.49084685 | 0.31176738 | 0.49719006 |
| BRSK2 | 0.09272557 | 25.0451076 | 1 | 0.4934179 | 0.31085867 | 0.49719006 |
| WEE1 | 0.09292122 | 25.0979547 | 1 | 0.49450235 | 0.31047572 | 0.49719006 |
| CSNK1E | 0.01843466 | 4.97918799 | 39 | 0.50993513 | 0.30504845 | 0.49719006 |
| HIPK2 | 0.01223446 | 3.30451907 | 118 | 0.5136999 | 0.30373091 | 0.49719006 |
| CDC42BPB | 0.03901733 | 10.5385524 | 7 | 0.5178687 | 0.30227494 | 0.49719006 |
| NUAK2 | 0.09942628 | 26.8549642 | 1 | 0.53055708 | 0.29786287 | 0.49719006 |
| SGK1 | 0.02138158 | 5.77514867 | 31 | 0.54557683 | 0.29267843 | 0.49719006 |
| PDK1 | 0.01555031 | 4.2001274 | 71 | 0.55333051 | 0.29001856 | 0.49719006 |
| GSK3A | 0.01223789 | 3.30544454 | 161 | 0.60028265 | 0.27415894 | 0.49719006 |
| FLT1 | 0.05807502 | 15.6860194 | 4 | 0.60272925 | 0.27334441 | 0.49719006 |
| YES1 | 0.03550185 | 9.58902255 | 12 | 0.61055183 | 0.27074816 | 0.49719006 |
| CDK6 | 0.02473358 | 6.68052176 | 29 | 0.62773364 | 0.26508922 | 0.49719006 |
| PRKACA | 0.01836257 | 4.95971704 | 60 | 0.62940176 | 0.26454303 | 0.49719006 |
| LYN | 0.04049727 | 10.9382838 | 10 | 0.64491068 | 0.25949254 | 0.49719006 |
| PRKCA | 0.01418279 | 3.83076009 | 131 | 0.66485492 | 0.25307165 | 0.49719006 |
| ERBB2 | 0.06383491 | 17.2417617 | 4 | 0.6665785 | 0.2525207 | 0.49719006 |
| AKT1 | 0.01310424 | 3.5394452 | 168 | 0.67543204 | 0.24970066 | 0.49719006 |
| GTF2F1 | 0.12795749 | 34.5612243 | 1 | 0.68869348 | 0.24550809 | 0.49719006 |
| CDK7 | 0.01650916 | 4.45911282 | 111 | 0.74784981 | 0.22727538 | 0.49719006 |
| BLK | 0.04882617 | 13.1879116 | 9 | 0.75030653 | 0.22653506 | 0.49719006 |
| MAPK11 | 0.02205044 | 5.95580827 | 55 | 0.75419564 | 0.22536588 | 0.49719006 |
| MARK4 | 0.03149969 | 8.50804306 | 24 | 0.75478029 | 0.22519041 | 0.49719006 |
| RAF1 | 0.01605656 | 4.33686387 | 123 | 0.75941509 | 0.22380215 | 0.49719006 |

| | | | | | | |
|---|---|---|---|---|---|---|
| PRKD2 | 0.07312862 | 19.751988 | 4 | 0.76960076 | 0.22076838 | 0.49719006 |
| CDK9 | 0.04809257 | 12.9897694 | 10 | 0.77803477 | 0.21827426 | 0.49719006 |
| MARK3 | 0.03350049 | 9.04845735 | 23 | 0.79207214 | 0.21415931 | 0.49719006 |
| NLK | 0.03364408 | 9.08724147 | 24 | 0.81300685 | 0.20810707 | 0.49719006 |
| ERBB4 | 0.10786742 | 29.1349123 | 2 | 0.81648599 | 0.20711112 | 0.49719006 |
| NEK3 | 0.10824322 | 29.2364137 | 2 | 0.8194316 | 0.20627011 | 0.49719006 |
| SLK | 0.07790038 | 21.0408367 | 4 | 0.82249642 | 0.20539721 | 0.49719006 |
| HCK | 0.05284162 | 14.2724839 | 10 | 0.86127211 | 0.1945441 | 0.49719006 |
| DYRK3 | 0.16007434 | 43.2359617 | 1 | 0.86670352 | 0.19305224 | 0.49719006 |
| CHEK2 | 0.05896943 | 15.9276011 | 10 | 0.96867523 | 0.16635363 | 0.49719006 |
| AKT2 | 0.02988076 | 8.07077063 | 45 | 0.9733324 | 0.16519406 | 0.49719006 |
| MAPK8 | 0.02040141 | 5.51040649 | 112 | 0.97951901 | 0.1636618 | 0.49719006 |
| MAK | 0.06177865 | 16.6863685 | 10 | 1.01791283 | 0.15435969 | 0.49719006 |
| SGK2 | 0.13494022 | 36.4472547 | 2 | 1.02869298 | 0.15181198 | 0.49719006 |
| MAPK6 | 0.19296099 | 52.1186216 | 1 | 1.04898024 | 0.14709361 | 0.49719006 |
| AKT3 | 0.03968178 | 10.7180187 | 28 | 1.05522463 | 0.1456613 | 0.49719006 |
| MAPK13 | 0.03210672 | 8.67200281 | 45 | 1.05609548 | 0.1454623 | 0.49719006 |
| GRK4 | 0.1146277 | 30.9608568 | 3 | 1.06488578 | 0.14346381 | 0.49719006 |
| SGK3 | 0.10750855 | 29.0379807 | 4 | 1.15070739 | 0.12492632 | 0.46762959 |
| CSNK1G1 | 0.0528477 | 14.2741262 | 18 | 1.15566076 | 0.12390997 | 0.46762959 |
| CDK4 | 0.0339992 | 9.18315973 | 52 | 1.21090724 | 0.11296548 | 0.4346033 |
| PAK4 | 0.04218781 | 11.3948968 | 34 | 1.24379216 | 0.10678803 | 0.42627005 |
| NUAK1 | 0.11661567 | 31.4978077 | 4 | 1.25166121 | 0.10534667 | 0.42627005 |
| SRPK1 | 0.16867511 | 45.559025 | 2 | 1.29312007 | 0.09798477 | 0.4051012 |
| MAP4K4 | 0.12072689 | 32.6082436 | 5 | 1.45035241 | 0.07348014 | 0.35708768 |
| PTK2 | 0.27571777 | 74.4711685 | 1 | 1.50766593 | 0.06582002 | 0.32557405 |
| EGFR | 0.09225381 | 24.917688 | 10 | 1.55205594 | 0.06032442 | 0.30784897 |
| AURKC | 0.08227882 | 22.2234482 | 13 | 1.57027529 | 0.05817554 | 0.30784897 |
| MAP2K4 | 0.04491143 | 12.1305442 | 53 | 1.66280953 | 0.04817528 | 0.28392667 |
| ABL1 | 0.08853273 | 23.9126247 | 13 | 1.69525366 | 0.04501366 | 0.27244938 |
| MET | 0.10570915 | 28.5519632 | 10 | 1.7878897 | 0.0368969 | 0.24927907 |
| AURKB | 0.04155063 | 11.2227952 | 73 | 1.79233532 | 0.03653963 | 0.24927907 |
| DAPK1 | 0.08223364 | 22.2112473 | 20 | 1.9465671 | 0.02579333 | 0.18801979 |
| SRRM1 | 0.21590459 | 58.3156725 | 3 | 2.03714618 | 0.0208177 | 0.16960301 |
| IGF1R | 0.21680535 | 58.5589668 | 3 | 2.04579348 | 0.02038835 | 0.16960301 |
| RPS6KA1 | 0.09112853 | 24.6137489 | 19 | 2.11217509 | 0.01733572 | 0.1500623 |
| EPHB1 | 0.17589764 | 47.5098227 | 5 | 2.13411521 | 0.01641668 | 0.1500623 |
| TRKB | 0.24390872 | 65.8795683 | 3 | 2.30598646 | 0.01055569 | 0.11695709 |
| FGR | 0.125199 | 33.8161597 | 12 | 2.33274117 | 0.00983087 | 0.11346459 |
| NEK2 | 0.0307777 | 8.31303297 | 249 | 2.36802007 | 0.00894178 | 0.10769017 |
| CDK1 | 0.02059561 | 5.56285977 | 654 | 2.3944963 | 0.00832161 | 0.10477658 |
| TTK | 0.04436109 | 11.9818985 | 114 | 2.40612412 | 0.00806139 | 0.10477658 |
| MAP2K7 | 0.09204338 | 24.8608484 | 25 | 2.44818402 | 0.00717892 | 0.09942798 |
| KDR | 0.26396842 | 71.2976775 | 3 | 2.49855997 | 0.00623495 | 0.09558412 |
| CSNK1G3 | 0.14641072 | 39.5454282 | 10 | 2.50127273 | 0.00618739 | 0.09558412 |
| TRKA | 0.32616477 | 88.0968654 | 2 | 2.52758458 | 0.00574251 | 0.09558412 |
| TRKC | 0.32616477 | 88.0968654 | 2 | 2.52758458 | 0.00574251 | 0.09558412 |
| FYN | 0.2531669 | 68.380194 | 4 | 2.76535204 | 0.00284307 | 0.06562753 |
| AURKA | 0.03766938 | 10.1744715 | 226 | 2.83024121 | 0.00232565 | 0.058564 |
| PAK2 | 0.07295305 | 19.7045665 | 60 | 2.97311322 | 0.00147398 | 0.04536578 |
| RPS6KB1 | 0.08148966 | 22.0102977 | 48 | 2.98703972 | 0.00140847 | 0.04536578 |
| SRC | 0.16388966 | 44.2664775 | 12 | 3.07560342 | 0.00105039 | 0.04156525 |
| PTK6 | 0.69825405 | 188.597908 | 1 | 3.84960497 | 5.92E-05 | 0.00546191 |
| ULK1 | 0.74873967 | 202.234037 | 1 | 4.12942529 | 1.82E-05 | 0.00251842 |
| PKM | 1.16813101 | 315.511333 | 1 | 6.45393332 | 5.45E-11 | 1.51E-08 |
