## Supplementary table 6 for "MAP4K inhibition by the CNS penetrant inhibitor famlasertib restrains medulloblastoma dissemination without developmental toxicity"

| Kinase.Gene | mS | Enrichment | m | z.score | p.value | FDR |
|---|---|---|---|---|---|---|
| MAPK15 | -0.6131029 | -289.32343 | 1 | -3.7106976 | 0.00010334 | 0.0071566 |
| ILK | -0.528482 | -249.39084 | 1 | -3.2003091 | 0.0006864 | 0.0318103 |
| NEK1 | -0.09996 | -47.171141 | 27 | -3.1992066 | 0.00068903 | 0.0318103 |
| STK11 | -0.1656142 | -78.153377 | 9 | -3.0350383 | 0.00120253 | 0.04758565 |
| PRKCB | -0.0281835 | -13.299784 | 243 | -2.8490896 | 0.00219223 | 0.06072467 |
| PRKD1 | -0.1217792 | -57.467626 | 14 | -2.7961017 | 0.00258616 | 0.06222414 |
| PKD2/PRKD2 | -0.2617507 | -123.52026 | 3 | -2.7566023 | 0.00292027 | 0.06222414 |
| PBK | -0.2972743 | -140.28383 | 2 | -2.5537651 | 0.00532826 | 0.09238222 |
| CAMK2B | -0.0809009 | -38.177167 | 26 | -2.5532491 | 0.00533616 | 0.09238222 |
| PLK3 | -0.2309777 | -108.99844 | 3 | -2.4351215 | 0.0074434 | 0.11392299 |
| ATM | -0.0470195 | -22.188495 | 63 | -2.3524279 | 0.00932565 | 0.1230098 |
| MKNK1 | -0.2629418 | -124.08232 | 2 | -2.2609159 | 0.01188223 | 0.13165513 |
| MKNK2 | -0.2629418 | -124.08232 | 2 | -2.2609159 | 0.01188223 | 0.13165513 |
| MOK | -0.1801696 | -85.022102 | 4 | -2.1989408 | 0.01394107 | 0.14852599 |
| CHEK1 | -0.0549694 | -25.940101 | 39 | -2.1503294 | 0.01576458 | 0.16173294 |
| RPS6KA3 | -0.0604328 | -28.518265 | 28 | -1.9963805 | 0.02294626 | 0.20750408 |
| BRAF | -0.1610515 | -76.000242 | 4 | -1.9683194 | 0.02451565 | 0.20750408 |
| CDKL5 | -0.2872243 | -135.54123 | 1 | -1.7451682 | 0.0404778 | 0.28030873 |
| DMPK | -0.0750899 | -35.434965 | 13 | -1.6790496 | 0.04657119 | 0.29318682 |
| ATR | -0.0483099 | -22.797459 | 27 | -1.5804689 | 0.05699976 | 0.32222316 |
| PRKACB | -0.0455893 | -21.513618 | 24 | -1.4096925 | 0.07931525 | 0.41453443 |
| GSK3B | -0.0148059 | -6.9869035 | 172 | -1.3388014 | 0.09031767 | 0.43891221 |
| ROCK2 | -0.0331703 | -15.653055 | 34 | -1.2411003 | 0.10728436 | 0.44622467 |
| PRKACG | -0.0374393 | -17.667619 | 26 | -1.2166036 | 0.11187754 | 0.44622467 |
| LIMK1 | -0.1966583 | -92.803103 | 1 | -1.1989213 | 0.11527927 | 0.44622467 |
| LIMK2 | -0.1966583 | -92.803103 | 1 | -1.1989213 | 0.11527927 | 0.44622467 |
| TESK1 | -0.1966583 | -92.803103 | 1 | -1.1989213 | 0.11527927 | 0.44622467 |
| PLK1 | -0.013521 | -6.3805859 | 131 | -1.0796922 | 0.14013963 | 0.48523348 |
| EEF2K | -0.1183228 | -55.836546 | 2 | -1.0273446 | 0.1521291 | 0.49786934 |
| PRKDC | -0.0137829 | -6.5041781 | 113 | -1.0195672 | 0.15396689 | 0.49786934 |
| CDK3 | -0.0400172 | -18.884128 | 16 | -1.0165764 | 0.1546775 | 0.49786934 |
| CSF1R | -0.0658865 | -31.091871 | 6 | -1.0047179 | 0.15751635 | 0.49786934 |
| WEE1 | -0.1629912 | -76.915613 | 1 | -0.9958593 | 0.15965926 | 0.49786934 |
| VRK1 | -0.0897087 | -42.333566 | 3 | -0.959309 | 0.16870154 | 0.49786934 |
| CAMK2A | -0.0257943 | -12.172338 | 31 | -0.9373832 | 0.17428073 | 0.49786934 |
| TEC | -0.1466267 | -69.193196 | 1 | -0.8971571 | 0.18481755 | 0.49786934 |
| PRKG1 | -0.018225 | -8.6003691 | 52 | -0.8848366 | 0.18812238 | 0.49786934 |
| CDC42BPA | -0.0364128 | -17.1832 | 14 | -0.8695765 | 0.19226594 | 0.49786934 |
| WNK1 | -0.0981481 | -46.31614 | 2 | -0.8552593 | 0.19620382 | 0.49786934 |
| KIT | -0.097768 | -46.136741 | 2 | -0.8520165 | 0.19710245 | 0.49786934 |
| MAP4K6 | -0.0681599 | -32.164662 | 4 | -0.847772 | 0.19828249 | 0.49786934 |
| PRKCQ | -0.0660462 | -31.167229 | 4 | -0.8222751 | 0.20546018 | 0.49786934 |
| PIM2 | -0.0511595 | -24.142201 | 6 | -0.7871408 | 0.21559974 | 0.49786934 |
| EPHA1 | -0.0496459 | -23.427909 | 6 | -0.764778 | 0.22220187 | 0.49786934 |
| EPHA3 | -0.0496459 | -23.427909 | 6 | -0.764778 | 0.22220187 | 0.49786934 |
| EPHA4 | -0.0496459 | -23.427909 | 6 | -0.764778 | 0.22220187 | 0.49786934 |
| MARK2 | -0.0479071 | -22.607386 | 6 | -0.7390894 | 0.22992634 | 0.49786934 |
| EPHA7 | -0.0518745 | -24.479583 | 5 | -0.7282002 | 0.23324551 | 0.49786934 |
| PRKD2 | -0.0578407 | -27.295042 | 4 | -0.7232923 | 0.23475017 | 0.49786934 |
| CSNK2A2 | -0.0125607 | -5.927405 | 61 | -0.6915267 | 0.24461729 | 0.49786934 |
| PRKCI | -0.015062 | -7.1077857 | 40 | -0.6553989 | 0.25610546 | 0.49786934 |
| TLK2 | -0.0417471 | -19.700466 | 6 | -0.6480809 | 0.25846631 | 0.49786934 |
| MAPKAPK2 | -0.025622 | -12.091049 | 15 | -0.6480277 | 0.25848352 | 0.49786934 |
| CSNK1D | -0.0074979 | -3.5382408 | 102 | -0.5858158 | 0.27899964 | 0.49786934 |

| | | | | | | |
|---|---|---|---|---|---|---|
| PIM3 | -0.0400609 | -18.904764 | 5 | -0.5688733 | 0.28472107 | 0.49786934 |
| TLK1 | -0.0356127 | -16.805627 | 6 | -0.5574506 | 0.2886098 | 0.49786934 |
| IGF1R | -0.0481394 | -22.717002 | 3 | -0.5250418 | 0.29977706 | 0.49786934 |
| AAK1 | -0.0756362 | -35.692753 | 1 | -0.4689794 | 0.31954217 | 0.49786934 |
| PLK2 | -0.0292962 | -13.824898 | 6 | -0.4641312 | 0.32127686 | 0.49786934 |
| PDHK3 | -0.0505492 | -23.854195 | 2 | -0.4492501 | 0.32662563 | 0.49786934 |
| GRK1 | -0.0226577 | -10.692198 | 9 | -0.4483229 | 0.3269601 | 0.49786934 |
| MAP2K5 | -0.0151192 | -7.1347764 | 18 | -0.4411186 | 0.32956358 | 0.49786934 |
| CAMK2G | -0.0143802 | -6.7860428 | 16 | -0.3980616 | 0.3452924 | 0.49786934 |
| IRAK4 | -0.0635575 | -29.992812 | 1 | -0.396127 | 0.34600566 | 0.49786934 |
| MAP3K5 | -0.0327523 | -15.455818 | 3 | -0.3642953 | 0.35781875 | 0.49786934 |
| MAP3K6 | -0.0327523 | -15.455818 | 3 | -0.3642953 | 0.35781875 | 0.49786934 |
| CSNK1A1 | -0.0045746 | -2.1587723 | 76 | -0.3519643 | 0.36243252 | 0.49786934 |
| PRKCA | -0.0029608 | -1.3971806 | 131 | -0.3506791 | 0.36291455 | 0.49786934 |
| PRKCD | -0.0059992 | -2.8310055 | 50 | -0.3462352 | 0.36458297 | 0.49786934 |
| EPHB2 | -0.0262865 | -12.404631 | 4 | -0.342656 | 0.36592863 | 0.49786934 |
| MAPKAPK3 | -0.0371469 | -17.52961 | 2 | -0.3349306 | 0.36883874 | 0.49786934 |
| EPHA2 | -0.016295 | -7.6895962 | 9 | -0.3331918 | 0.36949475 | 0.49786934 |
| CDK14 | -0.0355413 | -16.77197 | 2 | -0.3212359 | 0.37401582 | 0.49786934 |
| MST1 | -0.051027 | -24.079659 | 1 | -0.3205495 | 0.3742759 | 0.49786934 |
| PKD1 | -0.0066073 | -3.117979 | 31 | -0.2930478 | 0.38474281 | 0.49786934 |
| CDK2 | -0.0005945 | -0.2805525 | 312 | -0.2891 | 0.38625241 | 0.49786934 |
| LYN | -0.0119143 | -5.6223807 | 10 | -0.2676626 | 0.39447954 | 0.49786934 |
| WNK3 | -0.0413147 | -19.496436 | 1 | -0.2619702 | 0.39667222 | 0.49786934 |
| EPHA5 | -0.0222144 | -10.482996 | 3 | -0.2542081 | 0.39966741 | 0.49786934 |
| EPHA6 | -0.0222144 | -10.482996 | 3 | -0.2542081 | 0.39966741 | 0.49786934 |
| FLT3 | -0.0385029 | -18.169537 | 1 | -0.2450107 | 0.40322406 | 0.49786934 |
| PLK4 | -0.0046733 | -2.2053163 | 30 | -0.2243908 | 0.41122662 | 0.49786934 |
| ACTR2 | -0.0179292 | -8.4608156 | 3 | -0.2094415 | 0.41705181 | 0.49786934 |
| CDK5 | -0.0008509 | -0.4015362 | 128 | -0.2026669 | 0.4196977 | 0.49786934 |
| GSG2 | -0.0199875 | -9.4321274 | 2 | -0.1885651 | 0.42521684 | 0.49786934 |
| ADRBK2 | -0.0117083 | -5.5251433 | 5 | -0.186487 | 0.42603144 | 0.49786934 |
| PDPK1 | -0.0154494 | -7.2905975 | 3 | -0.1835354 | 0.42718896 | 0.49786934 |
| CLK3 | -0.0133427 | -6.2964407 | 3 | -0.161527 | 0.43583917 | 0.49786934 |
| IRAK1 | -0.0243312 | -11.481904 | 1 | -0.1595344 | 0.43662393 | 0.49786934 |
| NEK6 | -0.0243312 | -11.481904 | 1 | -0.1595344 | 0.43662393 | 0.49786934 |
| TAF1 | -0.0239065 | -11.28147 | 1 | -0.1569726 | 0.43763322 | 0.49786934 |
| MAPK10 | -0.0021783 | -1.0279352 | 34 | -0.1511357 | 0.43993434 | 0.49786934 |
| GRK7 | -0.0089849 | -4.2399729 | 5 | -0.1497571 | 0.44047811 | 0.49786934 |
| CSNK1E | -0.0016375 | -0.7727387 | 39 | -0.1414981 | 0.44373824 | 0.49786934 |
| RIPK1 | -0.0185422 | -8.7500795 | 1 | -0.1246182 | 0.45041289 | 0.49786934 |
| MAP3K10 | -0.0175709 | -8.291717 | 1 | -0.1187598 | 0.45273283 | 0.49786934 |
| GRK5 | -0.0050559 | -2.3859 | 6 | -0.1060042 | 0.45778949 | 0.49786934 |
| MAPK14 | -0.0002542 | -0.1199506 | 46 | -0.0970848 | 0.46132952 | 0.49786934 |
| TTK | 0.00083091 | 0.3921066 | 114 | -0.0829571 | 0.46694284 | 0.49786934 |
| MAP3K1 | -0.0101293 | -4.7800397 | 1 | -0.0738762 | 0.47055446 | 0.49786934 |
| STK25 | -0.0094012 | -4.4364344 | 1 | -0.0694845 | 0.472302 | 0.49786934 |
| STK10 | -0.0040325 | -1.9029253 | 2 | -0.0524716 | 0.47907646 | 0.49786934 |
| PDK3 | -0.006476 | -3.0560444 | 1 | -0.0518413 | 0.47932756 | 0.49786934 |
| PDK4 | -0.006476 | -3.0560444 | 1 | -0.0518413 | 0.47932756 | 0.49786934 |
| GSK3A | 0.00144656 | 0.6826324 | 161 | -0.0514694 | 0.47947575 | 0.49786934 |
| PAK6 | -0.0020435 | -0.9643146 | 4 | -0.0502128 | 0.4799764 | 0.49786934 |
| PRKAA1 | 0.00111411 | 0.52574757 | 38 | -0.0373658 | 0.48509665 | 0.49786934 |
| CAMK1D | -0.0001544 | -0.0728804 | 6 | -0.0335893 | 0.48660234 | 0.49786934 |
| ULK1 | 0 | 0 | 1 | -0.0127813 | 0.49490116 | 0.49786934 |
| ADRBK1 | 0.0018905 | 0.89212717 | 51 | -0.0098462 | 0.49607198 | 0.49786934 |

| | | | | | | |
|---|---|---|---|---|---|---|
| MELK | 0.00216642 | 1.02233585 | 3 | 0.00049447 | 0.49980274 | 0.49980274 |
| OXSR1 | 0.00343963 | 1.62316484 | 2 | 0.01126397 | 0.49550642 | 0.49786934 |
| MAP2K3 | 0.00262304 | 1.2378155 | 25 | 0.0151979 | 0.49393715 | 0.49786934 |
| CSNK2A1 | 0.0022896 | 1.08046197 | 269 | 0.01686709 | 0.49327132 | 0.49786934 |
| WNK2 | 0.00542809 | 2.56151691 | 1 | 0.01995814 | 0.49203838 | 0.49786934 |
| PAK5 | 0.00361278 | 1.70487092 | 6 | 0.02206778 | 0.49119694 | 0.49786934 |
| ROCK1 | 0.00257717 | 1.216165 | 75 | 0.02392707 | 0.49045539 | 0.49786934 |
| PNCK | 0.00793496 | 3.74451249 | 1 | 0.03507831 | 0.48600865 | 0.49786934 |
| MAPK7 | 0.00310317 | 1.46438614 | 54 | 0.04361638 | 0.4826051 | 0.49786934 |
| CAMK2D | 0.0036838 | 1.7383852 | 28 | 0.0499385 | 0.4800857 | 0.49786934 |
| PDGFRB | 0.00658207 | 3.10608064 | 7 | 0.07121927 | 0.47161162 | 0.49786934 |
| CLK4 | 0.01305115 | 6.15884345 | 2 | 0.09324828 | 0.46285316 | 0.49786934 |
| TRKB | 0.01293397 | 6.10354311 | 3 | 0.11298113 | 0.45502276 | 0.49786934 |
| CDK19 | 0.0210935 | 9.95402977 | 1 | 0.11444373 | 0.45444302 | 0.49786934 |
| TGFBR2 | 0.00666771 | 3.1464944 | 20 | 0.12269256 | 0.45117528 | 0.49786934 |
| MAP3K7 | 0.01832443 | 8.647305 | 2 | 0.13822828 | 0.44503 | 0.49786934 |
| SLK | 0.01520825 | 7.17677927 | 4 | 0.15789398 | 0.43727017 | 0.49786934 |
| MAP2K6 | 0.00637261 | 3.00723884 | 38 | 0.15814823 | 0.43717 | 0.49786934 |
| CAMK1G | 0.02258983 | 10.6601493 | 2 | 0.17461129 | 0.43069254 | 0.49786934 |
| TRPM7 | 0.01315088 | 6.20590428 | 7 | 0.17604297 | 0.43013009 | 0.49786934 |
| AURKC | 0.01056739 | 4.98675522 | 13 | 0.18372351 | 0.42711518 | 0.49786934 |
| BUB1 | 0.02386727 | 11.2629739 | 2 | 0.1855076 | 0.42641546 | 0.49786934 |
| BLK | 0.01247661 | 5.8877174 | 9 | 0.18741348 | 0.42566823 | 0.49786934 |
| STK39 | 0.0241218 | 11.3830874 | 2 | 0.1876787 | 0.42556426 | 0.49786934 |
| MST3 | 0.0204431 | 9.64710313 | 3 | 0.19142769 | 0.42409527 | 0.49786934 |
| MAPK9 | 0.00756535 | 3.57008985 | 42 | 0.21288567 | 0.41570807 | 0.49786934 |
| STK3 | 0.03764772 | 17.7659714 | 1 | 0.21429015 | 0.4151604 | 0.49786934 |
| STK4 | 0.03764772 | 17.7659714 | 1 | 0.21429015 | 0.4151604 | 0.49786934 |
| CAMK4 | 0.01059548 | 5.00000933 | 18 | 0.21690559 | 0.41414096 | 0.49786934 |
| BRSK1 | 0.02809276 | 13.256983 | 2 | 0.22155016 | 0.41233204 | 0.49786934 |
| ITK | 0.02052452 | 9.68552711 | 4 | 0.22202386 | 0.41214765 | 0.49786934 |
| LMTK2 | 0.03903026 | 18.4183944 | 1 | 0.22262893 | 0.41191216 | 0.49786934 |
| CDK16 | 0.02565096 | 12.1046939 | 3 | 0.24583329 | 0.40290564 | 0.49786934 |
| UHMK1 | 0.02610034 | 12.3167596 | 3 | 0.25052796 | 0.40108954 | 0.49786934 |
| TRPM6 | 0.01465363 | 6.91505128 | 12 | 0.26189223 | 0.39670226 | 0.49786934 |
| MAPK1 | 0.00479767 | 2.26402339 | 272 | 0.26644836 | 0.39494696 | 0.49786934 |
| AURKA | 0.00510228 | 2.40776719 | 226 | 0.27049456 | 0.3933899 | 0.49786934 |
| HIPK2 | 0.00629778 | 2.97192256 | 118 | 0.27378165 | 0.39212621 | 0.49786934 |
| NEK2 | 0.00510745 | 2.41020883 | 249 | 0.28441768 | 0.38804516 | 0.49786934 |
| PKD3 | 0.02351195 | 11.0952944 | 5 | 0.28852103 | 0.38647397 | 0.49786934 |
| CDC7 | 0.02666308 | 12.5823168 | 4 | 0.29607308 | 0.38358714 | 0.49786934 |
| TRKA | 0.03865241 | 18.2400833 | 2 | 0.3116218 | 0.37766398 | 0.49786934 |
| TRKC | 0.03865241 | 18.2400833 | 2 | 0.3116218 | 0.37766398 | 0.49786934 |
| DYRK1A | 0.02530074 | 11.939429 | 5 | 0.31264619 | 0.37727474 | 0.49786934 |
| BMX | 0.03999172 | 18.8721046 | 2 | 0.32304586 | 0.37333026 | 0.49786934 |
| DYRK2 | 0.0214622 | 10.1280189 | 8 | 0.32998561 | 0.37070542 | 0.49786934 |
| CDK20 | 0.05817466 | 27.4526423 | 1 | 0.33809795 | 0.36764469 | 0.49786934 |
| ACVR1B | 0.0589855 | 27.835277 | 1 | 0.3429885 | 0.36580355 | 0.49786934 |
| SMG1 | 0.04362063 | 20.5845912 | 2 | 0.35399978 | 0.36166952 | 0.49786934 |
| CSNK1G2 | 0.01573788 | 7.426709 | 19 | 0.35804607 | 0.36015442 | 0.49786934 |
| PAK4 | 0.01232971 | 5.81839316 | 34 | 0.35909981 | 0.35976021 | 0.49786934 |
| NLK | 0.01457193 | 6.87649927 | 24 | 0.36795761 | 0.35645242 | 0.49786934 |
| MAPK3 | 0.00569296 | 2.68650923 | 308 | 0.37830106 | 0.35260348 | 0.49786934 |
| MARK4 | 0.0153893 | 7.26221393 | 24 | 0.39210917 | 0.34748878 | 0.49786934 |
| ABL1 | 0.02071283 | 9.77439072 | 13 | 0.40435436 | 0.34297608 | 0.49786934 |
| CDK4 | 0.01234164 | 5.82402373 | 52 | 0.44461549 | 0.32829883 | 0.49786934 |

| | | | | | | |
|---|---|---|---|---|---|---|
| MAP3K8 | 0.04541962 | 21.4335338 | 3 | 0.45235312 | 0.32550731 | 0.49786934 |
| HCK | 0.02610309 | 12.3180552 | 10 | 0.45745174 | 0.32367319 | 0.49786934 |
| MAP2K2 | 0.01621587 | 7.65227243 | 29 | 0.45787033 | 0.3235228 | 0.49786934 |
| TNK2 | 0.08027543 | 37.8820017 | 1 | 0.47139825 | 0.31867818 | 0.49786934 |
| MAPK8 | 0.00956249 | 4.5125409 | 112 | 0.47512062 | 0.3173505 | 0.49786934 |
| CDK1 | 0.00521596 | 2.46141542 | 654 | 0.47767888 | 0.31643939 | 0.49786934 |
| LATS1 | 0.08240248 | 38.8857539 | 1 | 0.48422746 | 0.31411222 | 0.49786934 |
| NUAK2 | 0.08241626 | 38.8922614 | 1 | 0.48431064 | 0.31408271 | 0.49786934 |
| DAPK2 | 0.08317881 | 39.2521062 | 1 | 0.4889099 | 0.31245274 | 0.49786934 |
| DYRK1B | 0.08451608 | 39.8831667 | 1 | 0.49697565 | 0.30960311 | 0.49786934 |
| BRSK2 | 0.08738716 | 41.2380269 | 1 | 0.51429246 | 0.30352377 | 0.49786934 |
| CDK6 | 0.01812986 | 8.55548577 | 29 | 0.52003775 | 0.30151863 | 0.49786934 |
| NEK3 | 0.06438539 | 30.3834847 | 2 | 0.53111893 | 0.29766818 | 0.49786934 |
| PAK1 | 0.0147872 | 6.9780822 | 50 | 0.54028183 | 0.29450134 | 0.49786934 |
| ICK | 0.02919747 | 13.7782932 | 11 | 0.54167984 | 0.29401954 | 0.49786934 |
| PDHK4 | 0.02813873 | 13.2786771 | 12 | 0.54364535 | 0.29334277 | 0.49786934 |
| PDK2 | 0.02471744 | 11.6641675 | 16 | 0.54520575 | 0.29280601 | 0.49786934 |
| GRK6 | 0.05454088 | 25.7378583 | 3 | 0.54764132 | 0.29196911 | 0.49786934 |
| FLT1 | 0.04755547 | 22.4414418 | 4 | 0.54809705 | 0.29181263 | 0.49786934 |
| BCR | 0.06705246 | 31.6420768 | 2 | 0.55386851 | 0.28983442 | 0.49786934 |
| TBK1 | 0.06808521 | 32.1294309 | 2 | 0.56267764 | 0.28682721 | 0.49786934 |
| MAPKAPK5 | 0.04062972 | 19.1731769 | 6 | 0.56895768 | 0.28469243 | 0.49786934 |
| PRKCG | 0.01389597 | 6.55751042 | 68 | 0.5857445 | 0.27902361 | 0.49786934 |
| IKBKB | 0.01275153 | 6.01745064 | 86 | 0.59471079 | 0.27601841 | 0.49786934 |
| CDC42BPB | 0.03976095 | 18.7632055 | 7 | 0.60068093 | 0.27402626 | 0.49786934 |
| PRKG2 | 0.02131106 | 10.056695 | 27 | 0.60148542 | 0.27375836 | 0.49786934 |
| CDK7 | 0.01163209 | 5.48918766 | 111 | 0.60450894 | 0.27275266 | 0.49786934 |
| CDK9 | 0.03428423 | 16.1787381 | 10 | 0.61349235 | 0.26977542 | 0.49786934 |
| YES1 | 0.032622 | 15.3943304 | 12 | 0.6373171 | 0.26195916 | 0.49786934 |
| PAK7 | 0.10947585 | 51.66169 | 1 | 0.64751995 | 0.25864774 | 0.49786934 |
| NUAK1 | 0.05653927 | 26.6809001 | 4 | 0.65646824 | 0.25576145 | 0.49786934 |
| AURKB | 0.01514358 | 7.14625985 | 73 | 0.67119054 | 0.25104958 | 0.49786934 |
| MAK | 0.03752511 | 17.708111 | 10 | 0.67530635 | 0.24974058 | 0.49786934 |
| ACVR2B | 0.06697866 | 31.6072516 | 3 | 0.67757667 | 0.24902007 | 0.49786934 |
| PRKACA | 0.01697383 | 8.00995558 | 60 | 0.69400783 | 0.24383865 | 0.49786934 |
| MAP2K1 | 0.01600194 | 7.55131959 | 69 | 0.69554753 | 0.24335612 | 0.49786934 |
| MAP4K4 | 0.05473547 | 25.8296834 | 5 | 0.70962624 | 0.23896797 | 0.49786934 |
| PRKCZ | 0.02330943 | 10.9997285 | 32 | 0.72299731 | 0.23484077 | 0.49786934 |
| PIM1 | 0.03162154 | 14.922214 | 18 | 0.75494974 | 0.22513957 | 0.49786934 |
| ERBB4 | 0.09291318 | 43.8457604 | 2 | 0.77445526 | 0.21933081 | 0.49786934 |
| SRPK1 | 0.09368178 | 44.2084611 | 2 | 0.78101123 | 0.21739795 | 0.49786934 |
| CAMK1 | 0.03565948 | 16.8277191 | 15 | 0.78349719 | 0.2166676 | 0.49786934 |
| CHEK2 | 0.04333714 | 20.4508123 | 10 | 0.78616051 | 0.21588673 | 0.49786934 |
| MARK1 | 0.02851382 | 13.4556823 | 25 | 0.79599623 | 0.21301711 | 0.49786934 |
| MAPK6 | 0.13448922 | 63.4655034 | 1 | 0.79838749 | 0.21232283 | 0.49786934 |
| MARK3 | 0.0304709 | 14.379225 | 23 | 0.82010292 | 0.20607872 | 0.49786934 |
| SGK2 | 0.09875265 | 46.6014064 | 2 | 0.82426473 | 0.20489458 | 0.49786934 |
| HIPK1 | 0.09970419 | 47.0504368 | 2 | 0.83238114 | 0.20259692 | 0.49786934 |
| CLK1 | 0.01345016 | 6.34713381 | 156 | 0.85360477 | 0.19666201 | 0.49786934 |
| IRR | 0.14874136 | 70.1910952 | 1 | 0.88434899 | 0.18825392 | 0.49786934 |
| TXK | 0.14874136 | 70.1910952 | 1 | 0.88434899 | 0.18825392 | 0.49786934 |
| TYK2 | 0.14874136 | 70.1910952 | 1 | 0.88434899 | 0.18825392 | 0.49786934 |
| JAK2 | 0.10637696 | 50.1993186 | 2 | 0.88929851 | 0.18692133 | 0.49786934 |
| DAPK1 | 0.03516462 | 16.5941931 | 20 | 0.8913564 | 0.186369 | 0.49786934 |
| RET | 0.08758113 | 41.3295651 | 3 | 0.89280712 | 0.18598024 | 0.49786934 |
| EPHB1 | 0.07500803 | 35.3963109 | 5 | 0.9830381 | 0.16279434 | 0.49786934 |

| | | | | | | |
|---|---|---|---|---|---|---|
| PRKAA2 | 0.02089512 | 9.86041421 | 77 | 0.99374022 | 0.16017468 | 0.49786934 |
| DYRK3 | 0.16722134 | 78.911804 | 1 | 0.99581059 | 0.15967109 | 0.49786934 |
| SGK1 | 0.03178382 | 14.9987967 | 31 | 0.99619655 | 0.15957733 | 0.49786934 |
| CSNK1G1 | 0.04560549 | 21.5212451 | 18 | 1.11279058 | 0.13289919 | 0.4659883 |
| MOS | 0.19054746 | 89.9194069 | 1 | 1.13650157 | 0.12787336 | 0.45411435 |
| INSR | 0.08776002 | 41.4139798 | 5 | 1.15502159 | 0.12404079 | 0.44622467 |
| PAK3 | 0.06019452 | 28.4058141 | 11 | 1.1617496 | 0.1226686 | 0.44622467 |
| BTK | 0.13881702 | 65.507795 | 2 | 1.16600571 | 0.12180607 | 0.44622467 |
| VRK2 | 0.19562599 | 92.3159668 | 1 | 1.16713261 | 0.12157841 | 0.44622467 |
| VRK3 | 0.19562599 | 92.3159668 | 1 | 1.16713261 | 0.12157841 | 0.44622467 |
| NEK4 | 0.07103684 | 33.5223067 | 8 | 1.17570893 | 0.11985561 | 0.44622467 |
| PRKCT | 0.03041046 | 14.3507052 | 49 | 1.19447132 | 0.11614683 | 0.44622467 |
| NME1 | 0.20072781 | 94.7235164 | 1 | 1.19790411 | 0.11547717 | 0.44622467 |
| PDK1 | 0.026206 | 12.3666216 | 71 | 1.22414837 | 0.11044813 | 0.44622467 |
| ARAF | 0.03901017 | 18.408913 | 31 | 1.23887071 | 0.10769669 | 0.44622467 |
| GRK4 | 0.12191316 | 57.5308607 | 3 | 1.25146788 | 0.10538191 | 0.44622467 |
| MTOR | 0.0450824 | 21.2743985 | 24 | 1.26948356 | 0.10213433 | 0.44622467 |
| EIF2AK2 | 0.03427427 | 16.1740367 | 45 | 1.3010107 | 0.0966274 | 0.44609648 |
| NEK11 | 0.11128117 | 52.5136183 | 4 | 1.31681733 | 0.09394993 | 0.44108697 |
| SGK3 | 0.11170011 | 52.7113171 | 4 | 1.32187101 | 0.09310555 | 0.44108697 |
| CSNK2B | 0.1598118 | 75.4152405 | 2 | 1.34508698 | 0.08929857 | 0.43891221 |
| BCKDK | 0.0820769 | 38.7321159 | 8 | 1.36404794 | 0.08627624 | 0.43451851 |
| DAPK3 | 0.164388 | 77.5747486 | 2 | 1.384121 | 0.0831607 | 0.42658361 |
| LCK | 0.10688527 | 50.4391906 | 5 | 1.4129599 | 0.07883376 | 0.41453443 |
| AKT3 | 0.0482726 | 22.7798557 | 28 | 1.47301611 | 0.07037335 | 0.38222386 |
| AKT2 | 0.03880728 | 18.3131687 | 45 | 1.4844183 | 0.06884899 | 0.38142341 |
| CLK2 | 0.0659612 | 31.1271121 | 17 | 1.58765248 | 0.05618247 | 0.32222316 |
| PTK2 | 0.27143735 | 128.091371 | 1 | 1.62438715 | 0.0521466 | 0.30733211 |
| AKT1 | 0.02327448 | 10.9832346 | 168 | 1.65386249 | 0.04907773 | 0.29553328 |
| SRRM1 | 0.16088855 | 75.9233573 | 3 | 1.65863696 | 0.04859448 | 0.29553328 |
| PRKCH | 0.05387147 | 25.4219657 | 30 | 1.70967959 | 0.04366257 | 0.28126818 |
| MAPK11 | 0.04045354 | 19.0900363 | 55 | 1.71472415 | 0.04319791 | 0.28126818 |
| FGR | 0.08438788 | 39.8226668 | 12 | 1.71889548 | 0.0428167 | 0.28126818 |
| RPS6KB2 | 0.07720901 | 36.4349562 | 15 | 1.75408652 | 0.03970784 | 0.28030873 |
| MAP2K7 | 0.06072389 | 28.6556207 | 25 | 1.76736763 | 0.03858334 | 0.28030873 |
| RPS6KA2 | 0.03428191 | 16.1776426 | 89 | 1.83009151 | 0.03361813 | 0.25168166 |
| MAP2K4 | 0.04424102 | 20.8773502 | 53 | 1.84956631 | 0.03218804 | 0.2476691 |
| GTF2F1 | 0.31674308 | 149.47116 | 1 | 1.89764767 | 0.02887125 | 0.22849536 |
| EGFR | 0.10343644 | 48.8116872 | 10 | 1.93244682 | 0.02665219 | 0.21713696 |
| KDR | 0.19019203 | 89.7516788 | 3 | 1.96476533 | 0.0247207 | 0.20750408 |
| CHUK | 0.0363439 | 17.1506956 | 91 | 1.96917996 | 0.02446621 | 0.20750408 |
| PRKCE | 0.04468187 | 21.0853898 | 59 | 1.97187669 | 0.02431184 | 0.20750408 |
| MAPK12 | 0.05224174 | 24.6528946 | 44 | 2.00532199 | 0.02246432 | 0.20750408 |
| RAF1 | 0.03640378 | 17.1789561 | 123 | 2.29338271 | 0.01091299 | 0.13143037 |
| SRC | 0.11269821 | 53.182323 | 12 | 2.31040179 | 0.01043296 | 0.13136045 |
| FYN | 0.20046469 | 94.5993487 | 4 | 2.39263418 | 0.00836396 | 0.11584078 |
| MAPK13 | 0.06186838 | 29.1957091 | 45 | 2.41747927 | 0.00781421 | 0.11392299 |
| ERBB2 | 0.2104204 | 99.2974528 | 4 | 2.51272951 | 0.00599006 | 0.0976027 |
| PKM | 0.43510203 | 205.324785 | 1 | 2.61152707 | 0.00450694 | 0.08917308 |
| MET | 0.14770988 | 69.7043382 | 10 | 2.77688338 | 0.00274414 | 0.06222414 |
| PTK6 | 0.48018633 | 226.600079 | 1 | 2.88345201 | 0.00196671 | 0.06053105 |
| RPS6KA1 | 0.11418962 | 53.8861177 | 19 | 2.94640178 | 0.00160747 | 0.05565874 |
| CSNK1G3 | 0.21067151 | 99.41595 | 10 | 3.97776361 | 3.48E-05 | 0.00321165 |
| PAK2 | 0.08948701 | 42.2289487 | 60 | 4.08179667 | 2.23E-05 | 0.00309471 |
| RPS6KB1 | 0.1147168 | 54.1348958 | 48 | 4.70515498 | 1.27E-06 | 0.00035134 |
