## Supplementary table 7 for "MAP4K inhibition by the CNS penetrant inhibitor famlasertib restrains medulloblastoma dissemination without developmental toxicity"

| Kinase.Gene | mS | Enrichment | m | z.score | p.value | FDR |
|---|---|---|---|---|---|---|
| ULK1 | -1.003243444 | -560.9791944 | 1 | -5.981372034 | 1.11E-09 | 2.36E-08 |
| CSNK2A1 | -0.048476958 | -27.10664623 | 269 | -4.573578615 | 2.40E-06 | 3.91E-05 |
| PKM | -0.514333179 | -287.5974063 | 1 | -3.061266791 | 0.001102013 | 0.012210305 |
| TRKB | -0.272173976 | -152.1903172 | 3 | -2.79713566 | 0.002577894 | 0.023493319 |
| IGF1R | -0.231866381 | -129.6516976 | 3 | -2.380154085 | 0.0086527 | 0.061456357 |
| TRKA | -0.271520877 | -151.8251263 | 2 | -2.278335199 | 0.011353307 | 0.07313642 |
| TRKC | -0.271520877 | -151.8251263 | 2 | -2.278335199 | 0.011353307 | 0.07313642 |
| ATM | -0.041256997 | -23.06949231 | 63 | -1.871076489 | 0.03066724 | 0.162013937 |
| SLK | -0.157319352 | -87.96756562 | 4 | -1.8578739 | 0.031593462 | 0.162013937 |
| TBK1 | -0.221447328 | -123.8257216 | 2 | -1.855381605 | 0.031770876 | 0.162013937 |
| PTK2 | -0.30676838 | -171.5343167 | 1 | -1.821548377 | 0.03426177 | 0.163767739 |
| CSNK2A2 | -0.040828897 | -22.83011341 | 61 | -1.821167258 | 0.034290718 | 0.163767739 |
| PRKDC | -0.029501949 | -16.49647423 | 113 | -1.759547699 | 0.039242263 | 0.181168447 |
| PRKD2 | -0.14364809 | -80.32306616 | 4 | -1.694565706 | 0.045078919 | 0.201400975 |
| ATR | -0.05579208 | -31.19701056 | 27 | -1.676002868 | 0.046868815 | 0.206073998 |
| CSNK1A1 | -0.033163832 | -18.54407288 | 76 | -1.633676806 | 0.051163366 | 0.216266197 |
| STK11 | -0.092769548 | -51.87353751 | 9 | -1.630204609 | 0.05152913 | 0.216266197 |
| FLT3 | -0.273480176 | -152.9206991 | 1 | -1.622728541 | 0.052323724 | 0.216323455 |
| CDC42BPA | -0.071055971 | -39.7320526 | 14 | -1.547973263 | 0.060814367 | 0.240651138 |
| TGFBR2 | -0.058860381 | -32.91269837 | 20 | -1.524429869 | 0.063700682 | 0.245070678 |
| EPHA2 | -0.08391049 | -46.91985709 | 9 | -1.47146761 | 0.07058235 | 0.260684147 |
| EPHA1 | -0.096396937 | -53.90184862 | 6 | -1.384125271 | 0.08316005 | 0.276027183 |
| EPHA3 | -0.096396937 | -53.90184862 | 6 | -1.384125271 | 0.08316005 | 0.276027183 |
| EPHA4 | -0.096396937 | -53.90184862 | 6 | -1.384125271 | 0.08316005 | 0.276027183 |
| MARK4 | -0.047795907 | -26.72582566 | 24 | -1.346182273 | 0.08912187 | 0.290432448 |
| MAP2K4 | -0.032089725 | -17.94346972 | 53 | -1.317556385 | 0.093826097 | 0.301965653 |
| CSF1R | -0.091434127 | -51.12681585 | 6 | -1.311519245 | 0.094841198 | 0.301965653 |
| CSNK1D | -0.023084234 | -12.90790901 | 102 | -1.284589944 | 0.099467807 | 0.308676889 |
| NUAK1 | -0.108835546 | -60.85709063 | 4 | -1.278717251 | 0.100498321 | 0.308676889 |
| KDR | -0.120339162 | -67.28951656 | 3 | -1.226406387 | 0.110022892 | 0.331264577 |
| NEK1 | -0.039346259 | -22.00107363 | 27 | -1.165607422 | 0.121886603 | 0.346448691 |
| PDGFRB | -0.073233621 | -40.94971935 | 7 | -1.128994112 | 0.129450159 | 0.35773093 |
| LIMK1 | -0.178874069 | -100.0202211 | 1 | -1.057676406 | 0.145101498 | 0.373991432 |
| LIMK2 | -0.178874069 | -100.0202211 | 1 | -1.057676406 | 0.145101498 | 0.373991432 |
| TESK1 | -0.178874069 | -100.0202211 | 1 | -1.057676406 | 0.145101498 | 0.373991432 |
| GRK6 | -0.103671077 | -57.96929728 | 3 | -1.053975254 | 0.145947123 | 0.373991432 |
| CDK3 | -0.045682584 | -25.54412798 | 16 | -1.048664394 | 0.147166304 | 0.373991432 |
| ABL1 | -0.047562904 | -26.59553843 | 13 | -0.985745653 | 0.16212897 | 0.399980023 |
| EPHB2 | -0.08395564 | -46.94510334 | 4 | -0.981517738 | 0.163168746 | 0.399980023 |
| EPHB1 | -0.07343598 | -41.06287155 | 5 | -0.956876748 | 0.169314734 | 0.4114051 |
| MAP2K5 | -0.039182469 | -21.90948798 | 18 | -0.947564049 | 0.171675718 | 0.413514555 |
| CDC42BPB | -0.06101505 | -34.11751551 | 7 | -0.935913459 | 0.174658873 | 0.417073344 |
| ACTR2 | -0.087439009 | -48.8928835 | 3 | -0.886054707 | 0.187794024 | 0.435272017 |
| TTK | -0.015125608 | -8.457719026 | 114 | -0.850525555 | 0.197516479 | 0.449001853 |
| PRKD1 | -0.039808633 | -22.25961693 | 14 | -0.849666268 | 0.197755329 | 0.449001853 |
| PRKCI | -0.023799683 | -13.30796325 | 40 | -0.831467179 | 0.202854881 | 0.449526416 |
| SRC | -0.041258979 | -23.07060055 | 12 | -0.816645735 | 0.207065455 | 0.451630954 |
| SRRM1 | -0.07853898 | -43.91629369 | 3 | -0.793984018 | 0.213602368 | 0.460829289 |
| EPHA7 | -0.06098029 | -34.09807853 | 5 | -0.790527002 | 0.21461003 | 0.460829289 |
| MARK1 | -0.027815231 | -15.55331958 | 25 | -0.777250473 | 0.218505505 | 0.465584806 |
| PLK1 | -0.012804163 | -7.159647139 | 131 | -0.753044093 | 0.225711709 | 0.472659249 |
| FLT1 | -0.064486065 | -36.05838757 | 4 | -0.748946608 | 0.226944694 | 0.472659249 |
| FGR | -0.037328761 | -20.87295796 | 12 | -0.735329622 | 0.231069386 | 0.475398059 |
| IRAK4 | -0.124561608 | -69.65056281 | 1 | -0.73328538 | 0.231692195 | 0.475398059 |
| MARK2 | -0.051497281 | -28.7955066 | 6 | -0.72724232 | 0.233538763 | 0.47566351 |
| TRPM7 | -0.045066648 | -25.19971818 | 7 | -0.683893148 | 0.24702132 | 0.490613893 |
| CSNK1E | -0.01955594 | -10.93500832 | 39 | -0.662719063 | 0.253755249 | 0.495001437 |
| WEE1 | -0.108455311 | -60.64447603 | 1 | -0.6370876 | 0.262033893 | 0.499446292 |
| NUAK2 | -0.108183139 | -60.49228682 | 1 | -0.635462003 | 0.26256357 | 0.499446292 |
| UHMK1 | -0.062832013 | -35.1334985 | 3 | -0.631495638 | 0.26385825 | 0.499446292 |
| MAP2K7 | -0.021908653 | -12.25056451 | 25 | -0.60085993 | 0.273966642 | 0.499446292 |
| PKD1 | -0.019644642 | -10.98460754 | 31 | -0.593800789 | 0.276322681 | 0.499446292 |
| ERBB4 | -0.071787645 | -40.14117931 | 2 | -0.591259088 | 0.277173419 | 0.499446292 |
| CDK14 | -0.067928841 | -37.98346953 | 2 | -0.558665141 | 0.288195137 | 0.499446292 |

| | | | | | | |
|---|---|---|---|---|---|---|
| BRSK2 | -0.090360508 | -50.52648516 | 1 | -0.529013107 | 0.298398179 | 0.499446292 |
| CSNK1G2 | -0.022022085 | -12.31399191 | 19 | -0.526770673 | 0.299176425 | 0.499446292 |
| LYN | -0.029567374 | -16.5330576 | 10 | -0.524669571 | 0.299906457 | 0.499446292 |
| CDK4 | -0.013667273 | -7.642268409 | 52 | -0.511619416 | 0.304458697 | 0.499446292 |
| NME1 | -0.083856374 | -46.88959735 | 1 | -0.490165986 | 0.312008224 | 0.499446292 |
| MST3 | -0.049082604 | -27.44530254 | 3 | -0.489258176 | 0.312329464 | 0.499446292 |
| OXSR1 | -0.059691973 | -33.37769624 | 2 | -0.48909122 | 0.312388559 | 0.499446292 |
| CDK9 | -0.026071517 | -14.578295 | 10 | -0.458642361 | 0.323245506 | 0.499446292 |
| BRAF | -0.040092683 | -22.41844789 | 4 | -0.457558804 | 0.323634723 | 0.499446292 |
| ADRBK1 | -0.012246591 | -6.847871954 | 51 | -0.446079091 | 0.327770058 | 0.499446292 |
| PRKG2 | -0.015997911 | -8.945480872 | 27 | -0.440992286 | 0.329609291 | 0.499446292 |
| CDKL5 | -0.075332703 | -42.12345398 | 1 | -0.439256815 | 0.330237731 | 0.499446292 |
| TNK2 | -0.075137921 | -42.01453859 | 1 | -0.438093444 | 0.330659273 | 0.499446292 |
| PRKACB | -0.016369642 | -9.153340096 | 24 | -0.426648382 | 0.334817726 | 0.499446292 |
| MAP3K1 | -0.072576402 | -40.58222535 | 1 | -0.422794308 | 0.336222671 | 0.499446292 |
| MELK | -0.042384503 | -23.69995487 | 3 | -0.4199664 | 0.337255 | 0.499446292 |
| TAF1 | -0.071887768 | -40.19716493 | 1 | -0.418681316 | 0.337724526 | 0.499446292 |
| GRK5 | -0.029667196 | -16.58887453 | 6 | -0.407867698 | 0.341685405 | 0.499446292 |
| LMTK2 | -0.068386898 | -38.23959895 | 1 | -0.397771735 | 0.345399227 | 0.499446292 |
| STK39 | -0.04727639 | -26.43532967 | 2 | -0.384221173 | 0.350407263 | 0.499446292 |
| PLK4 | -0.012898762 | -7.212543616 | 30 | -0.363461992 | 0.3581299 | 0.499446292 |
| PDPK1 | -0.03658492 | -20.45702773 | 3 | -0.359969786 | 0.359434864 | 0.499446292 |
| WNK3 | -0.060772757 | -33.98203344 | 1 | -0.352294897 | 0.362308557 | 0.499446292 |
| CLK1 | -0.00616131 | -3.445192364 | 156 | -0.326215246 | 0.372130754 | 0.499446292 |
| PBK | -0.039757696 | -22.23113483 | 2 | -0.320713418 | 0.374213789 | 0.499446292 |
| ADRBK2 | -0.023632944 | -13.21472888 | 5 | -0.291741195 | 0.385242255 | 0.499446292 |
| TEC | -0.050431482 | -28.19954849 | 1 | -0.290529756 | 0.385705495 | 0.499446292 |
| AURKA | -0.004870541 | -2.723439036 | 226 | -0.276744545 | 0.390988133 | 0.499446292 |
| CSNK1G1 | -0.012437338 | -6.954531002 | 18 | -0.269843982 | 0.393640142 | 0.499446292 |
| GRK7 | -0.021385868 | -11.95824089 | 5 | -0.261730762 | 0.396764511 | 0.499446292 |
| PDHK3 | -0.031837939 | -17.80267938 | 2 | -0.253818026 | 0.399818073 | 0.499446292 |
| PDK3 | -0.04380984 | -24.49695425 | 1 | -0.250980798 | 0.400914478 | 0.499446292 |
| PDK4 | -0.04380984 | -24.49695425 | 1 | -0.250980798 | 0.400914478 | 0.499446292 |
| BLK | -0.015251803 | -8.528283214 | 9 | -0.241238227 | 0.404685242 | 0.499446292 |
| MAPK6 | -0.039858905 | -22.28772742 | 1 | -0.227383122 | 0.410062922 | 0.499446292 |
| PKD2 | -0.022929197 | -12.82121737 | 3 | -0.218701497 | 0.413441292 | 0.499446292 |
| STK3 | -0.037463167 | -20.94811321 | 1 | -0.213074142 | 0.415634567 | 0.499446292 |
| STK4 | -0.037463167 | -20.94811321 | 1 | -0.213074142 | 0.415634567 | 0.499446292 |
| CDK16 | -0.021829707 | -12.20642061 | 3 | -0.20732729 | 0.417877132 | 0.499446292 |
| MOS | -0.036199119 | -20.24130109 | 1 | -0.205524386 | 0.418581224 | 0.499446292 |
| EPHA5 | -0.020780794 | -11.61990475 | 3 | -0.196476301 | 0.42211869 | 0.499446292 |
| EPHA6 | -0.020780794 | -11.61990475 | 3 | -0.196476301 | 0.42211869 | 0.499446292 |
| BCKDK | -0.012455405 | -6.964633454 | 8 | -0.1802012 | 0.428497308 | 0.499446292 |
| PTK6 | -0.031357339 | -17.5339444 | 1 | -0.176605977 | 0.42990895 | 0.499446292 |
| MAP3K7 | -0.022426524 | -12.54014012 | 2 | -0.174323126 | 0.430805763 | 0.499446292 |
| HCK | -0.010981941 | -6.140723435 | 10 | -0.173641354 | 0.431073665 | 0.499446292 |
| AAK1 | -0.029752238 | -16.63642719 | 1 | -0.16701922 | 0.433677463 | 0.499446292 |
| CSNK2B | -0.020375623 | -11.39334717 | 2 | -0.156999893 | 0.437622466 | 0.499446292 |
| DYRK3 | -0.027517073 | -15.38659987 | 1 | -0.153669292 | 0.438935249 | 0.499446292 |
| GSG2 | -0.019231261 | -10.75345934 | 2 | -0.147333871 | 0.441434249 | 0.499446292 |
| CDC7 | -0.013757663 | -7.692811503 | 4 | -0.142977439 | 0.443154 | 0.499446292 |
| LATS1 | -0.022316795 | -12.47878345 | 1 | -0.122609688 | 0.451208091 | 0.499446292 |
| BTK | -0.014366578 | -8.033295808 | 2 | -0.106243607 | 0.457694537 | 0.499446292 |
| ILK | -0.019339605 | -10.8140413 | 1 | -0.104827881 | 0.458256193 | 0.499446292 |
| TLK2 | -0.008509929 | -4.758459319 | 6 | -0.098336422 | 0.460832579 | 0.499446292 |
| MAPKAPK3 | -0.013249215 | -7.408504607 | 2 | -0.096805632 | 0.461440376 | 0.499446292 |
| CDK2 | -0.002676064 | -1.496362736 | 312 | -0.093649484 | 0.4626938 | 0.499446292 |
| PRKACG | -0.004528603 | -2.532238823 | 26 | -0.083453021 | 0.466745666 | 0.499446292 |
| ITK | -0.008723892 | -4.878099922 | 4 | -0.082847215 | 0.466986513 | 0.499446292 |
| MOK | -0.008525823 | -4.767346417 | 4 | -0.080481206 | 0.467927272 | 0.499446292 |
| MAPK15 | -0.013661786 | -7.639200386 | 1 | -0.070916077 | 0.471732274 | 0.499446292 |
| FYN | -0.007438216 | -4.159194306 | 4 | -0.067489352 | 0.473096069 | 0.499446292 |
| CLK3 | -0.006279377 | -3.511211433 | 3 | -0.046459318 | 0.481472079 | 0.499446292 |
| HIPK2 | -0.002432563 | -1.360205212 | 118 | -0.041794576 | 0.48333123 | 0.499446292 |
| BMX | -0.00595545 | -3.330082887 | 2 | -0.035197782 | 0.485961016 | 0.499446292 |
| MAP3K10 | -0.007220374 | -4.037384559 | 1 | -0.032443575 | 0.487059156 | 0.499446292 |

| | | | | | | |
|---|---|---|---|---|---|---|
| MAP3K5 | -0.004627561 | -2.587572794 | 3 | -0.029371302 | 0.48828423 | 0.499446292 |
| MAP3K6 | -0.004627561 | -2.587572794 | 3 | -0.029371302 | 0.48828423 | 0.499446292 |
| MAP4K4 | -0.003364929 | -1.88155245 | 5 | -0.021055331 | 0.491600759 | 0.499446292 |
| KIT | -0.003081205 | -1.722904036 | 2 | -0.010920049 | 0.495643617 | 0.499446292 |
| PLK3 | -0.002420968 | -1.353721771 | 3 | -0.006544121 | 0.497389292 | 0.499446292 |
| NEK3 | -0.002256966 | -1.262017685 | 2 | -0.003957988 | 0.498420995 | 0.499446292 |
| WNK2 | -0.002322385 | -1.298597737 | 1 | -0.003189447 | 0.498727597 | 0.499446292 |
| WNK1 | -0.001952698 | -1.091881215 | 2 | -0.00138794 | 0.499446292 | 0.499446292 |
| BUB1 | -0.001370587 | -0.766385344 | 2 | 0.003528938 | 0.49859216 | 0.499446292 |
| RIPK1 | 0.000791 | 0.442299968 | 1 | 0.015405809 | 0.493854214 | 0.499446292 |
| MAP4K6 | 0.000945093 | 0.528463328 | 4 | 0.032652312 | 0.486975926 | 0.499446292 |
| MAK | 0.000122017 | 0.068227706 | 10 | 0.036082179 | 0.485608416 | 0.499446292 |
| AURKC | 0.000135729 | 0.075894769 | 13 | 0.041435291 | 0.483474439 | 0.499446292 |
| TLK1 | 0.001187762 | 0.664155525 | 6 | 0.043541006 | 0.482635139 | 0.499446292 |
| DYRK1B | 0.007683989 | 4.296622106 | 1 | 0.056575436 | 0.477441701 | 0.499446292 |
| ACVR2B | 0.004720678 | 2.639640615 | 3 | 0.067336115 | 0.473157063 | 0.499446292 |
| SRPK1 | 0.010312635 | 5.76647073 | 2 | 0.102212999 | 0.459293805 | 0.499446292 |
| CAMK1G | 0.011647591 | 6.512931886 | 2 | 0.113488897 | 0.454821483 | 0.499446292 |
| NEK4 | 0.005636379 | 3.151669159 | 8 | 0.125428623 | 0.450092114 | 0.499446292 |
| CAMK1 | 0.004200228 | 2.348622893 | 15 | 0.138529031 | 0.444911164 | 0.499446292 |
| BCR | 0.017140895 | 9.584598361 | 2 | 0.159888897 | 0.436484298 | 0.499446292 |
| MKNK1 | 0.0172312 | 9.635093528 | 2 | 0.160651667 | 0.43618388 | 0.499446292 |
| MKNK2 | 0.0172312 | 9.635093528 | 2 | 0.160651667 | 0.43618388 | 0.499446292 |
| IRR | 0.027446816 | 15.34731457 | 1 | 0.174612506 | 0.430692061 | 0.499446292 |
| TXK | 0.027446816 | 15.34731457 | 1 | 0.174612506 | 0.430692061 | 0.499446292 |
| TYK2 | 0.027446816 | 15.34731457 | 1 | 0.174612506 | 0.430692061 | 0.499446292 |
| ACVR1B | 0.029470376 | 16.47881966 | 1 | 0.186698585 | 0.425948483 | 0.499446292 |
| CDK19 | 0.029901022 | 16.71962208 | 1 | 0.189270696 | 0.424940331 | 0.499446292 |
| PNCK | 0.032358573 | 18.09379999 | 1 | 0.203948865 | 0.419196729 | 0.499446292 |
| CDK6 | 0.004978585 | 2.783853465 | 29 | 0.217651785 | 0.413850217 | 0.499446292 |
| STK25 | 0.035702978 | 19.96387612 | 1 | 0.223923931 | 0.411408247 | 0.499446292 |
| MST1 | 0.039125703 | 21.87774611 | 1 | 0.244366777 | 0.403473385 | 0.499446292 |
| PDHK4 | 0.010824355 | 6.052606611 | 12 | 0.260957154 | 0.397062774 | 0.499446292 |
| PAK7 | 0.043049167 | 24.0716121 | 1 | 0.267800377 | 0.39442649 | 0.499446292 |
| NLK | 0.007520852 | 4.205401349 | 24 | 0.272388498 | 0.392661658 | 0.499446292 |
| PKD3 | 0.020268287 | 11.33332841 | 5 | 0.294573871 | 0.384159712 | 0.499446292 |
| PRKG1 | 0.005165441 | 2.888336536 | 52 | 0.299498356 | 0.382279913 | 0.499446292 |
| ICK | 0.013922404 | 7.78492905 | 11 | 0.311217153 | 0.377817772 | 0.499446292 |
| CAMK1D | 0.01950095 | 10.90425987 | 6 | 0.311463349 | 0.377724201 | 0.499446292 |
| CLK4 | 0.036266488 | 20.27897151 | 2 | 0.32143603 | 0.373939991 | 0.499446292 |
| SMG1 | 0.036538976 | 20.4313377 | 2 | 0.323737644 | 0.373068334 | 0.499446292 |
| PRKAA1 | 0.008503258 | 4.754729272 | 38 | 0.378918335 | 0.352374254 | 0.499446292 |
| CHEK2 | 0.018338141 | 10.25405727 | 10 | 0.380135164 | 0.351922542 | 0.499446292 |
| IRAK1 | 0.061929811 | 34.6290182 | 1 | 0.380568448 | 0.351761749 | 0.499446292 |
| NEK6 | 0.061929811 | 34.6290182 | 1 | 0.380568448 | 0.351761749 | 0.499446292 |
| PRKCG | 0.006064466 | 3.391040634 | 68 | 0.386768269 | 0.349463885 | 0.499446292 |
| MTOR | 0.011478759 | 6.418526938 | 24 | 0.388197043 | 0.348935111 | 0.499446292 |
| DYRK1A | 0.027558144 | 15.40956558 | 5 | 0.391932259 | 0.347554133 | 0.499446292 |
| CDK7 | 0.004761265 | 2.662335191 | 111 | 0.412143546 | 0.340117106 | 0.499446292 |
| CDK20 | 0.067766485 | 37.89268536 | 1 | 0.415429042 | 0.338913925 | 0.499446292 |
| PLK2 | 0.027817177 | 15.55440779 | 6 | 0.433129933 | 0.332460189 | 0.499446292 |
| MARK3 | 0.013420673 | 7.504378103 | 23 | 0.435647677 | 0.331546187 | 0.499446292 |
| PRKCE | 0.007847001 | 4.387772861 | 59 | 0.442042685 | 0.329229159 | 0.499446292 |
| PRKCA | 0.004908674 | 2.74476128 | 131 | 0.457813621 | 0.323543175 | 0.499446292 |
| EEF2K | 0.055023034 | 30.76698629 | 2 | 0.479865954 | 0.315661356 | 0.499446292 |
| PRKCQ | 0.039002321 | 21.80875542 | 4 | 0.487259717 | 0.313037146 | 0.499446292 |
| IKBKB | 0.007158031 | 4.002524396 | 86 | 0.495527048 | 0.310114067 | 0.499446292 |
| PRKCB | 0.003624384 | 2.026630983 | 243 | 0.50395474 | 0.307146591 | 0.499446292 |
| HIPK1 | 0.05835804 | 32.63180683 | 2 | 0.508035572 | 0.305714198 | 0.499446292 |
| GRK1 | 0.026880921 | 15.03088551 | 9 | 0.513697783 | 0.303731651 | 0.499446292 |
| GTF2F1 | 0.095703135 | 53.51389824 | 1 | 0.582285752 | 0.28018711 | 0.499446292 |
| EIF2AK2 | 0.012817224 | 7.166950206 | 45 | 0.585187565 | 0.279210803 | 0.499446292 |
| PAK1 | 0.014056894 | 7.860131266 | 50 | 0.66919715 | 0.251684862 | 0.494444729 |
| CLK2 | 0.025861736 | 14.46099267 | 17 | 0.680911642 | 0.247963701 | 0.490613893 |
| MAP2K6 | 0.017149649 | 9.589493531 | 38 | 0.697261863 | 0.242819465 | 0.487398491 |
| SGK3 | 0.057605889 | 32.21122988 | 4 | 0.70948608 | 0.239011443 | 0.483256714 |

| | | | | | | |
|---|---|---|---|---|---|---|
| GRK4 | 0.072020167 | 40.27119789 | 3 | 0.763548496 | 0.222568176 | 0.470621257 |
| DYRK2 | 0.046950853 | 26.25330037 | 8 | 0.823366173 | 0.205149897 | 0.451004138 |
| BRSK1 | 0.096815964 | 54.13615391 | 2 | 0.832876082 | 0.20245731 | 0.449526416 |
| CHUK | 0.012972198 | 7.253606425 | 91 | 0.840994927 | 0.200175388 | 0.449526416 |
| PIM2 | 0.058580285 | 32.75607877 | 6 | 0.883194877 | 0.188565495 | 0.435272017 |
| GSK3B | 0.009543164 | 5.336208671 | 172 | 0.887610885 | 0.187375048 | 0.435272017 |
| GSK3A | 0.010117994 | 5.657634168 | 161 | 0.902322506 | 0.183442787 | 0.434304717 |
| CHEK1 | 0.024738742 | 13.83305297 | 39 | 0.989445267 | 0.161222668 | 0.399980023 |
| MAPKAPK5 | 0.067712295 | 37.8623839 | 6 | 1.01679637 | 0.154625155 | 0.389374253 |
| SGK2 | 0.123978725 | 69.32463498 | 2 | 1.06231034 | 0.144047411 | 0.373991432 |
| PRKCZ | 0.029728964 | 16.62341292 | 32 | 1.064863474 | 0.143468858 | 0.373991432 |
| PAK3 | 0.05338009 | 29.84830847 | 11 | 1.092840099 | 0.137232006 | 0.372679075 |
| PAK2 | 0.022514037 | 12.58907419 | 60 | 1.124331606 | 0.130436188 | 0.35773093 |
| RPS6KB2 | 0.04808202 | 26.88581008 | 15 | 1.153606831 | 0.124330698 | 0.347874782 |
| RPS6KA3 | 0.034985976 | 19.56295317 | 28 | 1.1622336 | 0.122570295 | 0.346448691 |
| SGK1 | 0.033173222 | 18.54932353 | 31 | 1.162629947 | 0.122489837 | 0.346448691 |
| PIM3 | 0.086276599 | 48.24290385 | 5 | 1.176136104 | 0.119770246 | 0.346448691 |
| NEK11 | 0.098873285 | 55.28653684 | 4 | 1.202440093 | 0.114596532 | 0.337694035 |
| PAK6 | 0.100023893 | 55.92991736 | 4 | 1.216184526 | 0.111957318 | 0.333464271 |
| CDK5 | 0.017059012 | 9.538812477 | 128 | 1.273578111 | 0.101406487 | 0.308676889 |
| PAK5 | 0.086347943 | 48.28279696 | 6 | 1.289436314 | 0.098623222 | 0.308676889 |
| CAMK2D | 0.041894507 | 23.42596576 | 28 | 1.38057398 | 0.083704994 | 0.276027183 |
| AURKB | 0.025279711 | 14.13554385 | 73 | 1.381301187 | 0.083593187 | 0.276027183 |
| CAMK2G | 0.056222771 | 31.43783764 | 16 | 1.385928455 | 0.082884376 | 0.276027183 |
| NEK2 | 0.013018136 | 7.279293207 | 249 | 1.395474359 | 0.081436421 | 0.276027183 |
| CAMK4 | 0.054040398 | 30.21753001 | 18 | 1.414697919 | 0.078578542 | 0.276027183 |
| VRK1 | 0.137592045 | 76.93673457 | 3 | 1.441888763 | 0.074666878 | 0.272141122 |
| RPS6KA2 | 0.024893255 | 13.91945121 | 89 | 1.503407721 | 0.066366968 | 0.248427706 |
| MAPK10 | 0.041675275 | 23.30337884 | 34 | 1.51368337 | 0.065053082 | 0.246845258 |
| PRKAA2 | 0.027329056 | 15.28146725 | 77 | 1.52604671 | 0.063499116 | 0.245070678 |
| PRKACA | 0.032171973 | 17.98945942 | 60 | 1.571148197 | 0.058074115 | 0.233138113 |
| PDK2 | 0.065004583 | 36.34832502 | 16 | 1.595732335 | 0.055274283 | 0.225161418 |
| MAPK7 | 0.0354828 | 19.84076039 | 54 | 1.635834506 | 0.050937114 | 0.216266197 |
| MAPK11 | 0.036627919 | 20.48107131 | 55 | 1.701634201 | 0.044411981 | 0.201400975 |
| PIM1 | 0.068833558 | 38.48935588 | 18 | 1.789555727 | 0.036762681 | 0.17259767 |
| PRKCH | 0.054511519 | 30.48096481 | 30 | 1.841779292 | 0.03275372 | 0.162013937 |
| DAPK1 | 0.067248011 | 37.60277235 | 20 | 1.844006374 | 0.032591106 | 0.162013937 |
| PAK4 | 0.051450458 | 28.76932462 | 34 | 1.854117977 | 0.031861142 | 0.162013937 |
| DMPK | 0.087184748 | 48.75070929 | 13 | 1.916019291 | 0.02768132 | 0.153354514 |
| CAMK2A | 0.057098423 | 31.9274723 | 31 | 1.958250131 | 0.025100335 | 0.141893728 |
| ROCK1 | 0.038651944 | 21.61283612 | 75 | 2.091773185 | 0.018229405 | 0.105198858 |
| TRPM6 | 0.102078566 | 57.07882 | 12 | 2.149004533 | 0.01581702 | 0.09321946 |
| CDK1 | 0.01235273 | 6.90722108 | 654 | 2.159939964 | 0.015388659 | 0.092666488 |
| PRKCD | 0.049419882 | 27.63389651 | 50 | 2.162690525 | 0.015282495 | 0.092666488 |
| CSNK1G3 | 0.112914636 | 63.13797677 | 10 | 2.166427602 | 0.015139263 | 0.092666488 |
| CAMK2B | 0.074396145 | 41.59976275 | 26 | 2.320185876 | 0.010165412 | 0.068678516 |
| MAPK13 | 0.057179206 | 31.9726434 | 45 | 2.362593186 | 0.009073789 | 0.06283599 |
| MAPK9 | 0.059865647 | 33.47480903 | 42 | 2.386467029 | 0.008505566 | 0.061456357 |
| ARAF | 0.070762353 | 39.56787135 | 31 | 2.412637043 | 0.007918791 | 0.059283923 |
| VRK2 | 0.425972921 | 238.1893921 | 1 | 2.554881958 | 0.005311191 | 0.040866662 |
| VRK3 | 0.425972921 | 238.1893921 | 1 | 2.554881958 | 0.005311191 | 0.040866662 |
| MAPK12 | 0.065205397 | 36.46061295 | 44 | 2.654178591 | 0.003975087 | 0.032385267 |
| RAF1 | 0.039115916 | 21.87227379 | 123 | 2.709510393 | 0.00336913 | 0.028280271 |
| AKT3 | 0.084837554 | 47.43823957 | 28 | 2.73776578 | 0.003092906 | 0.026772965 |
| RPS6KB1 | 0.065653963 | 36.71143562 | 48 | 2.790760766 | 0.002629216 | 0.023493319 |
| ROCK2 | 0.079060579 | 44.20795403 | 34 | 2.815679608 | 0.002433709 | 0.023246118 |
| PDK1 | 0.056156461 | 31.40075988 | 71 | 2.916173065 | 0.001771769 | 0.017527862 |
| MAPK8 | 0.044459743 | 24.86035755 | 112 | 2.923293769 | 0.001731748 | 0.017527862 |
| PRKCT | 0.068672637 | 38.39937426 | 49 | 2.94588839 | 0.001610143 | 0.017154214 |
| MAPKAPK2 | 0.137964389 | 77.14493667 | 15 | 3.232774394 | 0.000612972 | 0.007074715 |
| AKT2 | 0.080789926 | 45.17494519 | 45 | 3.308579464 | 0.000468853 | 0.005646619 |
| MAPK14 | 0.081442248 | 45.53970101 | 46 | 3.371564164 | 0.000373713 | 0.004705388 |
| MAP2K3 | 0.115713348 | 64.70292044 | 25 | 3.509001895 | 0.000224896 | 0.002966483 |
| STK10 | 0.488592487 | 273.2040973 | 2 | 4.142074083 | 1.72E-05 | 0.000238344 |
| YES1 | 0.203288911 | 113.6721604 | 12 | 4.243044064 | 1.10E-05 | 0.000160739 |
| RPS6KA1 | 0.166304181 | 92.9915725 | 19 | 4.376174593 | 6.04E-06 | 9.29E-05 |

| | | | | | | |
|---|---|---|---|---|---|---|
| MAPK3 | 0.047977453 | 26.82734007 | 308 | 5.21646116 | 9.12E-08 | 1.58E-06 |
| AKT1 | 0.06923572 | 38.71423073 | 168 | 5.498316301 | 1.92E-08 | 3.54E-07 |
| MAP3K8 | 0.549712551 | 307.3803331 | 3 | 5.705270301 | 5.81E-09 | 1.15E-07 |
| MAPK1 | 0.065408671 | 36.57427687 | 272 | 6.6191772 | 1.81E-11 | 4.17E-10 |
| EGFR | 0.406950436 | 227.5526734 | 10 | 7.719963199 | 5.82E-15 | 1.47E-13 |
| MET | 0.419702893 | 234.6834085 | 10 | 7.960822391 | 8.54E-16 | 2.37E-14 |
| DAPK3 | 1.011495175 | 565.593278 | 2 | 8.55884852 | 5.70E-18 | 1.75E-16 |
| MAP2K1 | 0.176994107 | 98.96901112 | 69 | 8.869902906 | 3.66E-19 | 1.27E-17 |
| DAPK2 | 1.908205633 | 1067.002895 | 1 | 11.4077857 | 1.91E-30 | 7.56E-29 |
| INSR | 0.867839605 | 485.2660292 | 5 | 11.6141614 | 1.75E-31 | 8.06E-30 |
| MAP2K2 | 0.377140185 | 210.8838075 | 29 | 12.18781047 | 1.80E-34 | 1.00E-32 |
| LCK | 0.971279487 | 543.1060502 | 5 | 12.9956343 | 6.48E-39 | 4.48E-37 |
| ERBB2 | 1.21272061 | 678.1116135 | 4 | 14.50775048 | 5.41E-48 | 5.00E-46 |
| RET | 1.673607902 | 935.8239201 | 3 | 17.33195381 | 1.35E-67 | 1.87E-65 |
| JAK2 | 2.427396481 | 1357.316542 | 2 | 20.51846708 | 7.36E-94 | 2.04E-91 |
